# Impaired capillary-venous drainage contributes to gliosis and demyelination in white matter during aging

**DOI:** 10.1101/2024.02.11.579849

**Authors:** Stefan Stamenkovic, Franca Schmid, Gokce Gurler, Farzaneh Abolmaali, Nicolas A. Weitermann, Kevin T. Takasaki, Stephanie K. Bonney, Maria J. Sosa, Hannah C. Bennett, Yongsoo Kim, Jack Waters, Andy Y. Shih

## Abstract

The progressive loss of cerebral white matter during aging contributes to cognitive decline, but whether reduced blood flow is a cause or consequence remains debated. Using deep multi-photon imaging in mice, we examined microvascular networks perfusing myelinated tissues in cortical layer 6 and corpus callosum. We identified sparse, wide-reaching venules, termed principal cortical venules, that exclusively drain deep tissues and resemble vasculature at the human cortex and U-fiber interface. Aging involved selective constriction and rarefaction of capillaries in deep branches of principal cortical venules. This resulted in mild hypoperfusion that was associated with microgliosis, astrogliosis and demyelination in deep tissues, but not upper cortex. Inducing a comparable hypoperfusion in adult mice using carotid artery stenosis triggered a similar tissue pathology specific to layer 6 and corpus callosum. Thus, impaired capillary-venous drainage is a contributor to hypoperfusion and a potential therapeutic target for preserving blood flow to white matter during aging.

## INTRODUCTION

The gradual loss of cerebral white matter during aging contributes to cognitive decline.^1^ At the microstructural level, this process involves axon degeneration, demyelination, microgliosis and astrogliosis, which impairs neurotransmission and produces an inflammatory environment detrimental to white matter maintenance.^2,3^ Since oligodendrocytes are uniquely vulnerable to ischemia and myelin repair is energy-intensive^4,5^, age-related deficits in cerebral blood flow^6,7^ and vascular structure loss^8–10^ are often considered contributors to white matter deterioration. However, a direct causal link between cerebral hypoperfusion and white matter loss during aging remains unestablished.^11^ An alternative possibility is that vascular changes result as a consequence of white matter loss due to mechanisms unrelated to blood flow. Understanding this relationship is crucial for designing effective therapeutic strategies to preserve white matter integrity. Critical questions remain regarding the specific microvascular changes driving age-related hypoperfusion, and whether its severity is sufficient to cause white matter deterioration. Addressing these gaps is challenging in humans, as clinical studies are typically correlative and non-invasive imaging cannot resolve microvascular changes. This underscores the need for complementary preclinical studies capable of resolving white matter microvasculature *in vivo* and manipulating blood flow to establish causal relationships.

In rodents, the corpus callosum (CC) is the largest white matter tract and a primary focus for white matter research.^12,13^ It consists of neuronal projections that serve local crosstalk between neighboring cortical regions and long-range projections for cortical-subcortical communication. However, the anatomy and vascular architecture of the rodent CC more closely resembles that of the human superficial white matter (or U-fibers) that lie adjacent to the cortical gray matter, as opposed to the human CC. Superficial white matter facilitates both short-range communication between neighboring gyri and longer-range communication with deeper subcortical white matter, which are essential for complex cognitive processes.^14^ Like deep subcortical white matter, superficial white matter undergoes age- and disease-related degeneration, and these changes are increasingly recognized as an early biomarker of white matter degeneration and a contributor to cognitive decline.^15^ Further, vascular contributions to loss of superficial white matter has been implicated in chronic conditions, such as Alzheimer’s disease^16^, cerebral small vessel disease^17^, and multiple sclerosis^18^

To better understand the microvascular architecture of the human cerebral cortex and superficial white matter, Duvernoy and colleagues used vascular casts in post-mortem tissues to reveal large, horizontally projecting branches of principal cortical veins/venules (PCVs) within deep cortical layers and the U fiber tracts.^19^ Their findings suggested that the quality of PCVs drainage can significantly influence blood flow in these regions. However, their structure, function, and perfusion territories remain largely unexamined. To address this knowledge gap, we leveraged recent advances in deep multi-photon imaging to extend *in vivo* imaging to the deep cortex and CC of adult and aged mice.^20,21^ This allowed us to ask whether murine white matter is drained by an equivalent of the human PCV. Our *in vivo* studies of vascular physiology were complemented by light-sheet imaging and *in silico* modeling to provide a comprehensive view of how age-related vascular changes affect cerebral perfusion. These studies revealed specific microvascular alterations in PCV branching networks that contribute to age-related hypoperfusion, offering potential vascular targets to improve cerebral blood flow and mitigate age-related white matter decline.

## RESULTS

### The murine cerebrovasculature contains an equivalent to human PCVs

In the human brain, PCVs exhibit a conical structure, with a large trunks spanning all cortical layers and deeper branches extending over long distances along the interface between cortical gray matter and underlying U-fiber white matter (**Fig. 1a, b**).^19^ Individual PCVs are surrounded by rings of penetrating arterioles, suggesting that each serves as the sole output for multiple sources of blood input to relatively large tissue regions (**Fig. 1c**). This implicates PCVs as bottlenecks in perfusion of the deep cortex and superficial white matter in humans.

**Figure 1.**
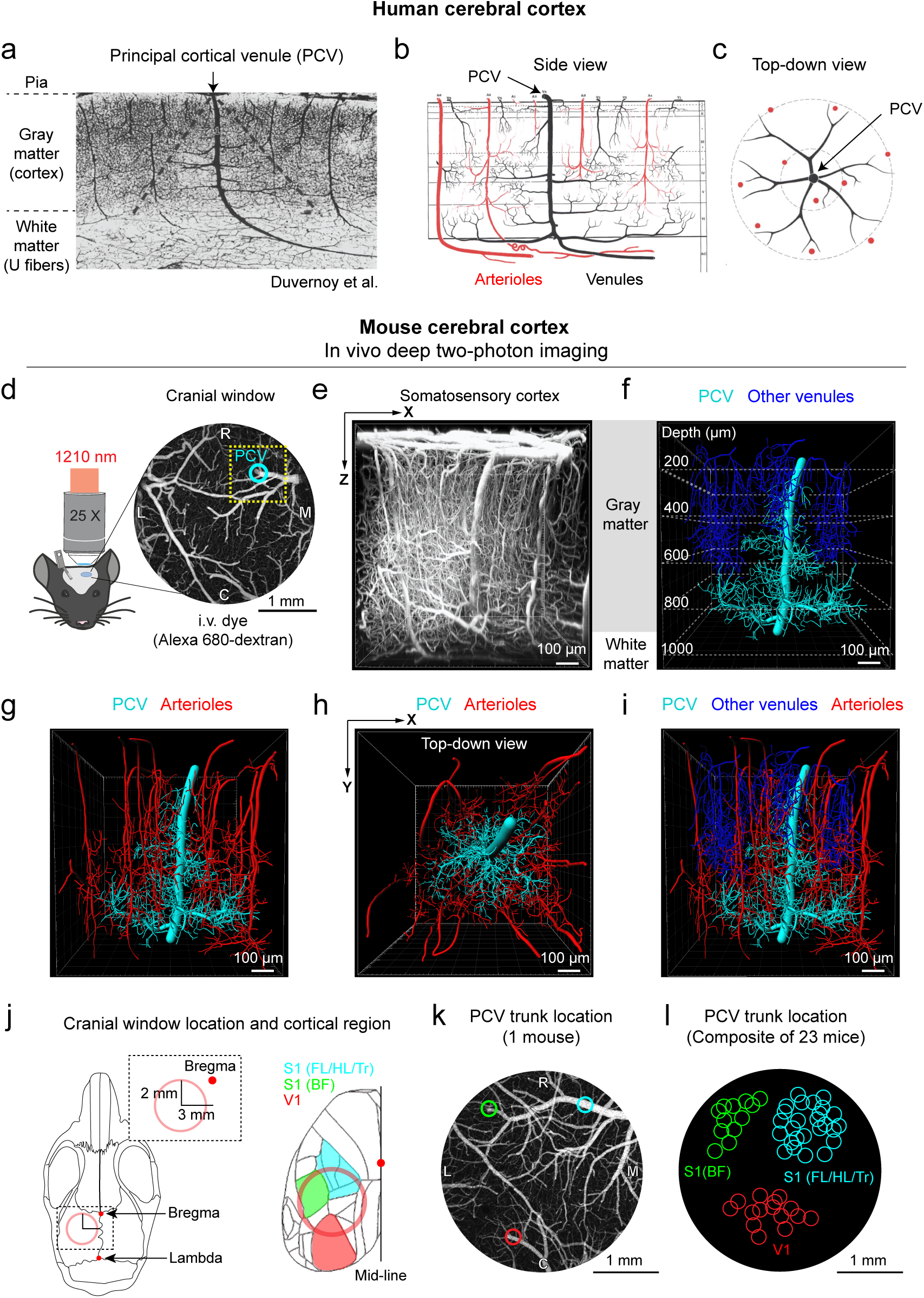
Principal cortical venules in human and murine cortex. **a.** PCV visualized in India ink-filled human brain sections from Duvernoy et al.^19^ **b, c.** Schematics from Duvernoy et al. show PCV structure from side (b) and above (c). **d.** Schematic summary of *in vivo* multi-photon imaging set up. Image on right shows low magnification view of chronic window with PCV location marked with a cyan circle. **e.** Side view of the microvasculature in adult mouse somatosensory cortex showing a 900 x 900 x 1000 µm (x, y, z) volume collected by deep two-photon imaging in panel d (yellow dotted square). **f.** Imaris reconstruction of a PCV in panel (e) together with branching networks (cyan). All other ascending venules (blue) are also depicted. Dotted lines mark the depth below the pial surface. **g, h.** The same PCV alongside all penetrating arterioles (red), viewed from side (g) and top (h). **i.** Composite of all segmented vessel types. **j.** Schematic of the mouse skull with calvaria landmarks and bones labeled. The red circle marks the location of the implanted cranial windows and cortical sensory regions accessible within the window are shown on the right. S1 (FL/HL/Tr) – Somatosensory cortex forelimb, hindlimb, and trunk regions, S1 (BF) – Somatosensory cortex barrel field region, V1 – Primary visual cortex. **k.** PVC trunk locations in an example cranial window. **l.** The position of PCV trunk locations identified within cranial windows for n=23 mice, all verified by in vivo imaging. Cyan colored circles mark the locations of PCVs analyzed in this study.

To examine whether there is a structural correlate to the human PCV in the murine brain, we performed *in vivo* deep two-photon imaging.^20^ This approach used a far-red intravenous dye, Alexa 680-dextran, and longer wavelength excitation to achieve deeper tissue penetration than conventional two-photon microscopy (**Fig. 1d**). Imaging was performed on Thy1-YFP mice to provide reference to cortical layers with neuronal labeling. In a sub-regions within large cranial windows, we collected high-resolution data sets, averaging 900 µm (x) by 900 µm (y) by 1000 µm (z) in size, that centered around PCVs in the forelimb, hindlimb, and trunk region of the primary sensory cortex (S1-FL/HL/Tr)(**Fig. 1e, Supplementary Movie 1**). Segmentation of penetrating vessels within these volumes confirmed a vascular structure similar to human PCVs (**Fig. 1f, cyan vessel, Supplementary Movie 2**). Murine PCVs were also conical-shaped and extended progressively longer reaching branches with increasing tissue depth. The deeper branches were connected to elaborate capillary networks spanning the gray-white matter interface.

Although only one PCV was in each imaging volume collected, the same volume contained 23 ± 3 other ascending venules (**Fig. 1f, dark blue vessels**; n = 9 mice; mean ± SD). However, these other venules terminated at or before cortical layer 5. As such, PCVs represented <4% of all ascending cortical venules and were the only venules that reached the deep cortex and CC. The same tissue volume also contained 15 ± 2 penetrating arterioles, oriented in a ring around the trunk of the PCV, again similar to the human cortex (**Fig. 1g-i**). Approximately 50% of these penetrating arterioles had branches that reached deep cortical layers, ramifying upon reaching the CC. Thus, deeper tissues are served by multiple arteriolar inputs, but rare PCV outputs in mice and humans.

### PCVs are a rare penetrating vessel type draining predominantly from deeper tissues

To gain a broader view of PCV drainage territories, we surveyed their positions within the cranial imaging window (**Fig. 1j-l**). The main trunk of murine PCVs ascended to the brain surface either at the end of large diameter pial venules (**Fig. 1k, red and green circles**), or at locations hidden beneath large pial venules as they coursed along the surface (**Fig. 1k, cyan circle**). Individual PCVs tended to be centered within major representations of primary sensory cortex (S1): FL/HL/Tr, barrel field (BF), and visual (V1) regions (**Fig. 1l**), with one PCV per region, suggesting they may be strategically placed to serve major hubs in cortical and white matter function.

To better understand the proportion of penetrating vessel types, we examined light-sheet imaging data collected from optically-cleared *post-mortem* mouse brains with all vasculature labeled by fluorescent lectin and arterioles with α-SMA and SM22 (**Extended Fig. 1a,b**). The number and penetration depth of all penetrating arterioles, PCVs and other ascending venules was examined in 2 mm (x) by 2 mm (y) by 1.2 mm (z) regions of interest in the primary somatosensory cortex, which were ∼6-times larger in volume than sub-regions examined *in vivo* (**Extended Fig. 1c**). This volume contained on average 55 ± 10 descending arterioles (n = 4 mice; mean ± SD), with a homogenous distribution of termination points across the cortex (**Extended Fig. 1d,e**). There were 105 ± 12 non-PCV ascending venules that terminated with highest density in the upper cortical layers but with some reaching ∼900 µm in depth. However, we found only 3.2 ± 0.5 vessels with PCV structure, corresponding to ∼3% of total ascending venules, consistent with *in vivo* data and confirming their sparsity relative to other vessel types.

PCV trunk diameters were on average 57.5 ± 15 µm in diameter (n = 23; mean ± SD), while other ascending venules averaged 19.8 ± 7.9 µm in diameter (n = 90; mean ± SD).^22^ This suggests that each PCV could support at least 8-times more blood flow than other individual ascending venules, facilitating drainage in broad regions of deep cortex and the CC.

### Defining the structure and nomenclature of PCVs

To understand age-related changes in PCV structure, we imaged adult (5-7 months old) and aged (22-24 months old) Thy1-YFP mice using *in vivo* deep two-photon imaging. With both age groups, PCV structure could be consistently visualized up to ∼1000 µm of depth (**Fig. 2a-d**). We developed a system to reliably identify the same vascular zones between animals and in different cortical layers (**Fig. 2e**). The main ascending vessel was denoted the “trunk” of the vessel, and larger vessels directly connected to the trunk were termed “branches”. Branches were numerous and of different diameters, with the largest diameter branches extending from the PVC trunk at greater cortical depths (**Supplementary Fig. 1**). This core branching structure of PCVs did not differ between age groups.

**Figure 2.**
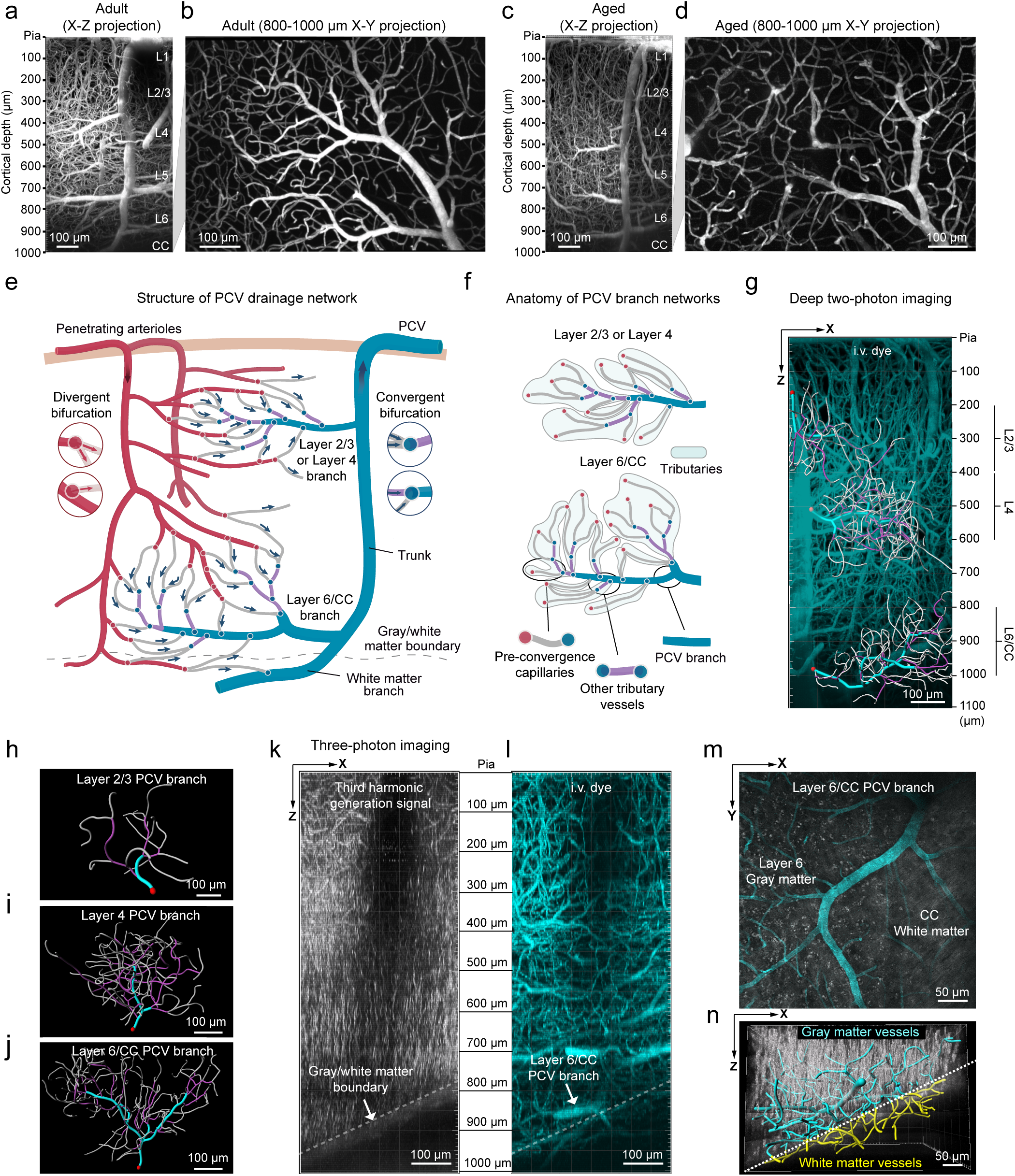
Structure of PCVs and their branching networks. **a-d.** Maximum projection images of deep two-photon image stacks showing PCVs in a 6 month old adult mouse (a,b), and a 22 month old age mouse (c,d). **e,f.** Schematic of vessel types and their nomenclature in PCV networks. **g.** Side projection of *in vivo* deep two-photon image stack showing a PCV in an adult mouse, overlayed with Imaris reconstructions of PCV branch vascular networks in layers 2/3, 4 and 6/CC. **h-j.** Top-down views of Imaris reconstructions of PCV branch vascular networks in layers 2/3, 4 and 6/CC, respectively. Red circles correspond to the point where the main branch joins the main trunk of the PCV. The gray-colored vessels correspond to pre-convergence capillaries, violet-colored vessels represent other tributary vessels, while the cyan-colored segments correspond to the main branch segments. **k, l.** Three-photon imaging of PCV branches concurrent with 3HG signal to define the gray-white matter boundary. Side projections of three-photon image stacks showing the 3HG signal fluorescence from myelin (k) and the vasculature labeled with the FITC-dextran i.v. dye (l) in an awake adult mouse (6 months old). The transition from gray to white matter is visible in the 3HG signal image (white dotted line). **m.** Maximum projection image showing a top-down view of the Layer 6/CC branch (750-1000 µm depth) from panel l. The white matter appears as a striated 3HG signal coming from axon bundles of the CC. **n.** Side view of reconstructed PCV branch in panel m. Vascular segments draining layer 6 and the CC are labeled in cyan and yellow, respectively. The dotted white line marks the location of the gray/white matter transition.

Each PCV branch received blood from dense networks of surrounding capillaries, which were characterized in detail. The common strategy of counting vascular branch order from penetrating vessel trunks in the upper cortex was insufficient to categorize these vessels.^23^ This was due to the difficulty in defining a “zero-order” vessel as PCVs ramified into large, horizontally oriented branches. Instead, we relied upon analysis of both microvascular branching structure and blood flow direction. Capillaries closer to arterioles receive blood flow from divergent bifurcations (**Fig. 2e**, red circles and left insets). On the venular side, capillaries drain into convergent bifurcations as blood flow merges (**Fig. 2e**, blue circles and right insets). Our initial characterization focused on vessels downstream of the last divergent bifurcation. We defined a capillary sub-type termed the “pre-convergence capillary”, which lies between the last divergent and first convergent bifurcation (**Fig. 2f**). Pre-convergence capillaries lie in the middle of the arterio-venous circuit. Downstream of pre-convergence capillaries are other capillary-sized vessels that we collectively termed “other tributary vessels”. Other tributary vessels are connected exclusively by points of convergence. Together, pre-convergence capillaries and other tributary vessels formed vascular units termed “tributaries”, and multiple tributaries could drain into PCV branches.

We analyzed larger PCV branches and their tributaries in layer 2/3, layer 4 and layer 6/CC (gray-white matter interface) because these networks were well separated from each other (**Fig. 2g**). Cortical layers were estimated based on the endogenous Thy1 YFP fluorescence of pyramidal cortical neurons in layer 2/3 and layer 5 (**Supplementary Fig. 2a**). As expected, there was a greater complexity in PCV branch structure with increasing cortical depth, with 6.1 ± 2.2, 21.7 ± 5.3 and 24.7 ± 5.6 tributaries per PCV branch in layers 2/3, 4 and 6/CC, respectively (n = 23 mice, aged groups pooled; mean ± SD)(**Fig. 2h-j**).

During deep two-photon imaging, the cortical depth of gray-white matter transition could be estimated based on a reduction in vascular density and shift to more planar capillary orientations (**Supplementary Fig. 2b**). However, to more clearly identify the gray-white matter transition, we also conducted *in vivo* three-photon microscopy in a subset of mice. Third-harmonic generation (THG) fluorescence allows visualization of myelin and vascular structure could be visualized with intravenously injected FITC-dextran (**Fig. 2k,l**). This imaging confirmed that layer 6/CC PCV branches were located at the gray-white matter interface, where vertically oriented myelin fibers in the cortex transition into horizontally oriented fibers of the CC (**Fig 2m**). Thus, layer 6/CC PCV branches drain blood from cortical layer 6 and CC (**Fig. 2n**) and deep multi-photon imaging can be used to study their structure and physiology *in vivo*.

### Simplification of tributary structure due to age-related regression of pre-convergence capillaries

We next characterized the architecture of PCV tributaries. Pre-convergence capillaries were the predominant vessel type in PCV tributaries, representing 60.12 ± 1.47% of the total vessel segments (n = 23 mice; mean ± SD) and 81.60 ± 4.95% of the total vessel length (n = 23 mice; mean ± SD)(**Fig. 3a,b**). Interestingly, the percentage of total vessel segments and total vessel length represented by pre-convergence capillaries increased with age in PCV tributaries across all cortical layers (**Fig. 3c, Supplementary Fig. 3a,b**). Yet, in deeper layers, there was a reduction in vascular length density (**Fig. 3d,e**, **Supplementary Fig. 3c,d**) and reduction in PCV tributary complexity (**Fig. 3f, Supplementary Fig. 3e-g**). This suggested that age-related regression of pre-convergence capillaries resulted in simplified PCV tributaries, affecting all cortical layers but layer 6/CC most strongly.

**Figure 3.**
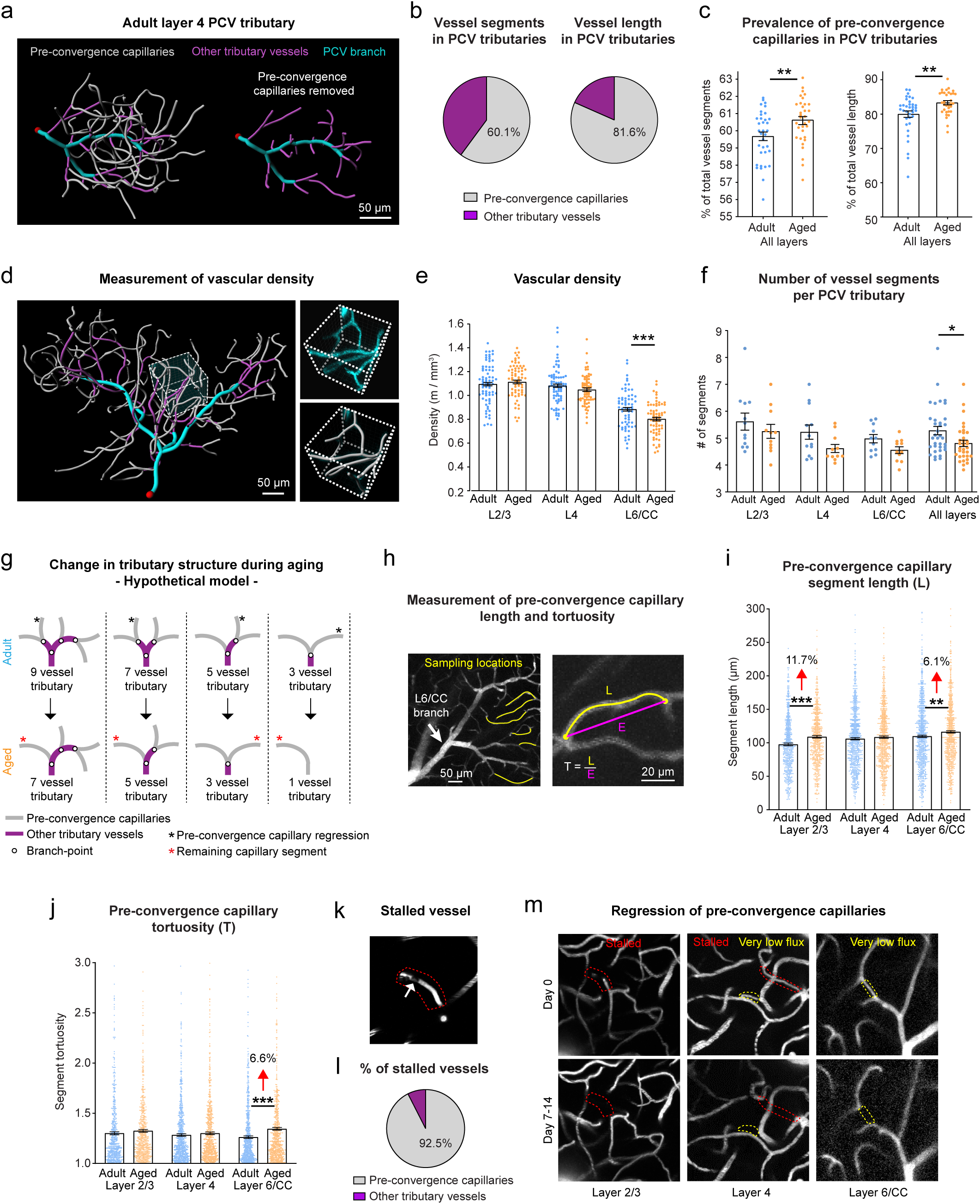
Reduced vascular density and simplified PCV tributary structure in aged mice. **a**. Reconstruction of adult Layer 4 PCV tributary with (left) and without (right) pre-convergence capillaries. **b.** Pie charts showing percentage of total vascular segments (left), and total vascular length (right) represented by pre-convergence capillaries and other tributary vessels in all PCV tributaries across layers and age. **c.** Age-related change in percentage of pre-convergence capillary segments and length in PCV tributaries (also see **Supplementary Fig. 3a,b**). **d.** Reconstruction of adult Layer 6/CC PCV tributary with example ROI for vascular length density measurements. Inset shows raw data and segmented vessels in an ROI. **e.** Vascular length density across layers in adult and aged mice. Two-way ANOVA with Holm-Sidak’s post-hoc test, Age group F(1,340)=8.679, p=0.003; Layer 2/3, p=0.401; Layer 4, p=0.119; Layer 6/CC, p<0.001. Adult, n=12 mice for layer 2/3 and layer 4, and n=11 mice for layer 6/CC; Aged, n=11 mice for all layers. Data shown as mean ± SEM. **f.** Average number of vascular segments in tributaries of adult and aged mice across layers. Two-way ANOVA with Holm-Sidak’s post-hoc test, Age group F(1,62)=6.186; p=0.016; Layer 2/3, p=0.264; Layer 4, p=0.063; Layer 6/CC, p=0.202. Adult, n=12 mice for layer 2/3 and layer 4, and n=11 mice for layer 6/CC; Aged, n=11 mice for all layers. Data shown as mean ± SEM. **g.** Hypothetical model for change in tributary complexity of aged mice involving pre-convergence capillary regression (marked with a gray asterisk). Also See **Supplementary Fig. 4**. **h.** Measurement of structural properties of individual pre-convergence capillaries. **i.** Length of individual pre-convergence capillaries. Two-way ANOVA with Holm-Sidak’s post-hoc test. Age group F(1,4127)=22.481; p<0.001; Layer 2/3, p<0.001; Layer 4, p=0.305; Layer 6/CC, p=0.007. **j.** Tortuosity of pre-convergence capillaries. Two-way ANOVA with Holm-Sidak’s post-hoc test. Age group F(1,4127)=12.924; p<0.001; Layer 2/3, p=0.267; Layer 4, p=0.351; Layer 6/CC, p<0.001. **k.** Example of a stalled blood flow in pre-convergence capillary observed *in vivo* (encircled with dotted red lines). Plugging cell marked with white arrow. **l.** Pie chart showing vessel types experiencing stalled flow across all layers and both ages. **m.** Examples of pre-convergence capillaries in adult mice exhibiting very low (yellow dotted lines) or stalled (red dotted lines) blood flow (upper row images) that regressed over time (bottom row images).

We developed a hypothesis for how simplification of tributary structure may be occurring (**Fig. 3g**, **Supplementary Fig. 4**). Age-related regression of pre-convergence capillaries (**Fig. 3g**, black asterisks) reduces vascular density and the number of vessels forming the tributaries. The neighboring pre-convergence capillary elongates and becomes more tortuous because a branchpoint is lost and the downstream tributary vessel merges into the same segment (**Fig. 3g**, red asterisk). The observed increase in percentage of pre-convergence capillary segments and length within PCV tributaries is consistent with this postulation (**Fig. 3c**). Also consistent with our hypothesis, we detected increases in both the length and tortuosity of individual pre-convergence capillary segments in aged mice (**Fig. 3h-j**). Since PCV tributaries represent a smaller subset of total vessels in upper cortical layers, their age-related changes were less apparent in general quantifications of vascular density. However, the overall data indicated that PCV tributary rarefaction was occurring across all cortical layers.

We next considered potential causes of age-related pre-convergence capillary regression. Given their central location in the arterio-venous circuit, they are the capillaries with the lowest blood flow, which increases the likelihood of transient plugging by leukocytes^24^ and capillary regression with prolonged flow arrest.^9,25^ In the *in vivo* image stacks, we occasionally observed stalls, *i.e.*, capillaries lacking movement of blood cell shadows (**Fig. 3k**, white arrow). Blood flow stalls were low in number across all tributaries (0.57 ± 0.16% of all vessels examined from both age groups; n = 23 mice; mean ± SD). However, nearly all flow stalls observed (92.5%) occurred in pre-convergence capillaries, even after normalization to total vessel length (**Fig. 3l, Supplementary Fig. 3h**). Further, instances of spontaneous pre-convergence capillary regression were observed in separate imaging sessions, 7-14 days apart, and affected vessels that had stalls or supported very low blood flow during prior imaging (**Fig. 3m**). Capillary stalling observations was low because each capillary was only imaged for brief periods, preventing statistical comparison. However, the prevalence of stalls appeared similar across cortical layers and age groups (**Supplementary Fig 3i**). It is expected that stalls and regressions would have more deleterious effects in the sparser networks of layer 6/CC.

### Altered structure and hemodynamics of pre-convergence capillaries in aged mice

Capillary diameter is a major determinant for blood flow resistance.^26,27^ We next measured vessel diameters and blood flow using line-scans in both isoflurane-anesthetized and awake mice (**Fig. 4a-c**). In a subset of mice, the same vessels were imaged under awake and anesthetized conditions for a direct comparison. This revealed significant reductions in pre-convergence capillary diameter in aged animals, with the most pronounced reductions in layer 6/CC in both anesthetized and awake conditions (**Fig. 4d,h**). Some diameter changes were small in magnitude (layer 4, anesthetized) and may not cause significant flow resistance. Interestingly, in awake mice, capillaries in layer 2/3 and 4 increased in diameter, and diameter reductions in layer 6/CC were larger (**Fig. 4h**). This difference may be attributed to isoflurane’s vasodilatory effects, as slight capillary dilation can obscure the true physiological differences in diameter between ages. Critically, the reduced diameter of capillaries was associated with pronounced age- and layer-specific reduction of RBC flux in layer 6/CC pre-convergence capillaries, irrespective of awake or anesthetized conditions (**Fig. 4e,f,i,j**) and sex (**Supplementary Fig. 5**). Further, RBC flux and velocity increased in layer 2/3 and layer 4 of aged mice (**Fig. 4e,f,i,j**), suggesting a redistribution of blood flow from deeper tissues to more superficial layers (**Supplementary Fig. 6,7**). The increase in capillary diameter and flow is consistent with prior imaging studies that have focused on upper cortex.^28,29^ Finally, aging may also be associated with reduced hematocrit, as linear density of RBCs was reduced across all layers in aged animals (**Fig. 4g,k**).^29^ These data emphasize the importance of measuring blood flow across cortical depth to understand the extent of age-related cerebral hypoperfusion.

**Figure 4.**
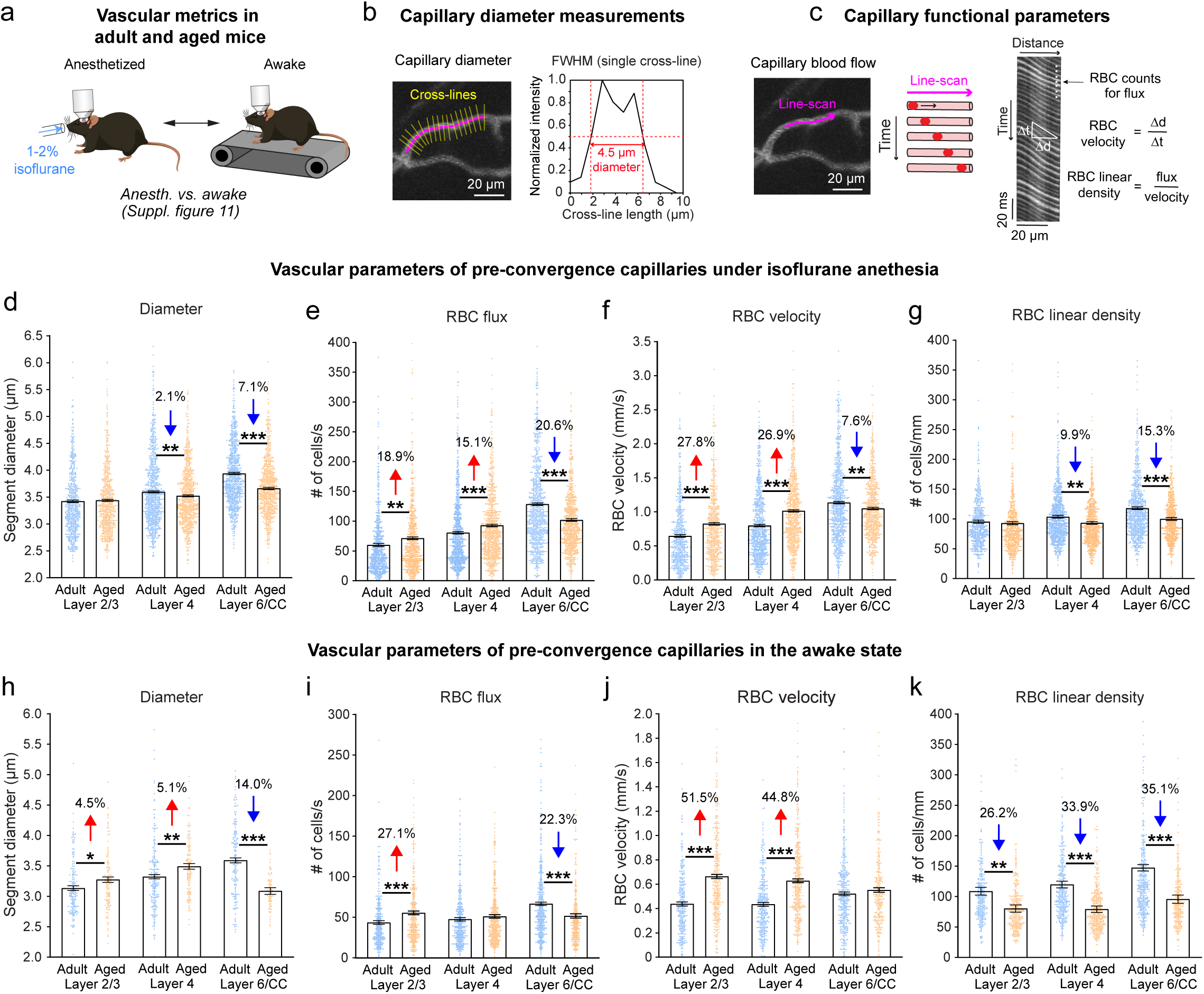
Aging involves a specific vasoconstriction and blood flow deficit in layer 6/CC. **a.** Imaging was performed in anesthetized and awake mice. For a subset of mice (**Supplementary Fig. 11**), the same vessels were imaged in both states. **b,c.** Assessment of capillary diameter and acquisition of line-scans for blood flow metrics. **d-g.** Diameter and blood flow parameters in pre-convergence capillaries in anesthetized mice. Diameter: Two-way ANOVA analysis with Holm-Sidak’s post-hoc comparison testing, Age group F(1,4147)=39.630; p<0.001; Layer 2/3, p=0.565; Layer 4, p=0.006; Layer 6/CC, p<0.001. RBC flux: Two-way ANOVA analysis with Holm-Sidak’s post-hoc comparison testing, Age group x Layer interaction F(2,4056)=48.065; p<0.001; Layer 2/3, p=0.001; Layer 4, p<0.001; Layer 6/CC, p<0.001. RBC velocity: Two-way ANOVA analysis with Holm-Sidak’s post-hoc comparison testing, Age group F(1, 4053)=55.672; p<0.001; Layer 2/3, p<0.001; Layer 4, p<0.001; Layer 6/CC, p=0.001. RBC linear density: Two-way ANOVA analysis with Holm-Sidak’s post-hoc comparison testing, Age group F(1,4043)=24.804; p<0.001; Layer 2/3, p=0.499; Layer 4, p=0.002; Layer 6/CC, p<0.001. Adult group includes n=12 mice for layers 2/3 and 4, and n=11 mice for layer 6/CC; the Aged group includes n=11 mice for all layers. Data shown as mean ± SEM. **h-k.** Diameter and blood flow parameters of pre-convergence capillaries in awake mice. Diameter: Two-way ANOVA analysis with Holm-Sidak’s post-hoc comparison testing, Age group x Layer interaction F(2,859)=33.596; p<0.001; Layer 2/3, p=0.019; Layer 4, p=0.003; Layer 6/CC, p<0.001. RBC flux: Two-way ANOVA analysis with Holm-Sidak’s post-hoc comparison testing, Age group F(2,1943)=20.674; p<0.001; Layer 2/3, p<0.001; Layer 4, p=0.227; Layer 6/CC, p<0.001. RBC velocity: Two-way ANOVA analysis with Holm-Sidak’s post-hoc comparison testing, Age group F(1, 1942)=130.757; p<0.001; Layer 2/3, p<0.001; Layer 4, p<0.001; Layer 6/CC, p=0.198. RBC linear density: Two-way ANOVA analysis with Holm-Sidak’s post-hoc comparison testing, Age group F(1,1928)=72.305; p<0.001; Layer 2/3, p=0.001; Layer 4, p<0.001; Layer 6/CC, p<0.001. Adult group includes n=5 mice for all layers; the Aged group includes n=6 mice for layer 2/3 and layer 4, and 5 mice for layer 6/CC. For plot h, the Adult group includes n=3 mice per layer; the Aged group includes n=2 mice per layer. Data shown as mean ± SEM.

We detected no age-related changes in the length and tortuosity in other tributary vessels, likely due to the shorter lengths of these vessel segments (**Supplementary Fig. 8a-d**). However, age-dependent reduction of vessel diameter (**Supplementary Fig. 8e, f**) and redistribution of blood over cortical layers was also evident in other tributary vessels (**Supplementary Fig. 9**). The observed blood flow differences in PCV tributaries were not due to altered cardiovascular status in aged mice, as we detected no major differences in heart rate in blood flow line-scans (**Supplementary Fig. 10**).

Reduced blood flow in deeper tissues could also be caused by constriction of upstream penetrating arterioles or their initial branches. However, we did not detect age-related difference in these upstream vessels in any cortical layer (**Extended Fig. 2**), supporting the idea that vasoconstriction and rarefaction in PCV tributaries was the primary driver of reduced blood flow.

Finally, in mice examined under both anesthetized and awake states, we leveraged isoflurane’s vasodilatory effect to probe for age-related differences in capillary reactivity. Isoflurane increased RBC flux and velocity over levels seen in awake mice in all cortical layers, with the largest increases in layer 6/CC (**Supplementary Fig. 11**). A similar magnitude of isoflurane-induced blood flow increase was seen with both age groups, suggesting a general preservation of reactivity. However, aging led to a reduced capacity to dilate capillaries in layer 6/CC, yet an increased capacity in layer 2/3, which may contribute to the abnormal redistribution of blood flow across layers.

### Vessel diameter is the main determinant of capillary flow in PCV networks

To examine which vascular changes were the strongest drivers of hemodynamic change, we performed Pearson correlation analyses between the structural and functional properties of pre-convergence capillaries in the anesthetized (**Supplementary Fig. 12 and Supplementary Table 1**) and awake state (**Supplementary Fig. 13 and Supplementary Table 2**). Vessel tortuosity was not correlated with RBC flux, but both capillary segment length and diameter were correlated. Of these two parameters, capillary diameter held the strongest relation with RBC flux, suggesting that local capillary diameter changes can strongly influence network perfusion.

### In silico modeling of age-related change in vascular structure recapitulates in vivo findings

To determine whether the degree of capillary constriction and rarefaction observed *in vivo* was sufficient to explain blood flow deficits seen with aging, we performed blood flow simulations in realistic microvascular networks derived from mouse barrel cortex.^30^ These vascular networks did not contain PCV trunks, but have been widely used and provided a close approximation to cortical regions studied *in vivo*. The *in silico* model based on these data is well-established and uses Poiseuille’s Law and the continuity equation to compute the pressure field and flow rates for all vessels.^30^ Moreover, the model accounts for the presence of red blood cells and their effect on the flow field (see Methods for additional details and model limitations).^31^

Four cases were studied in two microvascular networks, mimicking key changes seen in awake aged mice, where capillary diameter was increased in upper layers but decreased in layer 6/CC (**Fig. 5a**). Capillary density in layer 6/CC was also reduced to levels seen in aging. These modifications were sufficient to induce layer-specific blood flow changes. Most capillaries in layer 6/CC reduced in blood flow, while a smaller proportion of intermingled capillaries showed increased flow (**Fig. 5b**). The upper cortical layers exhibited scattered capillaries with increased capillary flow. On average, layer 6/CC showed a significant reduction of RBC flux, velocity and linear density with magnitudes exceeding *in vivo* levels (**Fig. 5c-e**). Total blood flow through arteriolar and venular branches across layers followed a similar pattern (**Fig. 5f, g**). Further, when capillary diameter change and density reduction were examined separately, the parameters had independent influences on reduction of RBC flux and velocity in layer 6/CC (**Supplementary Fig. 14**).

**Figure 5.**
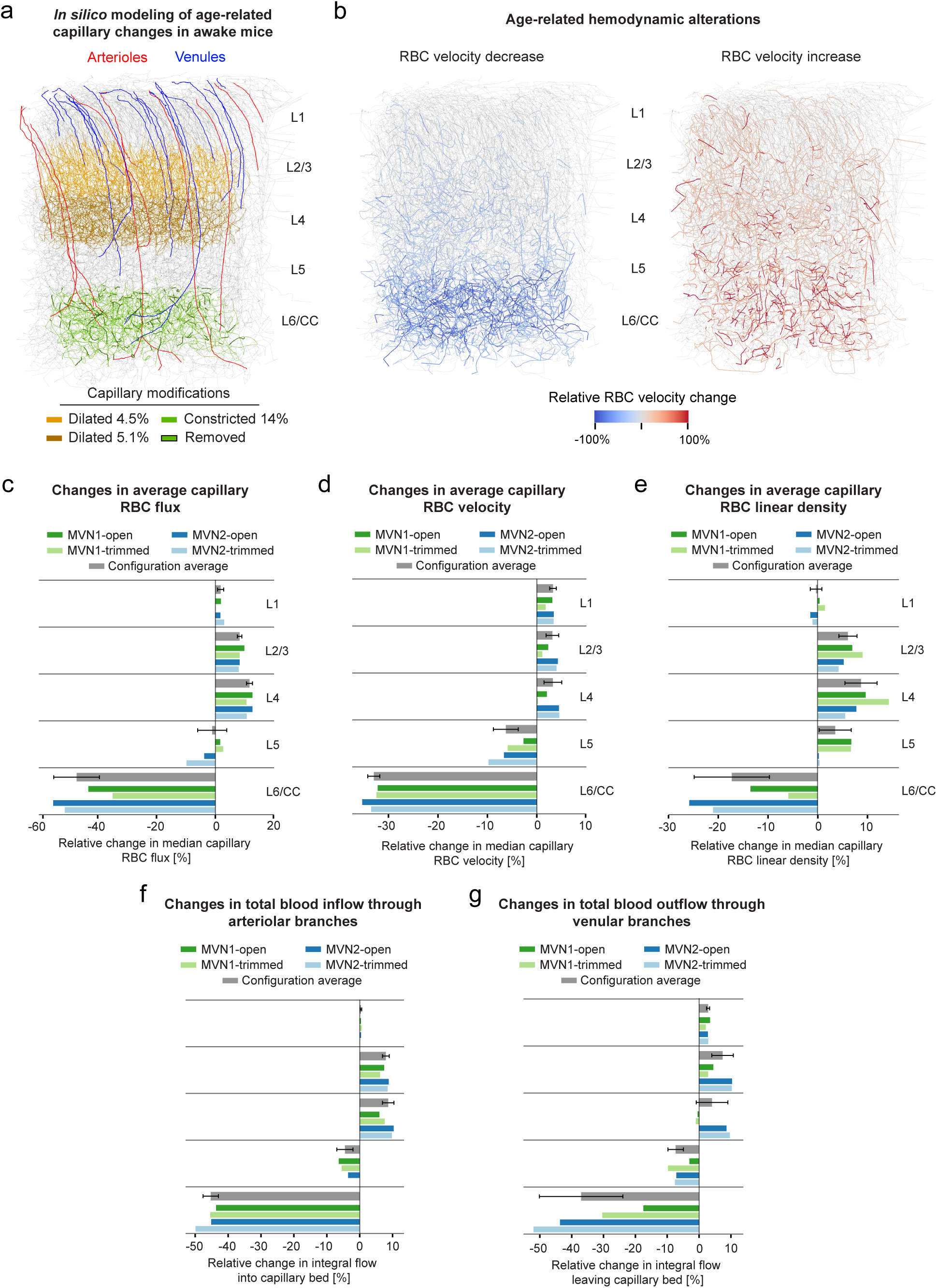
*In silico* modeling of capillary constriction and rarefaction captures features of age-related blood flow alteration. **a.** Image of an *in silico* microvascular network with labeled penetrating arterioles (red) and ascending venules (blue). Age-related structural changes from awake mice were implemented in the model, including capillary dilation (yellow segments), constriction (light green segments) and removed segments to mimic capillary regression (dark green segments). **b.** Image showing effect of age-related structural changes on microvascular network flow, with highlighted segments exhibiting reduced (left panel) or increased (right panel) RBC velocity. Only vessels with an absolute change larger than 10% are displayed. Panels (a) and (b) are MVN1-open configuration (see Methods). **c-e.** Changes in average capillary RBC flux (c), RBC velocity (d) and RBC linear density (e) across different cortical layers (n=4 network configurations, 2 microvascular networks x 2 boundary conditions). **f, g.** Changes in total blood inflow into the capillary network through arteriolar branches (f), and total blood outflow through venular branches (g) across different cortical layers (n=4 network configurations, 2 microvascular networks x 2 boundary conditions). All configuration average plots show median ± SD.

We also modeled capillary diameter reductions seen in anesthetized aged mice, which were specific to layer 6/CC. This yielded blood flow reductions similar to that observed *in vivo*, but did not induce blood flow increases in the upper cortex (**Supplementary Fig. 15**). This suggests that impaired flow to deep tissues does not simply reroute flow to upper layers, and that capillary dilations in upper layers may actively contribute to abnormal flow redistribution across layers. Overall, these simulations demonstrate that the magnitude of capillary changes seen *in vivo* are sufficient to produce marked changes in blood flow.

### Induced regression of pre-convergence capillaries in vivo causes broad capillary constriction and reduced blood flow in layer 6/CC

Since pre-convergence capillaries regress with age, we next mimicked this change in adult mice to explore its effect on local blood flow. In a previous study, we showed that optically-induced capillary regression in layer 2/3 caused upstream vasoconstriction and reduced capillary flow.^32^ Building on this, we conducted similar studies in deep PCV tributaries of awake adult mice using *in vivo* three-photon microscopy. Ablative line scans were used to rupture and induce the regression of pre-convergence capillaries in layer 4 and layer 6/CC. These regions were imaged before and at 3, 7 and 21 days after laser irradiation (**Fig. 6a**). Sham line scans irradiating the parenchyma in the immediate vicinity of the vessel were performed as controls (**Fig. 6b**).

**Figure 6.**
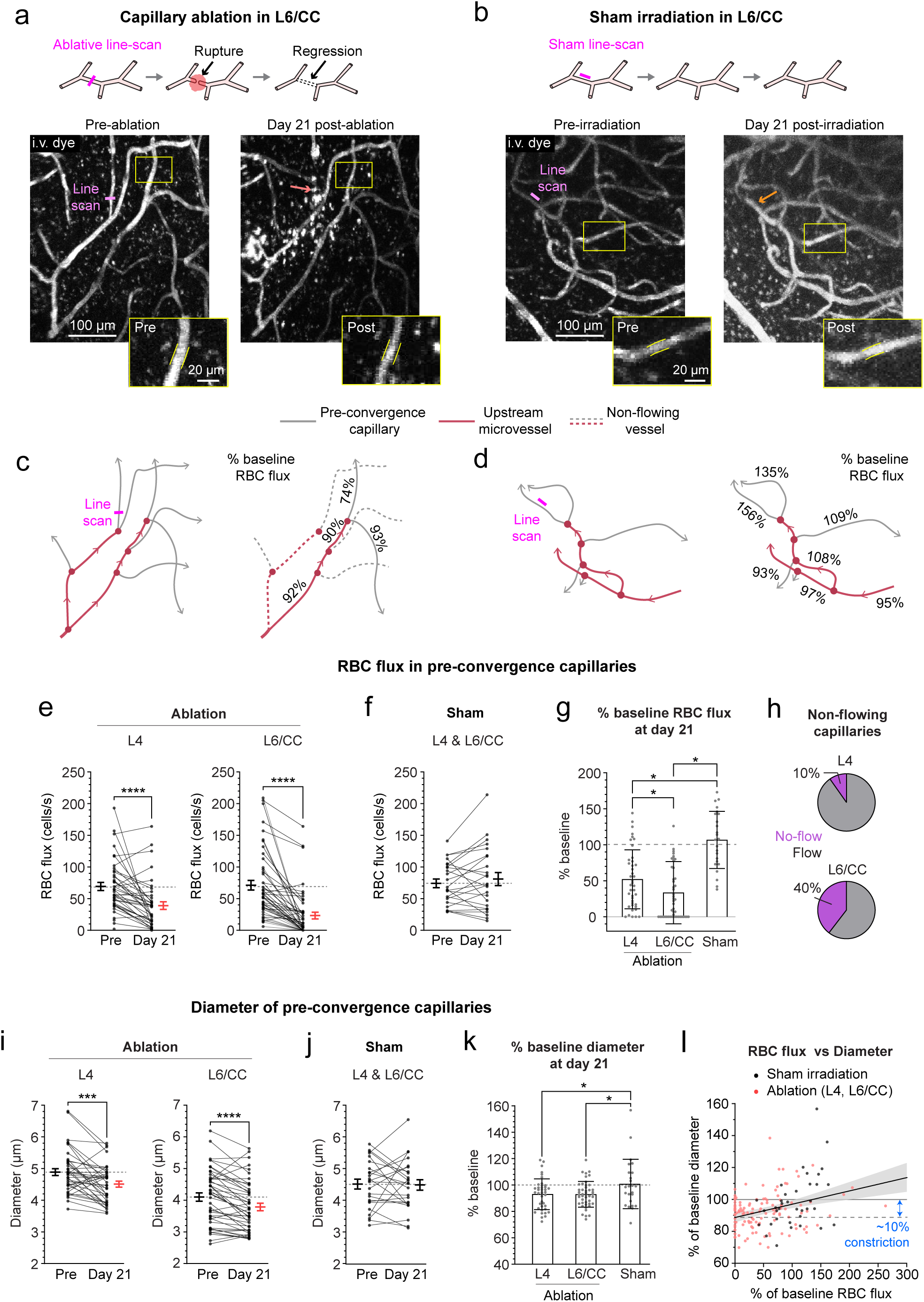
*In vivo* ablation of pre-convergence capillaries leads to broad vasoconstriction and hypoperfusion with worsened outcomes in layer 6/CC. **a,b.** Laser-induced ablation of pre-convergence capillaries in Layer 6/CC and off-target sham controls. Insets show changes evoked in a neighboring capillary before and 21 days post-irradiation. **c,d.** Vascular network schematics showing change in blood flow in panels (a) and (b). **e,f.** RBC flux of neighboring pre-convergence capillaries before and 21 days after capillary ablation in Layer 4 (left graph) and Layer 6/CC (right graph) in panel (e), or after Sham irradiation in panel (f). For Layer 4, n=8 regions of interest (ROIs), n=40 vessels; For L6/CC, n=9 ROIs, n=48 vessels, for Sham irradiation, n=4 ROIs, n=24 vessels. For all plots, data shown as mean ± SEM. Paired t-test. L4, p < 0.0001, t,df(4.804, 39); L6/CC, p < 0.0001, t,df (6.986, 47); Sham, p = 0.2998, t,df (1.061, 24). **g.** Percent change from baseline for pre-convergence capillary RBC flux at 21 days following laser irradiation. Data shown as mean ± SD. One-way ANOVA with Dunnett’s post-hoc comparison tests, p < 0.0001, F = 24.96 (2, 110). p < 0.05. **h.** Proportion of total capillaries with no-flow at 21 days after ablation. **i,j.** Pre-convergence capillary diameter before and 21 days after capillary ablation in Layer 4 (left graph) and Layer 6/CC (right graph) in panel (i), or after Sham irradiation in panel (j). For Layer 4, n=8 ROIs, n=40 vessels; for Layer 6/CC, n=9 ROIs, n=45 vessels; for Sham irradiation, n=4 ROIs, n=24 vessels. For all plots, Data shown as mean ± SEM. Paired t-test. Layer 4, p = 0.0003, t,df (4.004, 39); L6/CC, p < 0.0001, t,df (4.903, 44); Sham, p = 0.8694, t,df (0.1663, 24). **k.** Percent change from baseline for pre-convergence capillary diameter at 21 days following laser irradiation. Data shown as mean ± SD. One-way ANOVA with Tukey’s post-hoc comparison tests, p = 0.0357, F = 3.439 (2, 106). * p < 0.05. **l.** Change in RBC flux as a function of change in diameter across all capillaries examined. For all experiments, n = 5 adult mice (5-7 month old).

At day 21, regressions of single pre-convergence capillaries in layer 6/CC led to blood flow cessation or hypoperfusion in surrounding capillaries served by the same arteriolar branch (**Fig. 6a,c,e**). Blood flow reductions were more severe in layer 6/CC than layer 4 (70% vs. 50% reduction from baseline), with a more gradual worsening in deeper tissues (**Fig. 6e,g,h and Supplementary Fig. 16**). On average, sham irradiations performed across layer 4 and layer 6/CC did not affect blood flow, suggesting that these outcomes were not a result of non-specific laser damage (**Fig. 6f-h, Supplementary Fig. 16**).

Consistent with our prior studies in layer 2/3, capillary ablation resulted in general vasoconstriction in neighboring capillaries, while shams exhibited no change (**Fig. 6i-k, Supplementary Fig. 16**). Interestingly, relatively small vasoconstrictions in capillaries (∼10% from baseline) explained the range of blood flow changes observed (**Fig. 6l**), aligning with our observations in aged mice and *in silico* studies. Thus, experimentally induced capillary regression leads to hypoperfusion in local microvascular networks with worsened effects in deeper tissues. Further, capillary regression may be a trigger for broader hypoperfusion.

### Age-related reduction in blood flow is associated with white matter gliosis, demyelination and venous inflammation

We next determined whether age-related loss of vascular density and hypoperfusion in layer 6/CC was associated with tissue pathology. In mice we had imaged *in vivo*, we also extracted brain tissues and immunostained for markers of microglia (Iba1), astrogliosis (GFAP) and myelin (myelin basic protein, MBP)(**Supplementary Fig. 17**). Consistent with prior studies^33^, aged mice exhibited increased microgliosis and astrogliosis, as well as demyelination and reduced YFP fluorescence of axons in the CC (**Supplementary Fig. 18)**. These histological indices of tissue pathology correlated with the average RBC flux in layer 6/CC when examining both ages together (**Supplementary Fig. 18c,f,i,l**).

In a second cohort of mice examined histologically, but not imaged *in vivo,* we included adult (5-7 months), mid-aged (16-18 months), and aged (20-22 months) groups to better understand the temporal sequence of events in mid-cortex and layer 6/CC. We again examined markers of gliosis and myelin, but also added markers to assess vascular structure, inflammation and tissue hypoxia. Age-dependent increase in gliosis and demyelination was again observed in layer 6/CC, but not mid-cortex (**Fig. 7a-c**), reproducing outcomes from the first cohort and confirming that tissue changes were not an effect of cranial window surgery or *in vivo* imaging. Critically, a significant loss of vascular density was seen in both mid-cortex and layer 6/CC by mid-age, suggest that it occurs early enough to be a potential contributor to age-related gliosis and demyelination in layer 6/CC (**Fig. 7d**). VCAM1, a leukocyte adhesion molecule, was strongly upregulated in the CC of aged mice, with little to no staining in adult or mid-aged groups (**Fig. 7e**). Prominent staining was observed in PCV trunks and layer 6/CC capillary networks (**Fig. 7f.g**), suggesting a contribution to capillary stalling in white matter.

**Figure 7.**
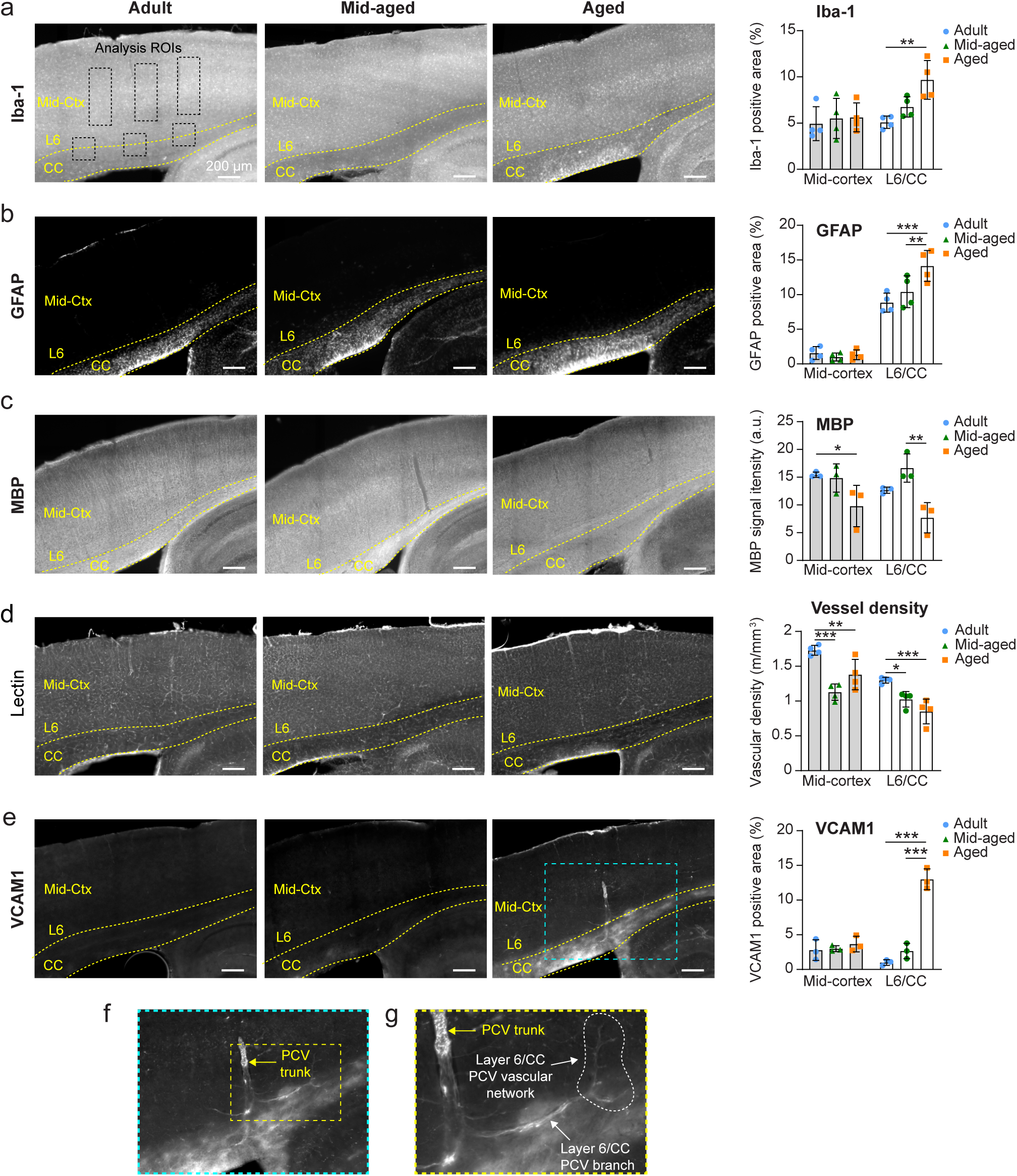
Gliosis, demyelination and reduced vascular density in the corpus callosum of aged mice. **a-e.** Epi-fluorescent images of adult (left column), mid-aged (center column) and aged (right column) mouse somatosensory cortex from sagittal brain sections with accompanying analyses across age and brain regions. Tissues were stained with anti-Iba1 antibody for microglia (a), anti-GFAP antibody for astrocytes (b), anti-MBP antibody for myelin (c), tomato lectin for labeling all blood vessels (d), and anti-VCAM1 antibody for VCAM1 expression (e). Bar plots showing assessment of immuno-staining levels in mid-cortex and layer 6/CC, with analysis ROIs similar to that shown in panel (a). For plots in panels (a-c), n=4 adult, n=4 mid-aged, and n=4 aged mice. For plots in panels (d,e), n=3 adult, n=3 mid-aged, and n=3 aged mice. For all plots, data shown as mean ± SD. Two-way ANOVA with Holm-Sidak post hoc testing. *p<0.05, **p<0.01, ***p<0.001. **f,g**. Magnified insets from panel (k) showing VCAM1 staining in PCV trunk and branching networks of Layer 6/CC

The vascular deficits observed with aging in layer 6/CC did not appear to cause significant hypoxia or induce hypoxic responses. A pimonidazole-HCl probe (Hypoxyprobe), which is sensitive to hypoxia (<10 mmHg pO_2_), was injected *in vivo* for *post-mortem* immunostaining and showed no elevation in aged mice (**Supplementary Fig. 19a,b**). Further, qPCR analyses in mid-aged and aged mice revealed no increase in expression of hypoxia-responsive genes with aging (*slc2a1, ldha, vegfa, pdk1*), which aligns with prior studies (**Supplementary Fig. 19c-f)**.^8^ On the contrary, some of these genes trended toward reduced expression, possibly reflecting an age-related dampening of HIF-1-alpha activity. Thus, deeper tissues do not become overtly hypoxic with age but experience a mild but chronic restriction in metabolic supply.

### Decreased capillary pericyte density in layer 6/CC of aged mice

Pericytes are mural cells that line brain capillaries and their contractile tone modulates basal capillary flow.^23^ Pericytes are lost at higher levels than endothelial cells during aging^10,34^, and experimentally-induced pericyte loss in adulthood causes capillary blood flow stalling, reduction in capillary structure, and leukocyte adhesion.^35,36^ We therefore examined how aging affected pericyte density in somatosensory cortex using our light-sheet imaging data (**Extended Fig. 3**). Aging was associated with a reduction in vascular length density and pericyte numbers starting from 400 µm of intracortical depth, which is approximately where layer 4 PCV branching networks are located (**Extended Fig. 3k, l**). When pericyte numbers were normalized to the total vascular length, a specific age-related reduction in pericyte density was detected at 800-1000 µm of depth, corresponding to layer 6/CC (**Extended Fig. 3m**). These data suggest that pericyte loss/dysfunction could be a contributor to age-related capillary changes.

### Mild hypoperfusion is sufficient to drive gliosis and advance toward demyelination in layer 6/CC

Our in vivo two-photon imaging revealed that relatively mild levels of blood flow reduction (∼20%) were associated with age-related tissue pathology in layer 6/CC. To investigate whether this degree of hypoperfusion is sufficient to cause tissue pathology, we induced a similar level of hypoperfusion in adult mice (5-7 months old) and examined markers of gliosis and myelin integrity. Mild hypoperfusion in one brain hemisphere was achieved through unilateral common carotid artery stenosis (UCCAS) (**Fig. 8a, b**) using a 0.20 mm radius micro-coil implanted around the animal’s left common carotid artery. Cranial windows were also implanted to assess capillary RBC flux changes in layer 4 and layer 6/CC at baseline and 3, 7, 14, and 21 days after UCCAS initiation. Brain tissues were then collected for histological assessment, with the contralateral hemisphere serving as an internal control.

**Figure 8.**
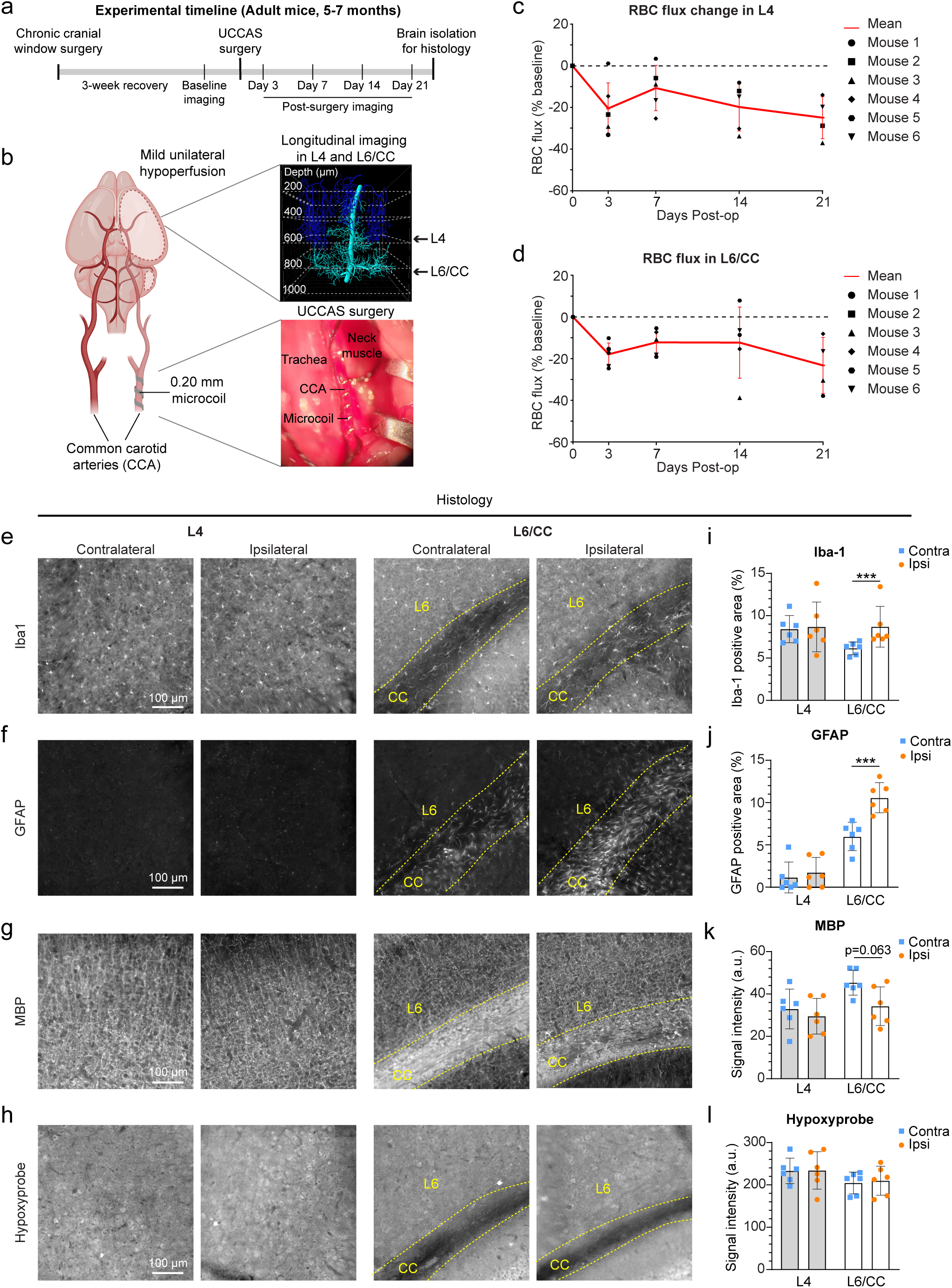
Mild unilateral hypoperfusion is sufficient to induce gliosis and demyelination in the ipsilateral corpus callosum. **a.** Schematic showing experimental timeline. **b.** Schematic on the left shows placement of a 0.20 mm microcoil around the common carotid artery to induce mild unilateral hypoperfusion. Images on the right show longitudinal *in vivo* deep-two photon imaging performed on ipsilateral somatosensory cortex Layer 4 and Layer 6/CC PCV branching networks (top), and microcoil placement during the UCCAS surgery (bottom). **c, d.** Average RBC flux over time in Layer 4 (c), and Layer 6/CC (d), collected in pre-convergence capillaries of PCVs in animals that underwent UCCAS surgery. Data presented as mean ± SD. Each dot in the graphs represent an average RBC flux of a population of pre-convergence capillaries from a PCV branch in a single animal. **e-h.** Epi-fluorescent images of contralateral and ipsilateral Layer 4 (left side) and Layer 6/CC (right side) from somatosensory cortex, stained with anti-Iba1 antibody for microglia (e), anti-GFAP antibody for astrocytes (f), anti-MBP antibody for myelin (g), and anti-Hypoxyprobe for labeling hypoxic tissue (h). **i-l.** Corresponding plots showing analyses of Iba-1 positive area (i), GFAP positive area (j), MBP signal intensity (k), and Hypoxyprobe signal intensity (l). For all plots, n=6 mice, Data shown as mean ± SD. Two-way ANOVA with Holm-Sidak post hoc testing. ***p<0.001. Iba1: Hemisphere x Layer Interaction effect F (1,5) = 27.48, p = 0.0033; GFAP: Hemisphere x Layer Interaction effect F (1,5) = 25.82, p = 0.0038; MBP: Hemisphere effect F (1,5) = 25.77, p = 0.0038; (l) Layer effect F (1,5) = 0.1858, p = 0.6844.

On average, both layer 4 and layer 6/CC exhibited relatively stable ∼20% reductions in capillary RBC flux over three weeks of imaging following UCCAS (**Fig. 8c,d**), comparable to the magnitude of blood flow decrease observed in layer 6/CC of aged mie (**Fig. 4e,i**). Histological analyses revealed increased Iba-1 and GFAP staining in the ipsilateral layer 6/CC, while layer 4 was unaffected (**Fig. 8e,f,i,j**). Mild hypoperfusion also produced a strong trend toward reduced myelination in layer 6/CC (**Fig. 8g,k**). No differences were seen with Hypoxyprobe staining, indicating that mild hypoperfusion caused tissue oxygen changes sub-threshold for probe detection, as seen with aging. These findings confirm that the degree of blood flow reduction occurring with aging is sufficient to drive tissue pathologies in layer 6/CC and is thus a key contributor to white matter injury and gliotic reactions.

## DISCUSSION

Cerebral white matter is highly vulnerable to hypoperfusion during aging and neurological disease, yet its supporting microvascular networks remained poorly defined. Using deep multi-photon imaging, we show that cortical layer 6 and CC in mice are exclusively drained by rare, wide-reaching penetrating venules called PCVs. Age-related capillary constriction and rarefaction within the deep branches of PCVs lead to mild hypoperfusion (∼20% decrease) in layer 6/CC, which is strongly linked to microgliosis, astrogliosis, and myelin loss. Moreover, inducing a comparable level of hypoperfusion in adult mice using common carotid artery stenosis resulted in similar tissue pathologies in layer 6/CC. These findings identify PCVs as critical bottlenecks and sources of flow impedance during aging. Given that murine cortical-CC vascular architecture mirrors that of the human cortical-U fiber interface^19^, similar mechanisms may contribute to age- and disease-driven degeneration of superficial white matter in humans.

Pre-convergence capillaries in PCV tributaries were the most vulnerable to blood flow disturbance and rarefaction. Regressed capillaries tended to be those that had previously stalled or supported very low blood flow, reinforcing a growing link between imbalanced capillary flow and the progressive loss of capillary network structure during aging.^9,25,35^ The abnormal contraction or death of pericytes in layer 6/CC can explain these flow disturbances^23,35,37^, though the triggers of age-related pericyte pathology remain unclear. Concurrently, the endothelium and surrounding glia show heightened expression of immune- and inflammation-related genes, particularly in the white matter^13,34,38^ Consistent with endothelial inflammation, we find that VCAM1 expression and pericyte loss are most prominent in layer 6/CC. Previous studies suggest that pericyte loss could drive endothelial inflammation and increased leukocyte adherence.^36^ Further, the white matter vasculature appears to be enriched in venous-associated vessels, where endothelial-leukocyte interactions are likely to be heightened. Thus, a cycle of capillary constriction and obstruction, metabolic insufficiency, pericyte loss, and vascular inflammation may perpetuate white matter deterioration. Future research should focus on identifying the specific molecular changes in white matter vasculature, leveraging emerging techniques such as cellular-resolution spatial transcriptomics.

Pericytes loss has been shown in clinical neuropathological studies of the aging brain^39^, and its effects have been broadly studied in experimental models that either block pericyte survival signals or to directly ablate pericytes in adulthood.^35,40,41^ However, the pathophysiological signals that trigger abnormal pericyte contraction are underexplored. More research is needed to understand the mechanism of capillary-level vasoconstriction, which may differ from arterial signaling, and to determine effective strategies to curtail this process.^42^ Pericytes contraction could be driven by membrane depolarization and calcium influx or by increased production of vasoconstrictive agonists such as endothelin-1 or thromboxane A2.^43–45^ Other potential factors also cannot be ruled out, including changes in the structure of the endothelium, basement membrane or other perivascular components. Our findings indicate that deep capillaries in the aged brain remain responsive to vasodilatory stimuli such as isoflurane, suggesting that clinically-approved therapeutics with known vasodilatory effects on pericytes and capillaries may still be effective.^23,46^ Recent studies have also shown that exercise can restore age-related vascular deficits in the CC.^47^

We found that capillary flux increases with cortical depth in the adult brain and this gradient is dampened during aging, consistent with prior work.^47^ The maintenance of higher blood flow rates in deeper tissue is likely necessary to compensate for the reduced oxygen content of RBCs as they travel farther from the arterial source and encounter sparser capillary networks. Supporting this idea, an *in vivo* study by Li et al. reported depth-dependent decreases in partial oxygen pressure (pO_2_) in penetrating arterioles and ascending venules.^48^ We postulate that the ∼20% reduction in RBC flux in layer 6/CC of aged mice lowers pO_2_ to levels that compromise metabolic function in white matter, but to an extent that is insufficient to trigger compensatory responses to hypoxia and remains undetectable by hypoxyprobe. Consequently, even modest age-related declines in blood flow can contribute insidiously to white matter dysfunction and tissue degradation. When compounded by age-related capillary rarefaction, localized regions of low pO_2_ become more pronounced, increasing susceptibility to secondary insults.^49,50^ For example, in a mouse model of increased AD risk (Apoe4), additional vascular insufficiencies in the CC further reduce tissue pO_2_ to levels detectable by hypoxyprobe (< 10mmHg).^12^ The development of *in vivo* multi-photon-based techniques to measure pO_2_ at depth will be an important advancement for testing these concepts.

In summary, deep *in vivo* multi-photon imaging is poised to reveal new insights into the etiology of cerebrovascular disease and complement clinical studies. Our study establishes a framework for navigating the complex microvascular networks of the deep cortex and CC in mice, which is essential for consistent measurements within and across animals. We established a baseline to understand microvascular change in disease models that incorporate aging as a factor. When combined with the broad-scale visualization of microvasculature enabled by light-sheet microscopy^10,51,52^, *in vivo* studies can now focus on specific brain regions most vulnerable during aging and disease.

## METHODS

### Mice

All procedures in this study were approved by the Institutional Animal Care and Use Committee at the Seattle Children’s Research Institute. The institution has accreditation from the Association for Assessment and Accreditation of Laboratory Animal Care (AAALAC), and all experiments were performed within guidelines. *In vivo* deep two-photon imaging studies comparing age differences were performed on adult (n = 12; 6 male and 6 female, 5 to 7-month old) and aged (n = 12; 9 male and 3 female, 22 to 24-month old) Thy1-YFP mice^53^ bred on the C57Bl/6 background (B6.Cg-Tg(Thy1-YFP)HJrs/J; Jax ID 003782). The number and sex of mice for other experiments are listed below in each relevant section. Age group ranges were determined by the Jax “Life Span as a Biomarker” criteria, which defines 3-6 months as “mature adult” stage, and 18-24 months as “old” stage in mice. Aged mice were visually checked for general health and any overt signs of illness such as tumors or weight loss resulted in exclusion from the study. The mice were housed on a 12-h light (6:00 to 18:00), 12-h dark cycle, with *ad libitum* access to chow and water. The use of adult and aged mice was interleaved over the entire period of study.

### Chronic cranial windows

Dexamethasone (40 µl; Patterson Veterinary) was subcutaneously administered to animals 4 h prior to the surgery, which helped to reduce brain swelling during the craniotomy. Anesthesia was induced with a cocktail consisting of fentanyl citrate (0.05 mg/kg), midazolam (5 mg/kg) and dexmedetomidine hydrochloride (0.5 mg/kg) (Patterson Veterinary). Under sterile conditions, the scalp was excised, and the periosteum cleaned from the skull surface. An aluminum flange for head fixation was attached to the right side of the skull surface using the C&B-Metabond quick adhesive cement (Parkell; S380). A circular craniotomy (dura left intact), ∼4 mm in diameter, was created over the left hemisphere and centered over 2-mm posterior and 3-mm lateral to bregma encompassing the primary somatosensory cortex. The craniotomy was sealed with a glass coverslip plug consisting of a round 4 mm glass coverslip (Warner Instruments #64-0724) glued to a round 5 mm coverslip (Warner Instruments #64-0731) with UV-cured optical glue (Edmund optics #37-322). The coverslip was positioned with the 4 mm side placed directly over the craniotomy, while the 5 mm coverslip laid on the skull surface at the edges of the craniotomy. Loctite Instant Adhesive 401 was carefully dispensed along the edge of the 5 mm coverslip to secure it to the skull. The area around the cranial window was then sealed with dental cement. Throughout surgery, body temperature was maintained at 37°C with a feedback-regulated heat pad (FHC Inc.), and the mice were provided medical air through a nose cone (20–22% oxygen and 78% nitrogen, moisturized by bubbling through water; AirGas Inc.). Imaging was initiated after a 3-week recovery period.

### In vivo deep two-photon imaging (isoflurane anesthesia)

Mice were maintained under light isoflurane anesthesia (∼1.5% minimum alveolar concentration) delivered by medical air (20– 22% oxygen and 78% nitrogen, moisturized by bubbling through water; AirGas Inc.), and body temperature was maintained at 37°C with a feedback-regulated heat pad (FHC Inc.) throughout the imaging period. To label the vasculature, 100 μL of 5% (w/v in sterile saline) 2MDa Alexa 680-dextran was injected through the retro-orbital vein. Alexa 680-dextran was custom conjugated using a previously published protocol.^20^ *In vivo* two-photon imaging was conducted using a Bruker Investigator microscope (run by Prairie View 5.5 software) coupled to a Spectra-Physics Insight X3 laser source (SpectraPhysics). Endogenous YFP fluorescence and Alexa 680-labeled microvasculature was imaged at 900 nm and 1210 nm excitation, and emission was collected through 525/70-nm and 700/75-nm bandpass filters, respectively. During imaging, laser power ranged between 4 and 145 mW exiting the microscope objective, with higher powers required for greater cortical depth (**Supplementary Fig. 20**).

### Imaging timeline

Each mouse underwent 5 sessions of *in vivo* deep two-photon imaging under anesthesia with <2.5 h of imaging time per session and 2-3 days between the sessions. During the first imaging session, low-resolution maps of the cranial window were collected using a 4-X 0.16 numerical aperture (NA) objective (Olympus; UPlanSAPO) for navigational purposes, as well as to identify PCV locations. Once the appropriate PCV was located, high-resolution imaging of PCV branches and associated microvascular networks was performed through the entire cortical depth. Image stacks were collected using a 25-X, 1.05 NA water-immersion objective lens (XLPlan N, Olympus) across a 483 μm x 483 μm field of view with lateral sampling resolution (x, y) of 0.943 µm/pixel and axial sampling resolution (z) of 1 µm/pixel. Two to four image stacks were often collected and stitched using an ImageJ/FIJI “Pairwise stitching” plugin to cover the region of interest.^54^

We analyzed PCV branches originating directly from the main PCV trunk between 200-400 µm, 400-600 µm and 800-1000 µm of intracortical depth, roughly corresponding to cortical layers 2/3, 4 and 6/CC, respectively (**Fig. 2g**). Branches at these depths drained blood from non-overlapping regions and generally covered upper, mid and deep layers of the cortex. To confirm the location of cortical layers 2/3 and 4 we examined endogenous fluorescence in pyramidal neurons of layers 2/3 and 5 (**Supplementary Fig. 2**). 3D visualization of high-resolution Z-stacks was performed using Imaris software version 7.7.2 (Bitplane, Oxford Instruments). Vascular segments presumed to belong to a PCV branch network were identified based on the evaluation of vascular structure from the image stacks. To ensure that we were including the correct starting points of the networks, all vessel segments starting from one branch order upstream of the putative starting points of the PCV network and downstream to the main branches of the PCV were marked. Then, the location of the marked vascular segments was labeled in the original Z-stack using ImageJ/FIJI to provide a map of where line scanning was to be performed in subsequent imaging sessions.

### Line scan data acquisition

During *in vivo* deep two-photon imaging sessions 2, 3 and 4 under anesthesia, line scan data were collected from vessel segments in line scan maps from layer 2/3, 4 and 6/CC (one cortical layer per session). Line scanning was performed using the 25-X, 1.05 NA water-immersion objective lens (XLPlan N, Olympus) and 3-X digital zoom was used to guide accurate placement of the line scan. Vessel segments were sampled individually with line scan duration set to ∼1.2 s at a sampling frequency of ∼2 kHz. During acquisition of line scan data, we also followed the number of stalled vessel segments. Vessel segments were considered stalled if they had no blood flow for more than 20 minutes. These vessels were reassessed periodically over the imaging session. To ensure that the laser powers used for line scans did not directly induce damage or alter blood flow, we longitudinally imaged the same population of Layer 6/CC vessels weekly for 5 weeks in a cohort of mice (**Supplementary Fig. 21**). We observed no overt signs of vascular damage, *i.e.*, dye extravasation indicative of blood-brain barrier damage or vascular regression. Blood flow was consistent and unperturbed across all time points examined.

### Analysis of the PCV branches over depth

During *in vivo* deep two-photon imaging session 5, we collected high-resolution Z-stacks through the entire cortex, starting at the pia mater and ending at ∼1000 µm of cortical depth. Lateral sampling (x, y) was 0.943 µm/pixel, and axial sampling (z) was 1 µm/pixel. As mentioned above, to capture the entire branching structure of the PCV we collected and stitched 4 adjacent and partially overlapping z-stacks arranged in a 2x2 square formation, with the main PCV trunk set in the center. Using the stitched Z-stacks, we analyzed the overall branching structure of PCVs in adult and aged mice. We measured the diameter of each PCV branch segment right next to the point where it joins the main trunk of the PCV, as well as the cortical depth at which the branch originates. Furthermore, we analyzed the number of PCV branches through cortical depth, after binning in 200 µm depth groups.

### Awake imaging

A subset of Thy1-YFP mice (n = 5 adult, 2 male and 3 female; and n = 6 aged, 5 male and 1 female) was imaged both under isoflurane anesthesia (as described above) and in the awake state. During awake imaging, mice were head fixed and allowed to freely move on a low resistance treadmill (Phenosys SpeedBelt). All mice were habituated for head fixation over 3 training sessions (each ∼2 days apart) where mice were head-fixed on the treadmill and placed in the light-shielded microscope cage with the objective over the cranial window to imitate the conditions during imaging. The mice were then held in the imaging setup for 20 min before being returned to the home cage. After training, each mouse underwent 7 imaging sessions over 3.5 weeks (2 imaging sessions per week, 3 days apart). The first session was performed under isoflurane anesthesia to collect low-resolution maps of the cranial window for navigational purposes, as well as to identify the location of PCVs, as described above. Once the appropriate PCV was located, high-resolution imaging was performed throughout the entire depth around the PCV. In the following 6 imaging sessions, line scan data were collected from vessel segments from layer 2/3, 4 and 6/CC (one cortical layer per session). We alternated imaging of the same cortical layer in the awake state and under anesthesia on separate sessions in immediate succession to obtain hemodynamic data from the same population of vessels under both conditions. At the end of each awake imaging session, a Z-stack of the imaged PCV vascular network was collected for analysis of vessel diameter.

### Defining the vascular networks of a PCV branch

We defined that PCV branch networks should include all vessel segments that converge blood flow towards that branch. Therefore, we considered the last points of blood flow divergence within the capillary network as the starting points for the PCV branch vascular network. Using 3D structural information from our image stacks and blood flow directionality extracted from line scan data, we were able to reliably identify the last points of convergence in the analyzed PCV vascular networks.

The first vessel segments in the PCV branch vascular network are located between the points of flow divergence and points of flow convergence, termed “pre-convergence” capillaries. Pre-convergence capillaries merge and form a system of downstream convergent vessel segments, termed “other tributary vessels”, which eventually lead to a larger diameter PCV branch. Pre-convergence capillaries and other tributary vessels were together termed “tributaries”. PCV branches were generally larger diameter offshoots from the PCV trunk and multiple tributaries of differing complexity could flow into branches, collectively forming PCV branching networks (**Fig. 2e, f**).

### Analysis of tributary structure

As the capillary network is predominantly composed of bifurcating branchpoint, the number of vessel segments in tributaries is almost always an odd number **(Fig. 3g)**. To analyze tributary structure, we measured the average number of vessel segments per tributary and separated tributaries into 3 groups: low (1 or 3 vessels), medium (5 or 7 vessels) and high (9 or more vessels) complexity. In the case of a tributary containing a trifurcation point and consisting of an even number of vessel segments, it was categorized as the first higher odd number tributary (i.e., 4 as 5, 6 as 7 etc.). For each analyzed PCV vascular network, we calculated the total number of tributaries belonging to a certain complexity group (low, medium, high) and expressed it in relation to the total number of tributaries in the PCV network (**Supplementary Fig. 3e-g**).

### Segmentation of PCV vascular networks

Vascular networks of PCV branches in Z-stacks were reconstructed in 3D using the Imaris, version 7.7.2 (Bitplane, Oxford Instruments). Vessel segmentation was performed manually using the “Auto Depth” tracing option in the Filament Tracer module of Imaris. “Center” and “Smooth” options in the Edit tab of the Filament tracer module were used to correct for any tracing mistakes. After segmentation of the networks, the length and straightness of each reconstructed vessel segment was extracted from the Statistics tab of the Filament tracer module. The straightness values reported by the software are reciprocal to tortuosity.

### Vessel diameter measurement

Lumen diameter of different vessel types was measured in maximally projected images from high-resolution Z-stacks. Maximum projections of 10-40 µm in thickness were used for analysis, dependent upon the size of the vessels. To reduce bias of measurement location, we used a custom ImageJ/Fiji macro called VasoMetrics to analyze lumen full-width at half maximum diameter at multiple, equidistant locations (spaced 1 µm) along each vessel segment of interest.^55^ The values across each vessel segment were used to calculate the average diameter of the vessel and the standard deviation of diameter along the measured region.

### Quantification of hemodynamic parameters and stalled vessels

For each line-scan captured, we calculated RBC flux by manually counting the number of blood cell shadows over the total duration of the line scan and represented the values as cells per second. The angle of the streaks in relation to the direction of the scan was used to determine RBC flow direction. Analysis of blood flow velocity and heart rate was performed using a previously published MATLAB algorithm.^56^ Linear RBC density values for each vessel segment were calculated by dividing the RBC flux with the blood flow velocity. Blood flow stalls were identified as capillaries lacking moving blood cell shadows for at least a period of 20 min during repeated observations within the same imaging session.

### Analysis of vascular density

Analysis of vascular density was performed in two ways using the Imaris software. First, we assessed vascular density (total vascular length per volume) in regions enriched with pre-convergence capillaries. This involved placing 6 small regions of interest (100 x 100 x 100 µm (x, y, z)) within each reconstructed PCV network (**Fig. 3d,e**).

Second, vascular density was measured more broadly in cortical layers by assessing larger regions of interest (300 x 300 x 100 µm (x, y, z)) within layers 2/3, 4 and 6/CC (**Supplementary Fig. 3c, d**). Two ROIs were analyzed per each layer, for a total 300 x 300 x 200 µm (x, y, z) ROI volume.

### In vivo three-photon imaging

To identify the gray-white matter transition, adult C57BL/6 mice (n = 3, 2 male and 1 female) were imaged using *in vivo* three-photon microscopy. A representative example is shown in **Fig. 2k-n**. A chronic cranial window was implanted over the sensory cortex. Briefly, under 0.5–2% isoflurane anesthesia, a head restraint bar was attached to the skull using C & B Metabond (Parkell) and a circular craniotomy 5-mm in diameter was opened over the left visual cortex at coordinates 2.7 mm lateral, 3 mm posterior to bregma. A durotomy was performed and the craniotomy was sealed with a plug consisting of a stack of three #1 coverslips (two round 5 mm coverslips and one round 6 mm coverslip), attached to each other using optical adhesive, and attached to the skull with Metabond.

Imaging was performed on a three-photon microscope built around a Coherent Monaco/Opera-F laser source (≤2 μJ, 50 fs pulses at 1 MHz; Coherent Inc.). The microscope head was based on a Movable Objective Microscope with three-dimensional (3D) objective translation, provided by Sutter Instrument. The scan lens and tube lens were a Thorlabs SL50-3P and Thorlabs TTL200MP, respectively, to enhance transmission at 1300 nm. An Olympus 25-X, 1.05 NA water-immersion objective lens (XLPlan N; 75% transmission at 1300 nm) was used, and image acquisition was controlled by ScanImage (MBF Bioscience) with acquisition gating for low repetition rate lasers. Animals were imaged in the awake state. To label the vasculature, prior to imaging mice were injected with 100 μL of 5% (w/v in sterile saline) 2 MDa dextran-FITC through the retro-orbital vein under a brief period of isoflurane anesthesia. Emitted green fluorescence was separated from incoming excitation light by a primary dichroic beam splitter (FF735-Di02, Semrock), then filtered by a bandpass filter (ET525-70m-2p, Chroma). Third harmonic generated (3HG) signal produced by blood vessels and myelinated axons was detected using a secondary dichroic beam splitter (Di02-R488, Semrock) and filtered by a bandpass filter (ET434/32m, Chroma) into a separate blue channel which allowed us to clearly discern the gray-to-white matter boundary.^57^

A separate batch of adult C57BL/6 mice (n = 5, 3 male and 2 female) was used for *in vivo* three-photon capillary ablation experiments (**Fig. 6**). After chronic cranial window implantation, using the procedure described above, mice were imaged to identify and map microvascular networks in layer 4 and layer 6/CC to be studied in greater detail. One week later, Day 0 baseline imaging was performed, when z-stacks of the chosen networks were collected to obtain baseline vessel diameter data and line scans for determination of baseline RBC flux. This was immediately followed by laser irradiation to induce pre-convergence capillary ablation or to perform sham line scans. Ablative line scans were performed by placing the line scan path of laser irradiation on the vessel lumen. Line scan laser power used to achieve ablation was 35-48 mW in layer 4 and 61-77 mW in layer 6/CC, with a 0.5 ms line scan period and 10,000 repetitions. Sham line scans were performed with the same settings for the corresponding layer and animal, but the line scan path was placed on the parenchyma in the immediate vicinity of the vessel lumen. Post-irradiation imaging was performed at day 3, 7 and 21. During each post-irradiation imaging day, line scans were performed to determine blood flow in the same population of vessels, while z-stacks of analyzed networks were collected at day 21 for measurement of vessel diameter. In each session, animals were imaged in the awake state. The vasculature was labeled by 2 MDa dextran-FITC, as described above.

### Tissue clearing and 3D immunolabeling for light-sheet imaging

Whole brain 3D light sheet fluorescence microscopy imaging was conducted at the Pennsylvania State University (PSU) and approved by the Institutional Animal Care and Use Committee at PSU. Four (2 male and 2 female) 2-month-old (young) and four (2 male and 2 female) 24-month-old (old) C57BL/6J mice were used in the study (**Extended Fig. 1, 3**). The iDISCO+ tissue clearing protocol was used with modifications.^10,51^ Brain samples were delipidated in SBiP buffer, consisting of ice-cold water, 50mM Na2HPO4, 4% SDS, 2-methyl-2-butanol and 2-propanol. This buffer becomes activated at room temperature and was therefore made and stored at 4°C prior to use. Each sample was submerged in 10 mL of SBiP buffer, rotated at room temperature with buffer changes at 3 and 6 hours, followed by incubation with fresh SBiP buffer overnight. For adequate delipidation, particularly with aged samples, each brain was then washed with SBiP for a total of 6 days, with daily buffer changes. After delipidation, brain samples were washed with B1n buffer, which consists of 0.1% Triton X-100, 1g of glycine, 0.01% 10N NaOH and 20% NaN_3_. Brain samples were washed with 10 mL of B1n buffer at room temperature for 2 days.

To begin immunolabeling, brains were rinsed 3 times for 1 hour each with PTwH buffer, consisting of 1X PBS, 0.2% Tween-20, 10mg heparin, and 2g of NaN_3_. For primary antibody incubations, antibodies were diluted in antibody solution consisting of PTwH buffer with 5% DMSO and 3% normal donkey serum. Antibodies to α-smooth muscle actin (α-SMA) (Rabbit anti-α-SMA, Abcam, cat: ab5694, dilution 1:1000) and transgelin (Sm22)(Rabbit anti-Sm22 Abcam; ab14106, dilution 1:1500) were combined to label the artery wall, as previously described.^10^ Pan-vascular labeling was achieved through staining with DyLight-594 labeled Lycopersicon Esculentum (Tomato) Lectin (Vector labs, cat. no.: DL-1177-1), which was added to both primary and secondary incubations at 1:100 concentration. Pericytes were labeled by combining PDGFRβ (Goat anti-PDGFRβ, R&D Systems; AF1042, dilution: 1:100) and Mouse Aminopeptidase N/CD13 (Goat anti-CD13, R&D Systems; AF2335, dilution: 1:100). Primary antibodies were incubated for 10 days at 37°C. Following primary incubation, PTwH buffer was changed 4-5 times for each sample over the course of 24 hours. A fresh antibody solution was used to dilute all secondary antibodies to a concentration of 1:500. For secondary antibodies, Alexa Fluor® 488-AffiniPure Fab Fragment Donkey Anti-Rabbit IgG (H+L) (Jackson ImmunoResearch laboratories; 711-547-003) was used to detect artery staining and Alexa Fluor® 647-AffiniPure Fab Fragment Donkey Anti-Goat IgG (H+L) (Jackson ImmunoResearch laboratories; 705-607-003) was utilized to detect pericyte staining. After secondary incubation for 10 days at 37°C, brains were washed 4-5 times in PTwH buffer for 24 hours. Brain samples were then dehydrated in a series of methanol dilutions in water (1-hour washes in 20%, 40%, 60%, 80% and 100%). An additional wash of 100% methanol was conducted overnight to remove any remaining water. The next day, brains were incubated in 66% dichloromethane/33% methanol for 3 hours and subsequently incubated in 100% dichloromethane twice for at least 15 minutes each. Brains were equilibrated in dibenzyl ether for at least two days before transitioning to ethyl cinnamate one day prior to imaging.

### Light-sheet imaging

A SmartSPIM light-sheet fluorescence microscope (LifeCanvas Technologies) was used to image cleared and stained mouse brains. Brains were supported in the custom sample holder by standardized pieces of dehydrated agarose consisting of 1% agarose in 1X TAE buffer. The sample holder arm was then submerged in ethyl cinnamate for imaging. We used a 3.6X objective (LifeCanvas, 0.2 NA, 12 mm working distance, 1.8 μm lateral resolution) and three lasers (488nm, 560nm, 642nm wavelengths) with a 2 μm step size. Acquired data was stitched using custom Matlab codes adapted from Wobbly Stitcher.^10,51^

### Analysis of light sheet imaging data

For the analysis of the abundance and penetration depth of cortical penetrating vessels, a 2 mm (x) by 2 mm (y) by 1.2 mm (z) ROI centered on the primary somatosensory cortex was cropped from whole brain light sheet data sets of 4 adult mice (**Extended Fig. 1c-e**). The pial penetration points of all penetrating vessels subtypes (penetrating arteriole, PCVs and other ascending venules) were identified within the ROI and the distance to vessel ending points within the tissue was recorded. For penetrating arterioles, we defined the ending point as the deepest region where α-SMA/Sm22 staining was still discernible. For ascending venules and PCVs, the ending point was defined as the region where the main trunk ended and ramified into many smaller branches, as visualized by lectin staining.

PCVs were clearly distinguished from other ascending venules by their large diameter, weak α-SMA/Sm22 staining of the vessel wall and specific branching pattern in the white matter of the CC. Image cropping and analysis was performed using the Fiji software.

For the analysis of vascular length density and pericyte density across different cortical layers, a 540 µm (x) by 540 µm (y) by 1000 µm (z) ROI centered on the primary somatosensory cortex was cropped from whole brain light sheet data sets of 4 adult and 4 aged mice (**Extended Fig. 3e, f**). The ROI volumes were then further divided into 200 µm thick (z) subsections (**Extended Fig. 3g, h**). The 3D segmentation of all vessels labeled with lectin, as well as annotation of all capillary PDGFRβ+CD13 labeled pericytes was performed using the Filament Tracer module of Imaris software (**Extended Fig. 3i, j**). Only mesh and thin strand pericytes on small diameter vessels, characterized by the “bump on the log” morphology, were included in the analysis. Vascular length density (total vascular length per mm^3^) and pericyte density (total number of pericyte cell bodies) were measured for each subsection. Then the normalized pericyte density was calculated by dividing the pericyte and vascular length density values for each subsection.

### In silico modeling of microvascular networks

Both microvascular networks used for *in silico* modeling in this work (MVN1 and MVN2) were acquired from the vibrissa primary sensory cortex of C57/BL6 male mice by Blinder et al.^58^ In these segmented and vectorized networks, each vessel is represented by an edge with a given length and diameter, and edges are connected at bifurcations (graph vertices). MVN1 and MVN2 are embedded in a tissue volume of ∼1.6 mm^3^ and ∼2.2 mm^3^ and contain ∼12,100 and ∼19,300 vessels, respectively. The vessels are labeled as pial arteries, penetrating arterioles, capillaries, ascending venules, and pial veins. Moreover, for the penetrating trees, we differ between the main perforator and off-shooting vessels. The predominant vascular length is composed of capillaries (96% and 94% of total vascular length in MVN1 and MVN2, respectively). As staining of mural cells is not available for the microvascular network reconstructions, the vessel identities are based on topology and diameter.^58^

### Blood flow modeling with discrete RBC tracking

The numerical model to simulate blood flow with discrete red blood cell (RBC) tracking in realistic microvascular networks has been described in Schmid et al.^30^ Below, we briefly summarize the key aspects of the modeling approach, which have been detailed in prior studies.^30,59^ The modeling approach is based on the small Reynolds number (Re < 1.0 for all vessels) in the microvasculature. As such the flow is laminar and mostly even in the Stokes regime, which allows describing the flow in individual vessels by Poiseuille’s law. The flow rate *q_ij_* in vessel *ij* between node *i* and *j* is computed by:

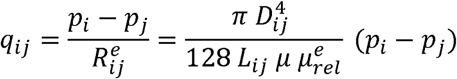

where *D_ij_* and *L_ij_* are the vessel diameter and the length and *p_i_* and *p_j_* are the pressure at node *i* and *j*, respectively. μ is the dynamic plasma viscosity and 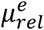 is the relative effective viscosity, which accounts for the presence of RBCs and is computed as a function of local hematocrit and vessel diameter as described in Pries et al. (*in vitro* formulation).^31^

To compute the local hematocrit, we track individual RBCs through the MVNs. We accounted for the Fahraeus effect (RBC velocity is larger than bulk flow velocity) and the phase separation (RBCs partition with a different ratio than the bulk flow at divergent bifurcations).^31^ At divergent bifurcations with a diameter >10 µm the phase separation is described by empirical equations.^31^ At smaller diameters, RBCs transit is single file and consequently, the RBC motion can be approximated by assuming that the RBCs follow the path of the largest pressure force.^30,60^ It is important to note that the RBC distribution and the flow field fluctuate in time and that the RBC distribution impacts the local flow field.^27,59^ In this study, we use the time-averaged flow field to compare changes in response to cortical layer-specific alterations in capillary diameters. The time-averaged flow field is computed by averaging over 14-20s. The exact averaging interval depends on the vessel-specific turn-over time (vessel length/RBC velocity) and ensures that 90% of all vessels are perfused at least six times.

Two configurations were considered regarding the boundary conditions for in- and outflow vertices below the cortical surface. In configuration 1 (“*open*”), fixed pressure values are assigned at all in- and outflows. These values are kept constant for all scenarios mimicking vascular alterations during aging (see below). Further details on assigning suitable pressure boundary conditions can be found in our prior studies.^30^ In configuration 2 (“*trimmed*”) we assume that no blood flow enters/leaves the simulation domain below the cortical surface and that all flow enters/leaves via the pial vessels at the cortical surface. This approach is equivalent to assigning no-flow boundary conditions at in- and outflows below the cortical surface or to removing all in-and outflows below the cortical surface. Configuration 1 (“*open*”) has the advantage that it does not under-predict perfusion, as is the case for trimmed networks.^61^ On the other hand, the number of boundary nodes is significantly small for configuration 2 (“*trimmed*”). As such, this setup is less sensitive to the assigned boundary values, which is especially desirable for relatively large microvascular alterations. Thus, for the current study, we decided to consider both open and trimmed boundary conditions for a total of four configurations: MVN1-open, MVN1-trimmed, MVN2-open, and MVN2-trimmed. The inflow hematocrit of 0.3 is constant for all simulations.

### Mimicking vascular alterations observed in aged mice in silico

We mimicked two age-related vascular alterations observed *in vivo* to understand their effects on microvascular perfusion. As we are interested in perfusion changes with respect to cortical depth, each capillary is assigned to one of the five cortical layers (L1, L2/3, L4, L5, L6). This is done by computing the average depth per capillary from the tortuous vessel coordinates and assigning the capillary to the respective layer based on the minimum and maximum depth for each layer (L1: up to 200 µm, L2/3: 200-400 µm, L4: 400-600 µm, L5: 600-800 µm, L6: below 800 µm). Only capillaries within the 5^th^ to 95^th^ percentile of all vessel depths and at least two branches from any inflow and outflow were considered for vasoconstriction and subsequent analyses. This is done to avoid confounding effects with respect to boundary conditions. For all configurations, the 95^th^ percentile is at a depth of at least 999 µm and we have 8,200-15,000 capillaries for analyses.

We focused on capillaries towards the venule end of the capillary bed to mimic changes to pre-convergence capillaries. To assess the position of individual capillaries along the path from arteriole to venule, the median distance to the penetrating arteriole trunk (*medianDistMainDA*) and the ascending venule trunk (*medianDistMainAV*) is calculated for each capillary. Therefore, all paths leading from an individual capillary to the arteriole trunk are recorded. The median of these paths lengths represents the characteristic *medianDistMainDA* per capillary. To mimic vasoconstriction in the anesthetized state (**Supplementary Fig. 16**), all capillaries belonging to layer 6 and with a *medianDistMainDA* > 3 and *medianDistMainAV* > 1 were constricted by 7%. This corresponds to 1168, 948, 1669, and 1610 for MVN1-open, MVN1-trimmed, MVN2-open, and MVN2-trimmed, respectively.

To reduce capillary density in layer 6, evidence from the current study suggests that shorter capillaries tend to regress more frequently during aging (**Supplementary Fig. 22**). Consequently, the maximum length for a regressing capillary was set to 70 µm. Moreover, as before, only capillaries with a *medianDistMainDA* > 3 and *medianDistMainAV* > 1 are eligible for regression. These two criteria (length and position along the capillary path) yielded 788, 627, 1165, and 1131 capillaries potentially affected by regression in MVN1-open, MVN1-trimmed, MVN2-open, and MVN2-trimmed networks. Considering that the total number of capillaries in layer 6 is 1707, 1413, 1789, and 1737 for these networks, it implied that 170, 141, 178, and 173 capillaries needed to be removed to mimic a density reduction by 10%. For each network, there were more capillaries that fit our selection criterion than needed to be removed from the network. Therefore, for each network, we generated 5 cases where candidate capillaries were randomly removed to match the target density reduction of 10%. These 5 cases were then used to compute the average change for a density reduction of 10% per configuration, increasing robustness of blood flow outcomes measured for each network. Capillary regression was modeled by constricting the selected capillaries by 97%. By comparing the average flow rate in the regressed capillaries to all flow rates in the network, we confirmed that the average flow rate in the regressed capillaries is close to zero (<2.5^th^ percentile), effectively removing these capillaries from the perfused network. Microvascular alterations during the awake state were characterized by a 4.5% dilation in Layer 2/3, a 5.1% dilation in Layer 4, and a 14% constriction and 10% reduced density in Layer 6 (**Fig. 5, Supplementary Fig. 14**). The changes were introduced in an equivalent way, as described for the anesthetized case above.

The time-averaged flow field was computed for all four configurations (MVN1-open, MVN1-trimmed, MVN2-open, and MVN2-trimmed) and three scenarios each: 1) capillary vasoconstriction in L6, 2) reduced capillary density in L6 and 3) vasoconstriction and density reduction in L6. Quantities of interest were the flow rate, the RBC velocity, the RBC flux, and the linear density. The flow rate was directly obtained from the pressure drop across the vessel and the effective flow resistance. The RBC velocity is computed by *vf* · *q_ij_* /*A_ij_*, where *A_ij_* is the vessel cross-section and *vf* is the velocity factor accounting for the increased velocity of RBCs in comparison to the bulk flow and which is defined as the ratio of discharge to tube hematocrit.^31^ The discharge hematocrit is calculated in function of the tube hematocrit and the vessel diameter as defined in.^31^ The RBC flux 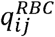 is the product of the flow rate *q_ij_* and the discharge hematocrit *htd* divided by the RBC volume V^RBC^, i.e., 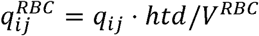 . The RBC volume is 49 fl. As with the *in vivo* experiments, the linear density is computed by dividing the RBC flux by the RBC velocity.

In addition to comparing average perfusion quantities per layer, we computed the integral capillary inflow and outflow per cortical layer (**Fig. 5f,g**). Therefore, all startpoints and endpoints of the capillary bed were identified. A capillary start point is a bifurcation at which the vessel label changes from arterial vessel to capillary. The transition point between capillary and venule vessel marks a capillary endpoint. Capillary start and endpoints were assigned to the different layers based on their cortical depth. To obtain the integral capillary inflow per cortical layer all arterial inflows into all capillary start points of the respective layer are summed (**Fig. 5f**). The equivalent computation is performed on the venule side to compute the integral outflow of the capillary bed (**Fig. 5g**).

There are some limitations of our *in silico* model in its ability to mimic the *in vivo* scenario. For example, the *in silico* model only considers microvascular alterations in an averaged sense. Thus, all capillaries are constricted by a similar magnitude, while *in vivo* a range of constrictions and dilations occurs. Moreover, our model does not account for additional age-related changes such as a reduced systemic hematocrit or vessel wall stiffening. Also, from a topological perspective it should be noted that the employed vascular networks do not contain a full PCV trunk. This forces capillary flow to be drained via shorter venules, which increases the flow resistance of deeper layers and could explain why predicted flow reductions are larger for the *in silico* model than observed *in vivo*. It is not possible mimic all aspects of biological complexity in an *in silico* model. Despite this, the mentioned limitations do not affect the conclusions in the context of how layer-wise alterations in capillaries redistribute blood flow.

### Immunohistology

Thy1-YFP mice, 5-7 month old (n=4) and 24 month old (n=5), were deeply anesthetized with euthasol and transcardially perfused with PBS followed by 4% paraformaldehyde. Brains were dissected, post-fixed overnight, cryoprotected in 30% sucrose with 0.001% sodium azide for 24 h, frozen in OCT and cryosectioned using a Leica cryostat. Brain sections (100 µm) were immuno-stained to detect microglia (rabbit anti-Iba1; Wako Chemicals, 019-19741; dilution: 1:250), astrocytes (mouse anti-GFAP; Santa Cruz, 2E1; dilution: 1:200), or myelin (mouse anti-MBP; Biolegend, SMI 99; dilution: 1:500) for 48 h at 4°C. Following incubation with primary antibodies, sections were incubated with appropriate Alexa Fluor conjugated secondary antibodies (ThermoFisher, dilution: 1:500) with DAPI for 2 h at room temperature. All antibody staining was performed in a solution of 2% TritonX-100, 10% goat serum and 0.1% sodium azide in PBS. Tissue sections were washed 3 times for 5 minutes in PBS after each antibody incubation. Following staining, tissue was mounted onto slides with Fluoromount-G (ThermoFisher; 00-4958-02).

Views of the entire sagittal brain section were obtain using an Olympus VSO120 Slide Scanner. High-resolution confocal images were taken from the CC using a Zeiss 710 LSM confocal microscope, and these images were used for analyses of data in **Supplementary Fig. 18**. Microglia/Iba1 cell density was determined within the CC underlying the somatosensory cortex by counting the number of Iba1+ cells divided by the total area using FIJI. Fluorescence intensity of GFAP, MBP, and Thy1-YFP was determined by outlining the CC and measuring the mean signal intensity using FIJI. All samples were stained and imaged within the same batch. Exposure times and laser powers were kept consistent between all samples.

A separate batch of adult (5-7 month old, n = 4), mid-aged (16-18 month old, n = 6) and aged (22-24 month old, n = 6) mice was injected with 80 mg/kg pimonidazole HCl (Hypoxyprobe, HP3-1000Kit) via the retro-orbital vein under isoflurane anesthesia. Under deep anesthesia, the mice were then transcardially perfused with PBS at 90 min after Hypoxyprobe administration. Brains were isolated and hemisected at the midline. One hemi-section was used for transcriptional analyses (see below), and the other was immersed in 4% PFA for 24 h and then transferred to 30% (w/v) sucrose for cryoprotection. Cryoprotected brain hemi-sections were embedded in OCT (Fisher Healthcare, 23-730-571) and cut into 100 μm sagittal free-floating sections on a Leica cryostat. Sections were then incubated with primary antibodies in antibody solution (10% goat serum, 0.02% triton-x 100, 0.02% sodium azide in PBS) for 48 h at 4°C with gentle shaking. Specific antibodies were used to immuno-stain for astrocytes (mouse anti-GFAP antibody; Sigma-Aldrich, C9205-2ML; dilution: 1:100), microglia (rabbit anti-Iba1 antibody; Wako Chemicals, 019-19741; dilution: 1:100), myelin (rat anti-Myelin Basic Protein antibody; Abcam, ab7349; dilution: 1:100), VCAM1 (rat anti-CD106 antibody; Biolegend, 105701; dilution 1:200), and Hypoxyprobe (rabbit anti-Hypoxyprobe antibody; Hypoxyprobe, PAB2627; dilution: 1:100), while the vasculature was labeled with Lycopersicon Esculentum Tomato Lectin (Vector labs, DL-1177-1 for DyLight®-594 labeled Lectin, DL-1178-1, DyLight® 649). After washing away unbound primary antibodies with PBS, sections were incubated with fluorescently labeled secondary antibodies (Goat anti-Rabbit IgG (H+L) Alexa Fluor™647, Goat anti-Rabbit IgG (H+L) Alexa Fluor™488, Goat anti-Rat IgG (H+L) Alexa Fluor™594) for 2 h at room temperature. Before mounting, the sections were incubated with 4′,6-diamidino-2-phenylindole (DAPI) to label nuclei.

Stained brain sections were imaged using an Evident Scientific APX100 microscope. All image processing and quantification was performed in ImageJ (version 1.54f). Three non-overlapping ROIs (300 x 150 x 24 (x, y, z) μm) from somatosensory midcortex and 3 non overlapping ROIs (150 x 150 x 24 (x, y, z) μm) from somatosensory layer 6/CC were obtained from adult, mid-aged and aged sections (4 mice/group). For analysis of microglia and astrocytes, Iba-1- and GFAP-positive area in each ROI was measured and averaged. Specifically, background was subtracted (rolling ball radius=50, two times) in maximally projected Z-stacks of 16-bit exported images. After despeckling, and thresholding (triangle thresholding) images were converted to a mask. The binary images were eroded accordingly to remove non-cellular signal. The binarized signal was measured and expressed as the percentage of the total ROI area. For analysis of MBP, VCAM1 and Hypoxyprobe staining, signal intensity was measured by recording the mean gray value of each ROI. For vessel density analysis, ROIs that included the mid-cortex and layer 6/CC of the somatosensory cortex were acquired from lectin-labeled sections. Vessel segmentation was performed in Imaris software version 7.7.2 (Bitplane, Oxford Instruments) from maximally projected Z-stacks, and conducted in a semi-automated fashion to ensure detection of only vasculature and not lectin-labeled cells, such as microglia.

### RNA transcription analyses of adult, mid-aged, and aged mice

Mice were anesthetized and transcardially perfused with PBS. The brains were dissected and hemisected at the midline, with the hemisected brains placed on their sides in ice-cold PBS to visualize internal structures. The cortex and underlying CC, above the hippocampus, was extracted. Then, the upper and lower cortices were separated using a scalpel, with lower cortex samples retaining the CC. The dissected tissue was immediately placed on dry ice to preserve RNA quality. RNA was extracted using the Qiagen RNeasy Mini Kit (cat no. 74104), and cDNA was synthesized with the BioRad iScript cDNA Synthesis Kit (cat no. 1708890). RNA and cDNA quantity and quality were assessed using a Nanodrop spectrophotometer. Expression of hypoxia-driven genes, including *slc2a1*, *ldha*, *pdk1*, and *vegfa*, was analyzed using the following primer sequences: *slc2a1* forward: TCAGGCGGAAGCTAGGAAC, *slc2a1* reverse: GGAGGGAAACATGCAGTCATC; *ldha* forward: AGCAGGTGGTTGAGAGTGCT, *ldha* reverse: GGCCTCTTCCTCAGAAGTCA; *pdk1* forward: CCCCGATTCAGGTTCACG, *pdk1* reverse: CCCGGTCACTCATCTTCACA; *vegfa* forward: CAGGCTGCTGTAACGATGAA, *vegfa* reverse: TTTGACCCTTTCCCTTTCCT. *Ppia* was used as a housekeeping gene with the following primer sequences: *ppia* forward: CCCTGGCACATGAATCCTGG, *ppia* reverse: GAGCTGTTTGCAGACAAAGTTC. Delta-delta Ct analysis was employed to assess changes in the expression of hypoxia-driven genes in the upper and lower cortices of adult, mid-aged, and aged mice.

### Unilateral Common Carotid Artery Stenosis (UCCAS) model experiment

A batch of adult (5-7 months old) C57BL/6 mice (n = 7, 3 male and 4 female) was used for UCCAS model studies. Each animal was first implanted with a chronic cranial window centered over the somatosensory cortex, as described above. After a 3-week recovery period, animals underwent baseline *in vivo* deep two-photon imaging to identify layer 4 and layer 6/CC PCV networks for analysis, and to perform line scans for baseline RBC flux measurements in pre-convergence capillaries. Three days later, mild unilateral cerebral hypoperfusion was induced by microcoil placement around the left common carotid artery (ipsilateral to imaged brain hemisphere). Under isoflurane anesthesia, mice were placed in the supine position on a feedback-regulated heat pad (FHC Inc.) under the surgical microscope. A surgical plane of anesthesia was maintained at ∼1.5% MAC isoflurane, delivered with medical air (20–22% oxygen and 78% nitrogen, moisturized by bubbling through water; AirGas Inc.). The ventral neck skin was cleaned with alternating swabs of 70% ethanol and 10% povidone-iodine solution. A 1.5 cm incision was made in the midline of the ventral neck. After resecting subcutaneous fat and salivary glands from the surgical field, the common carotid area, superior vena cava and vagal nerve was dissected from the connective tissue surrounding them. Two pieces of silk suture (4-0) were placed around the proximal and distal end of the common carotid artery for ease of manipulation. A microcoil with inner diameter 0.20 mm (Sawane Spring Co. Ltd.; 0.08*0.20*0.5*2.5 SWPA) was intertwined with the left common carotid artery to encase the artery within the center of the microcoil.^62^ The neck incision was sutured with 4-0 monofilament (ETHILON®, 662G). Post-surgery imaging was performed at day 3, 7, 14 and 21 to determine blood flow in the same population of pre-convergence capillaries on which baseline imaging was performed. In each imaging session, animals were imaged under 1.5% isoflurane anesthesia. To label the vasculature prior to imaging, mice were injected with 100 μL of 5% (w/v in sterile saline) 2 MDa dextran-Alexa 680 through the retro-orbital vein.

On post-surgery day 24, mice received 80 mg/kg pimonidazole HCl (Hypoxyprobe, HP3-1000Kit) via retroorbital injection under isoflurane anesthesia. After 90 minutes, the mice were then perfused transcardially with PBS, followed by 4% PFA under the overdose of Euthasol® (Patterson Veterinary, 07-805-9296). Brains were removed and immersed in 4% PFA for 24 h and then transferred to 30% (w/v) sucrose for cryoprotection. Cryoprotected brains were embedded using OCT (Fisher Healthcare, 23-730-571), and 100 μm coronal free-floating sections were obtained to provide comparison of brain hemispheres ipsilateral and contralateral to the stenosed carotid artery. The contralateral hemisphere was used as internal control for histological assessment. We immuno-stained for microglia (rabbit anti-Iba1 antibody; Wako Chemicals, 019-19741; dilution: 1:100), astrocytes (Cy3-conjugate mouse anti-GFAP antibody; Sigma-Aldrich, C9205-2ML; dilution: 1:100), myelin (rat anti-Myelin Basic Protein antibody; Abcam, ab7349; dilution: 1:100), and Hypoxyprobe (rabbit anti-Hypoxyprobe antibody; Hypoxyprobe, PAB2627; dilution: 1:100). Pan-vascular staining was achieved via Lycopersicon Esculentum Tomato Lectin (Vector labs, DL-1177-1 for DyLight® 594; DL-1178-1, DyLight® 649 labeled Lectin). Free-floating sections were incubated in above mentioned primary antibodies in antibody solution (10% goat serum, 0.02% Triton-X 100, 0.02% sodium azide in PBS) for 48 h at 4°C on gentle-shaking. After washing the primary antibodies, the tissues were incubated with fluorescently labeled secondary antibodies (Goat anti-Rabbit IgG (H+L) Alexa Fluor™647; Goat anti-Rabbit IgG (H+L) Alexa Fluor™488; Goat anti-Rat IgG (H+L) Alexa Fluor™594) for 2 h at room temperature. Before mounting, the sections were incubated with DAPI to label nuclei.

Stained brain sections were imaged using an Evident Scientific APEX100 microscope. All image processing and quantification was performed in ImageJ (version 1.54f). For each brain section, 3 nonoverlapping ROIs (150 x 150 x 24 (x, y, z) μm) were placed in ipsilateral mid-cortex, contralateral mid-cortex, ipsilateral Layer 6/CC and contralateral Layer 6/CC of the somatosensory cortex (12 ROIs per section). Analysis of immunostaining was performed as described above for the examinations of tissue changes in adult, mid-aged and aged mice.

### Statistics

All statistical analyses for *in vivo* and light-sheet data were performed with SPSS software. Analysis of all parameters was performed using a repeated-measures analysis of variance (ANOVA) model or a linear mixed-effects model, with Age (adult vs. aged) and Layer (2/3 vs. 4 vs. 6/CC) set as independent factors, and Animal as a within group factor. Therefore, data from each animal was assigned to the corresponding age group, and further subdivided into the 3 analyzed layers. This strategy considers the nested nature of the data; that is, that multiple vessels come from one animal and are thus not independent of one another. Strength and significance of correlation between analyzed parameters was determined using the Pearson correlation analysis in the GraphPad Prism version 9 software. All graphs were made using the GraphPad Prism version 9 software.

Analysis of age-dependent histological changes and UCCAS studies was performed using a two-way repeated measures ANOVA model in GraphPad Prism version 9 software. For histology, two independent factors within the model were Age (adult vs. mid-aged vs. aged) and Layer (4 vs. 6/CC) for aging sections with fluorescent intensity or percentage of positive area of the immunohistochemical marker as dependent variable. For UCCAS studies, Hemisphere (ipsilateral vs. contralateral to the stenosis) and Layer (4 vs. 6/CC) were determined as independent variables. *Post-hoc* tests were performed using Holm-Šídák test to further explore significant interactions. All statistical tests were performed at an alpha level of 0.05.

### Animal and data exclusion criteria

Out of 12 adult chronic cranial windows, 1 was damaged before the line scan data for layer 6/CC could be obtained. Out of 12 aged windows 1 was damaged before the line scan data for layer 6/CC could be obtained, while for another only the line scan data for layer 6/CC was obtained. This resulted in 12 complete data sets for layers 2/3 and 4, and 11 complete data sets for layer 6/CC in the adult group, as well as 11 complete data sets for all layers in the aged group.

For analysis of structural and functional parameters of vascular segments, if the signal quality did not allow for reliable analysis of lumen diameter, RBC flux or blood flow velocity these parameters were excluded from the analysis for the corresponding vessel segment, while the length and tortuosity data were included. Correlation analyses were performed only on the population of vessels where all analyzed structural and functional parameters could be obtained.

## DATA AVAILABILITY

Raw image files are stored on servers at Seattle Children’s Research Institute owing to their large size. These raw data can be provided by the corresponding author upon request. All source data, such as extracted metrics of vascular structure and blood flow used in figures, are provided as supplementary materials.

## CODE AVAILABILITY

All details regarding the simulation code used to produce the *in silico* results are available in the original publication of the *in silico* model. Further details and explanations are available from F.S. upon request.

## AUTHOR CONTRIBUTION

Conceptualization and experimental design by SS, FS, GG, FA, JW, and AYS. Experiments were conducted by SS, FA, GG, NW, KT, SKB, MJS, and HCB. Data analysis and statistics were performed by SS, GG, NW, MJS and SKB. *In silico* studies and analyses were performed by FS. The manuscript was written by SS and AYS with contributions from all authors.

## COMPETING INTERESTS STATEMENT

The authors declare no competing interests.

## ACKNOWLEDGMENTS

This work was supported by grants from the NIH/NIA (R01AG062738, R21AG069375, RF1AG077731, R01AG081840) and a pilot award from the Albert Institute for White Matter Research. FS received funding from the Swiss National Science Foundation (Grant No. 202192). SKB was supported by fellowships from the NIH/NINDS (F32NS117649) and NIH/NIA (K99AG080034). MJS was supported by a Diversity supplement for a grant from the NIH/NIA (R01AG062738-03S1). We also thank Tiago Figueiredo for creating the artwork used in **Fig. 2e,f**. We would also like to thank Juliane Gust, Yuandong Li, and Catherine Foster for their helpful comments and suggestions during manuscript preparation.

## FIGURE LEGENDS

**Extended Figure 1. Principal cortical venules in relation to other penetrating vessel types in mouse sensory cortex. a.** Light-sheet imaging data showing dorsal view of a mouse brain with arterioles labeled (α-SMA and Sm22). The dotted yellow square marks the region of interest (ROI), centered on the primary somatosensory cortex, used for analysis of the number and penetration depths of cortical penetrating vessels. **b.** Maximum projection images showing the side view of the ROI defined by the white dotted lines in panel (a), with labeling of cortical arterioles by Acta-2 and Sm22 (upper panel) and all blood vessels by lectin (lower panel). Pial penetration points of arterioles and venules are marked with arrows. **c.** Side projection of analyzed volume with colored circles marking the intracortical termination points of different penetrating vessel types. The inset on the right shows an enlarged volume denoted by a white square, with a venule termination point marked by a cyan circle. **d.** Plot of the average number of penetrating vessel types within the ROIs. N=4 adult mice. Data shown as mean ± SD. **e.** Plot of the intracortical termination depth of different penetrating vessel types. Each dot corresponds to a single vessel. Gray dotted line shows approximate depth of gray-white matter transition. N=218, 422, and 13 vessels for arterioles, other venules, and PCVs, respectively, collected over 4 adult mice.

**Extended Figure 2. Penetrating arteriole and arteriole-capillary transition zones of adult and aged mice are similar in diameter. a.** Maximally projected image in Layer 6/CC showing a top-down view of the vascular bed surrounding the main trunk of a PCV in an adult mouse. Red arrow marks a penetrating arteriole trunk. **b.** Inset showing a magnified view around the penetrating arteriole with the first three branch orders of vessel offshoots, referred to as arteriole-capillary transition zone. **c.** Average diameter of the penetrating arteriole trunk throughout cortical depth in adult and aged mice. Each dot corresponds to an average diameter of arteriole trunks at the corresponding depth in a single animal. One-way ANOVA with Holm-Sidak post-hoc test. Adult, n=13 mice, with 3-4 vessels per cortical depth per animal; Aged, n=12 mice, with 3-4 vessels per cortical depth per animal. Data shown as mean ± SD. **d-f.** Average diameter of vessels in the arteriole-capillary transition zone of adult and aged mice in Layer 2/3 (d), Layer 4 (e), Layer 6/CC (f). Each dot corresponds to an average diameter of vessels of the corresponding branch order in a single animal. One-way ANOVA with Holm-Sidak post-hoc test. Adult, n=13 mice (n=12 for Layer 6/CC); Aged, n=12 mice; branch order 1, n=3-4 vessels per layer per animal; branch order 2, n=6-8 vessels per layer per animal; branch order 3, n=12-16 vessels per layer per animal. Data shown as mean ± SD.

**Extended Figure 3. Reduction in vascular length and pericyte cell density in layer 6/CC of aged mice**. **a.** Maximum intensity projection of the dorsal half of a whole mouse brain with arterioles labeled using α-SMA + Sm22 immunostaining imaged by light-sheet microscopy. **b-d.** Top-down views of the 540 x 540 x 1000 µm (x, y, z) regions of interest (ROI) centered on the somatosensory cortex marked with a dotted yellow square in (a), with arterioles labeled using α- SMA + Sm22 antibodies (b), all vessels labeled using fluorescent lectin (c), and mural cells labeled using the combination of anti-PDGFRβ and anti-CD13 antibodies (d). **e,f.** 3D side views from Imaris showing lectin (e) and mural cell (f) staining throughout the entire cortical depth. **g, h.** Examples of 200 µm thick ROIs from (e) and (f) (marked with dotted yellow lines) with reconstructions of all capillary segments (g, gray lines)) and annotated capillary pericyte cell somata (h, blue circles). **i.** Top-down view of vessels (cyan) and mural cells (red). **j.** Magnified regions from (i) (marked with yellow squares) with annotated examples of capillary pericytes exhibiting the “bump on the log” morphology (marked with yellow arrows). **k.** Vascular length density in analyzed cortical subsections of adult and aged mice. Unpaired t-tests: 0-200 µm, p=0.385; 201-400 µm, p=0.054; 401-600 µm, p<0.001; 601-800 µm, p=0.003; 801-1000 µm, p<0.001. **l.** Pericyte density in analyzed cortical subsections of adult and aged mice. Unpaired t-tests: 0-200 µm, p=0.719; 201-400 µm, p=0.201; 401-600 µm, p=0.042; 601-800 µm, p=0.134; 801-1000 µm, p=0.004. **m.** Pericyte density in analyzed cortical subsections of adult and aged mice. Unpaired t-tests: 0-200 µm, p=0.862; 201-400 µm, p=0.804; 401-600 µm, p=0.891; 601-800 µm, p=0.606; 801-1000 µm, p=0.038. For all plots in k-m, n=4 animals per age group. Data shown as mean ± SEM.

## Notes

### Competing Interest Statement

The authors have declared no competing interest.

### Summary of Updates

This revision adds data from new experiments, notably for Figures 6, 7, and 8.

