## Supplementary Figures and Tables for "Impaired capillary-venous drainage contributes to gliosis and demyelination in white matter during aging"

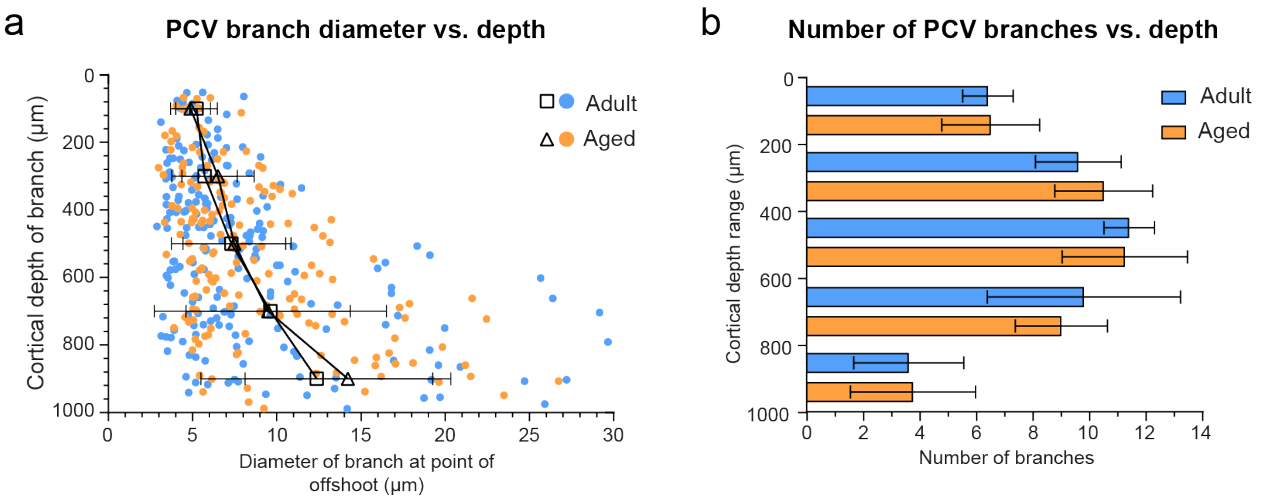


**Supplementary Figure 1. Number and diameter of main PCV branches in adult and aged mice. a.** Plot showing the diameter of PCV branches as a function of cortical depth in adult and aged mice. Each dot corresponds to a single PCV branch. No difference was found between adult and aged mice for average branch diameter in each 200 μm division of intracortical depth. Two way ANOVA analysis $F(1,358)=1.190$; $p=0.276$. Adult, n=5 mice, n=204 branches; Aged, n=4 mice, n=164 branches. Data shown as mean ± SD. **b.** Plot showing the number of PCV branches in each 200 μm division of intracortical depth in adult and aged mice. No difference was found between adult and aged mice for average number of branches. Two way ANOVA analysis $F(1,35)=0.005$; $p=0.946$. Adult, n=5 mice; Aged, n=4 mice. Data shown as mean ± SD.

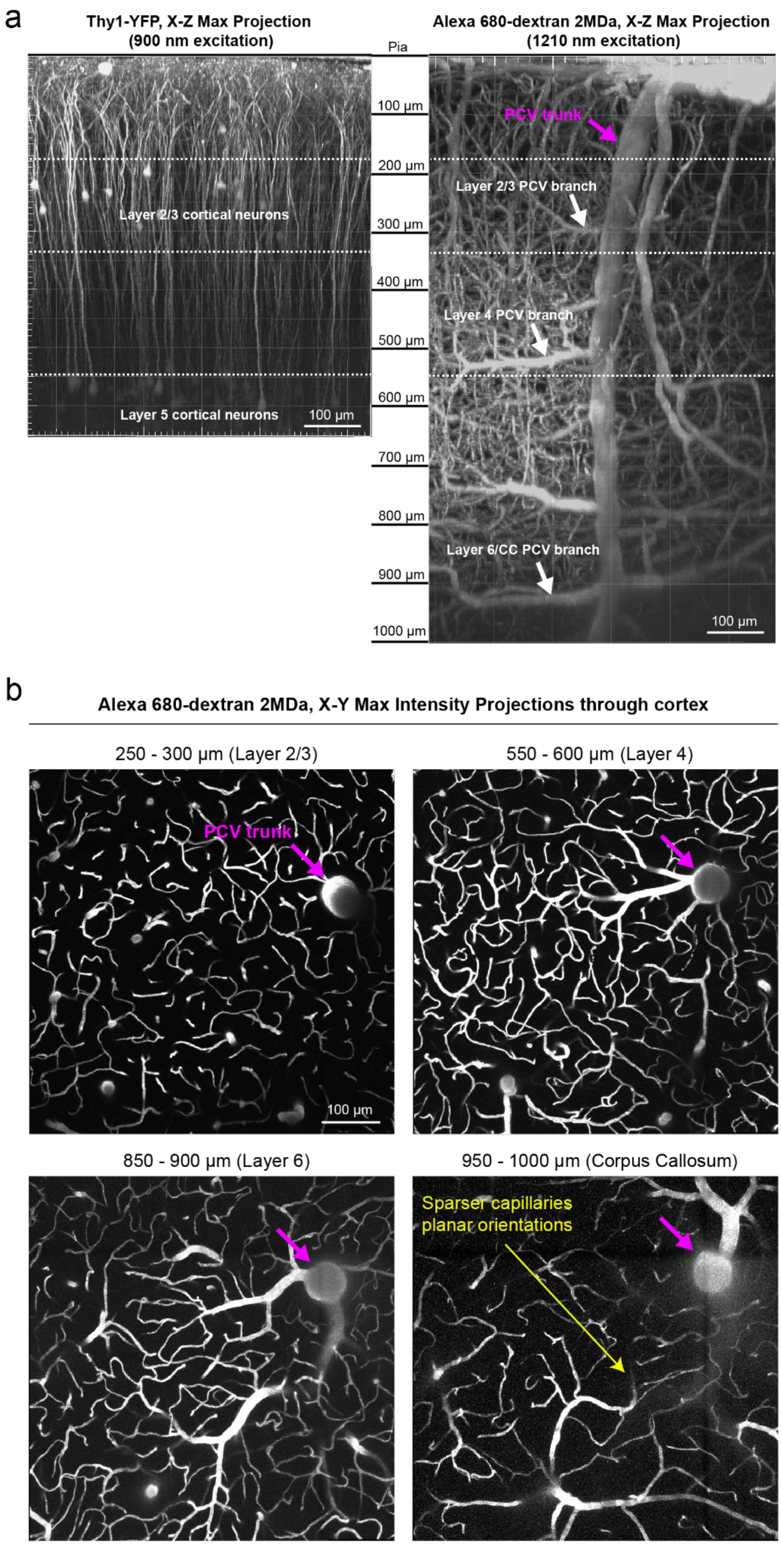


**Supplementary Figure 2. Deep two-photon imaging of Alexa 680-dextran in Thy1-YFP mice to confirm intracortical locations of analyzed PCV branches. a.** Side views of deep two-photon image stacks from an aged mouse showing the endogenous YFP signal in Thy1-YFP mice, imaged at 900 nm excitation (left panel) and the vasculature labeled by i.v. injection of Alexa 680-dextran 2MDa imaged at 1210 nm excitation (right panel). The dotted lines mark the approximate boundaries of layer 2/3 and layer 5, identified by the presence of YFP positive cortical pyramidal neurons. The arrows in the right panel mark the locations of the PCV main trunk and analyzed PCV branches in different cortical layers. **b**. Top-down views of maximum intensity projections of vascular Alexa-680 dextran across different cortical layers in an adult mouse somatosensory cortex. The white arrows point to the location of the PCV trunk.

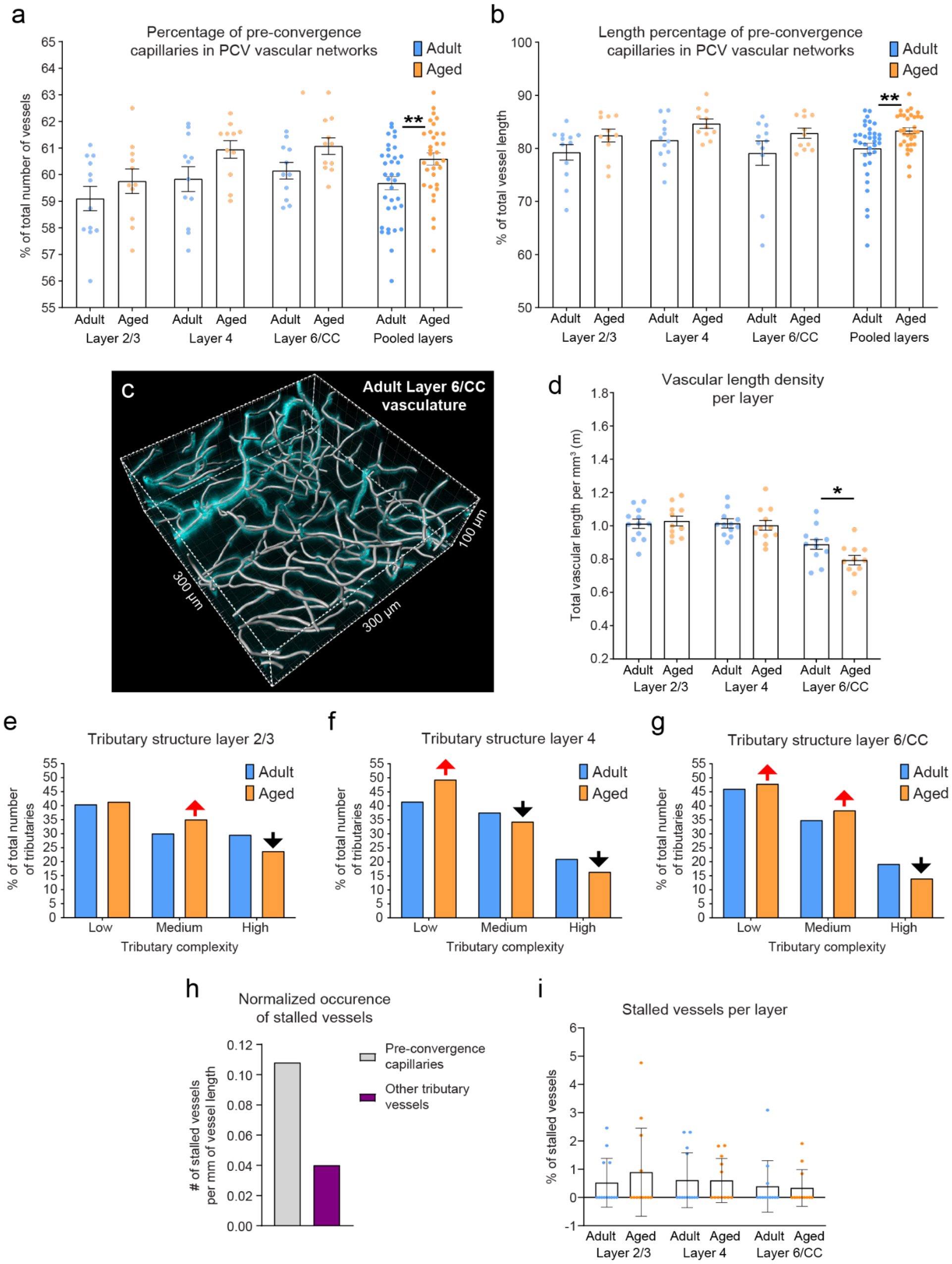


**Supplementary Figure 3. Changes in vascular density and tributary complexity in aged mice. a.** Percent ratio of pre-convergence capillaries in PCV vascular networks of adult and aged mice across analyzed cortical layers. Two-way ANOVA analysis, Age group F(1,62)=7.528; p=0.008. Adult, n=12 mice for layers 2/3 and 4, and 11 mice for layer 6/CC; Aged, n=11 mice for all layers. Data shown as mean ± SEM. **b.** Length percentage ratio of pre-convergence capillaries in PCV vascular networks of adult and aged mice across analyzed cortical layers. Two way ANOVA analysis, Age group F(1,62)=8.591; p=0.005. Adult, n=12 mice for layers 2/3 and 4, and 11 mice for layer 6/CC; Aged, n=11 mice for all layers. Data shown as mean ± SEM. **c.** Imaris reconstruction of all vascular segments within a 300 µm x 300 µm x 100 µm ROI. Two ROIs were analyzed per each layer. The measured vascular lengths in both ROIs were summed and divided by the total analyzed volume for calculation of vascular length density **d.** Vascular length density in adult and aged mice analyzed across cortical layers. Two-way ANOVA analysis with Holm-Sidak's post-hoc comparison testing, Layer F(2,62)=23.971; p<0.001. Layer 2/3, p=0.684. Layer 4, p=0.784. Layer 6/CC, p=0.025. Adult, n=12 mice for layers 2/3 and 4, and 11 mice for layer 6/CC; Aged, n=11 mice for all layers. Data shown as mean ± SEM. **e-g**. Percent ratio of tributaries of different complexities in PCV vascular networks of adult and aged mice in Layer 2/3 (e), Layer 4 (f) and Layer 6/CC (g). Complexity of tributaries are categorized as low (1 or 3 vessel segments), medium (5 or 7 vessels segments), or high (9 or more vessel segments). Red and black arrows point to increase or decrease in the percent ratio of the complexity group in aged mice. **h.** Occurrence of stalled vessels normalized to the total length of the population of corresponding vascular segments. **i.** Plot of the percentage of stalled vessels across cortical layers. Each dot corresponds to a single animal.  Data shown as mean ± SD.

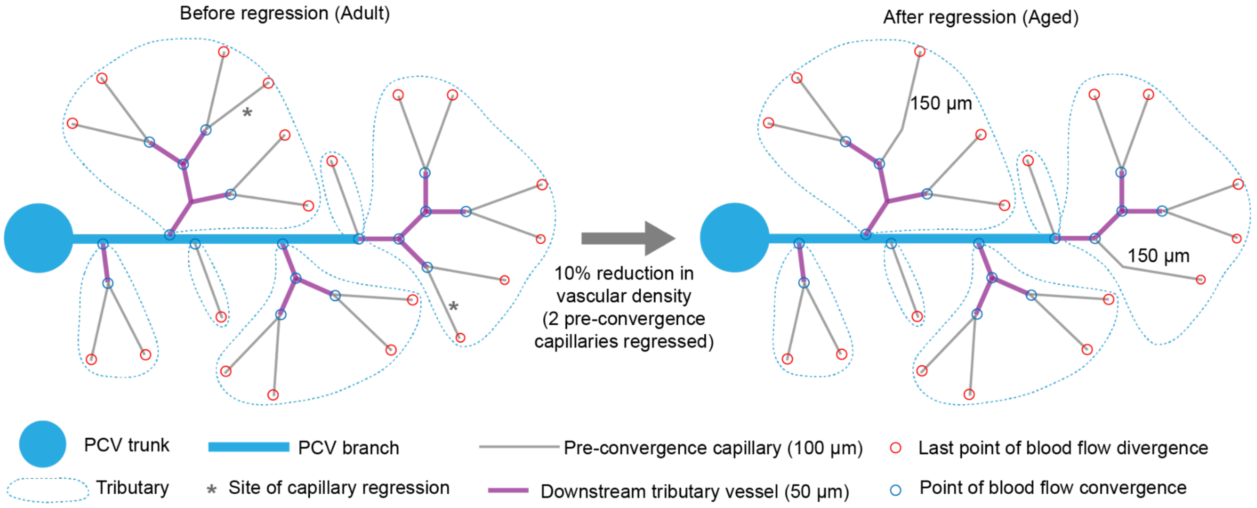


| Analzyed parameter | Before regression (Adult) | After regression (Aged) | Direction of change and empirical data |
|---|---|---|---|
| Number of pre-convergence capillaries | 20 | 18 | ↓ |
| Number of other tributary vessels | 14 | 12 | ↓ |
| Percentage of pre-convergence capillary segments in tributaries | 20/(20+14)*100 = 58.82% | 18/(18+12)*100 = 60.00% | ↑ (Fig. 3c, left) |
| Average number of vessels per tributary | (3+11+1+7+1+11)/6 = 5.67 | (3+9+1+7+1+9)/6 = 5.00 | ↓ (Fig. 3f) |
| Total pre-convergence capillary length | 20*100 µm = 2000 µm | 16*100 µm + 2*150 µm = 1900 µm | ↓ |
| Total downstream tributary vessel length | 14*50 µm = 700 µm | 12*50 µm = 600 µm | ↓ |
| Percentage of total pre-convergence capillary length in tributaries | 2000/(2000+700)*100 = 74.07 % | 1900/(1900+600)*100 = 76.00 % | ↑ (Fig. 3c, right) |
| Average pre-convergence capillary segment length | 2000/20 = 100.00 µm | 1900/18 = 105.56 µm | ↑ (Fig. 3l) |

**Supplementary Figure 4. Analysis of structural changes in PCV vascular networks after pre-convergence capillary regression in a hypothetical model.** Schematic representations of a PCV branch vascular network before (“Adult” state, left-hand side) and after (“Aged” state, right-hand side) pre-convergence capillary regression leading to ~10% reduction in vascular density. The table below summarizes the analyzed structural parameters, the method of calculation, and direction of their change following pre-convergence capillary regression. Figures with empirical data showing the same direction of change listed on the right side.

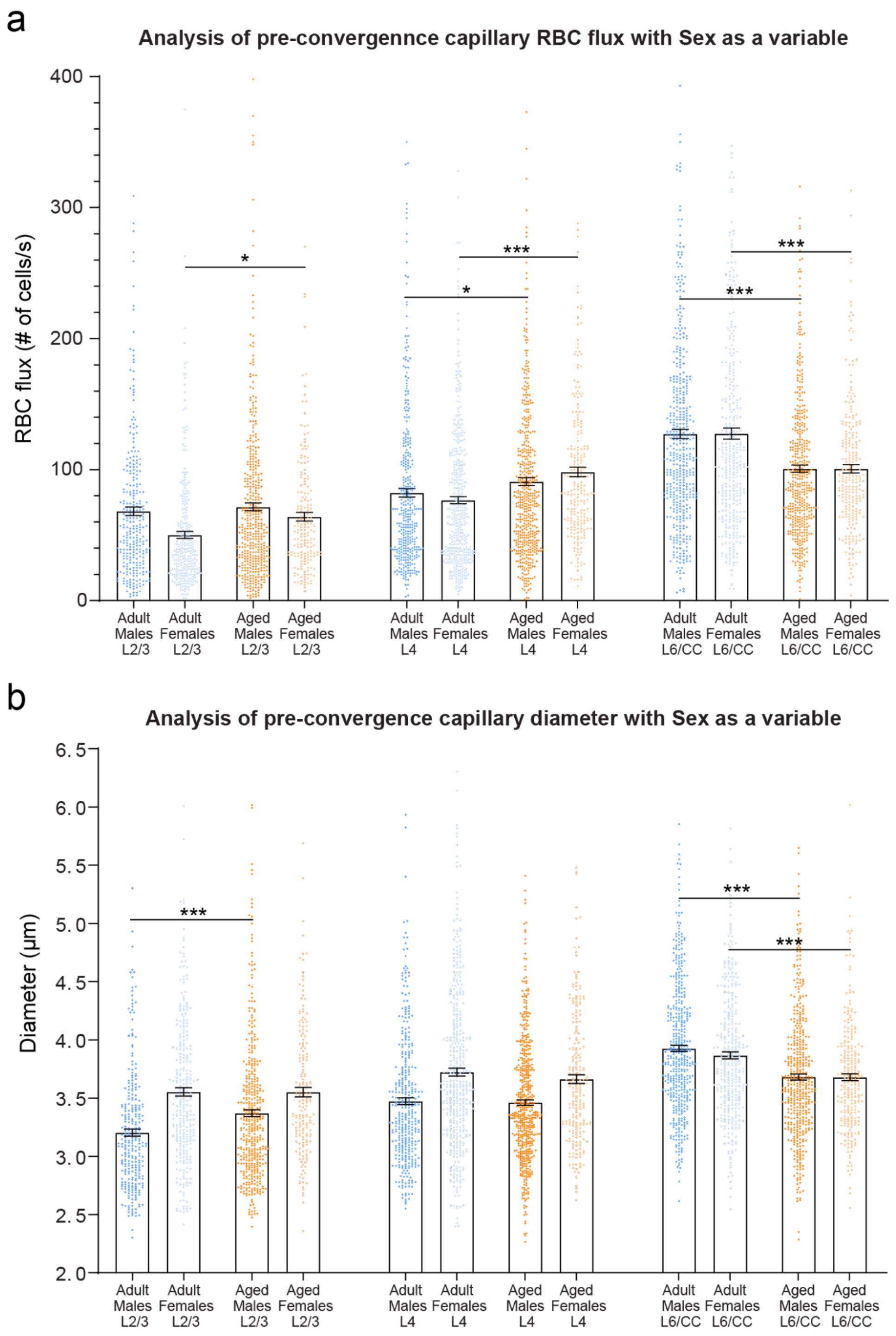


**Supplementary Figure 5. Sex differences in pre-convergence capillary flux and diameter of isoflurane anesthetized mice. a.** RBC flux in adult and aged mice analyzed across cortical layers and separated into male and female sexes. Three-way ANOVA analysis with Sidak's post-hoc comparison testing, Sex $F(1,4112)=3.886$; $p=0.049$. Layer 2/3 Adult females vs Aged females, $p=0.017$. Layer 4, Adult males vs Aged males, $p=0.042$, Adult females vs Aged females, $p<0.001$. Layer 6/CC, Adult males vs Aged males, $p<0.001$, Adult females vs Aged females, $p<0.001$. Adult, n=12 mice for layers 2/3 and 4, and 11 mice for layer 6/CC (6 males and 6 females); Aged, n=11 mice for all layers (8 males and 3 females). Data shown as mean ± SEM. **b.** Plot of pre-convergence capillary diameter in adult and aged mice analyzed across cortical layers and separated into male and female sexes. Three-way ANOVA analysis with Sidak's post-hoc comparison testing, Sex $F(1,4203)=69.895$; $p<0.001$. Layer 2/3 Adult males vs Aged males, $p<0.001$. Layer 6/CC, Adult males vs Aged males, $p<0.001$, Adult females vs Aged females, $p<0.001$. Adult, n=12 mice for layers 2/3 and 4, and 11 mice for layer 6/CC (6 males and 6 females); Aged, n=11 mice for all layers (8 males and 3 females). Data shown as mean ± SEM.

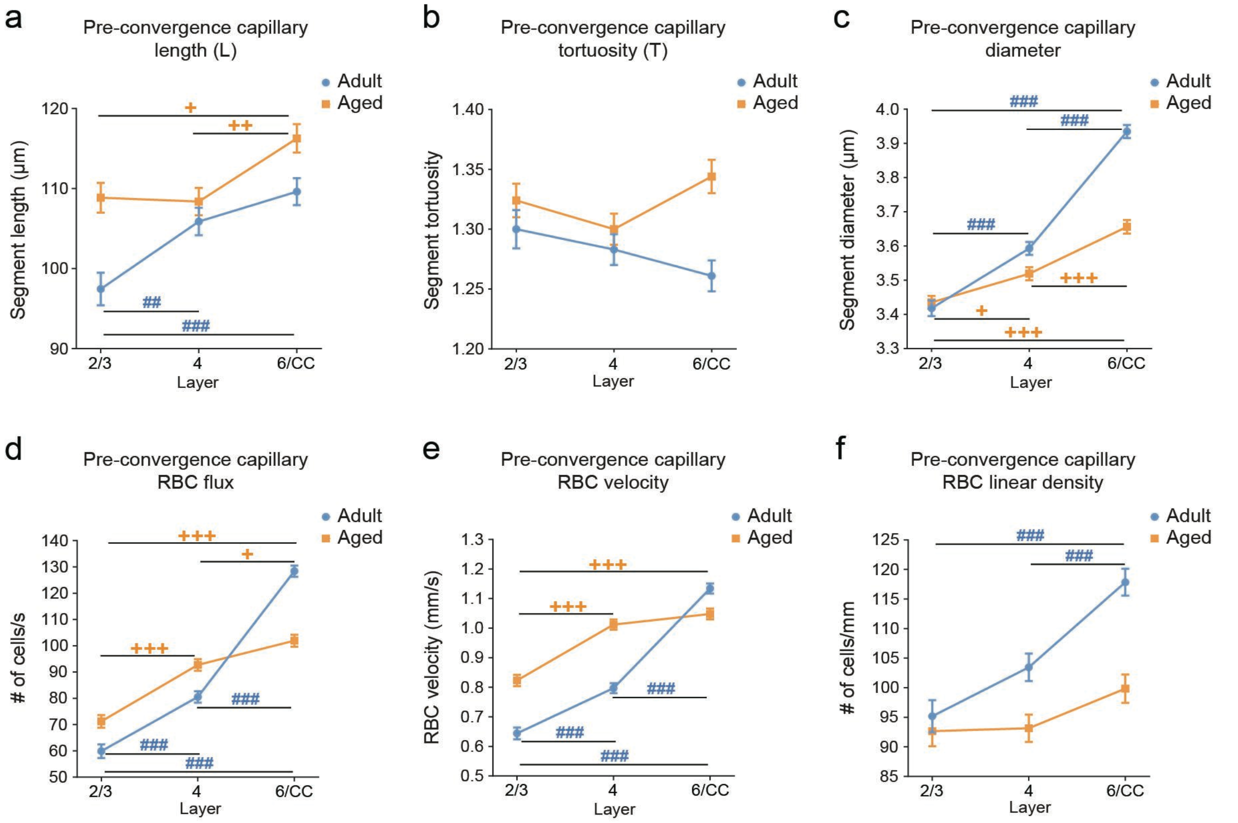


**Supplementary Figure 6. Age and cortical layer-specific changes in pre-convergence capillaries under isoflurane anesthesia. a.** Length of pre-convergence capillaries across different cortical layers within age groups. Two-way ANOVA analysis with Holm-Sidak's post-hoc comparison testing, Layer $F(2,4127)=15.139$; $p<0.001$. Adult, 2/3 vs. 4 $p=0.005$, 2/3 vs. 6/CC $p<0.001$, 4 vs. 6/CC $p=0.323$; Aged, 2/3 vs. 4 $p=0.997$, 2/3 vs. 6/CC $p=0.012$, 4 vs. 6/CC $p=0.004$. **b.** Tortuosity of pre-convergence capillaries across different cortical layers within age groups. Two-way ANOVA analysis with Holm-Sidak's post-hoc comparison testing, Layer $F(2,4127)=1.073$; $p=0.342$. Adult, 2/3 vs. 4 $p=0.774$, 2/3 vs. 6/CC $p=0.148$, 4 vs. 6/CC $p=0.555$; Aged, 2/3 vs. 4 $p=0.528$, 2/3 vs. 6/CC $p=0.673$, 4 vs. 6/CC $p=0.061$. **c.** Diameter of pre-convergence capillaries across different cortical layers within age groups. Two-way ANOVA analysis with Holm-Sidak's post-hoc comparison testing, Layer $F(2,4147)=170.157$; $p<0.001$. Adult, 2/3 vs. 4 $p<0.001$, 2/3 vs. 6/CC $p<0.001$, 4 vs. 6/CC $p<0.001$; Aged, 2/3 vs. 4 $p=0.010$, 2/3 vs. 6/CC $p<0.001$, 4 vs. 6/CC $p<0.001$. **d.** RBC flux in pre-convergence capillaries across different cortical layers within age groups. Two-way ANOVA analysis with Holm-Sidak's post-hoc comparison testing, Layer $F(2,4056)=228.048$; $p<0.001$. Adult, 2/3 vs. 4 $p<0.001$, 2/3 vs. 6/CC $p<0.001$, 4 vs. 6/CC $p<0.001$; Aged, 2/3 vs. 4 $p<0.001$, 2/3 vs. 6/CC $p<0.001$, 4 vs. 6/CC $p=0.011$. **e.** RBC velocity in pre-convergence capillaries across different cortical layers within age groups. Two-way ANOVA analysis with Holm-Sidak's post-hoc comparison testing, Layer $F(2,4053)=191.197$; $p<0.001$. Adult, 2/3 vs. 4 $p<0.001$, 2/3 vs. 6/CC $p<0.001$, 4 vs. 6/CC $p<0.001$; Aged, 2/3 vs. 4 $p<0.001$, 2/3 vs. 6/CC $p<0.001$, 4 vs. 6/CC $p=0.394$. **f.** RBC linear density in pre-convergence capillaries across different cortical layers within age groups. Two-way ANOVA analysis with Holm-Sidak's post-hoc comparison testing, Layer $F(2,4043)=19.459$; $p<0.001$. Adult, 2/3 vs. 4 $p=0.061$, 2/3 vs. 6/CC $p<0.001$, 4 vs. 6/CC $p<0.001$; Aged, 2/3 vs. 4 $p=0.999$, 2/3 vs. 6/CC $p=0.115$, 4 vs. 6/CC $p=0.129$. For all plots, the Adult group includes n=12 mice for layers 2/3 and 4, and 11 mice for layer 6/CC; the Aged group includes n=11 mice for all layers. Data shown as mean ± SEM. # denotes statistically significant differences between layers in adult mice, + denotes statistically significant differences between layers in aged mice.

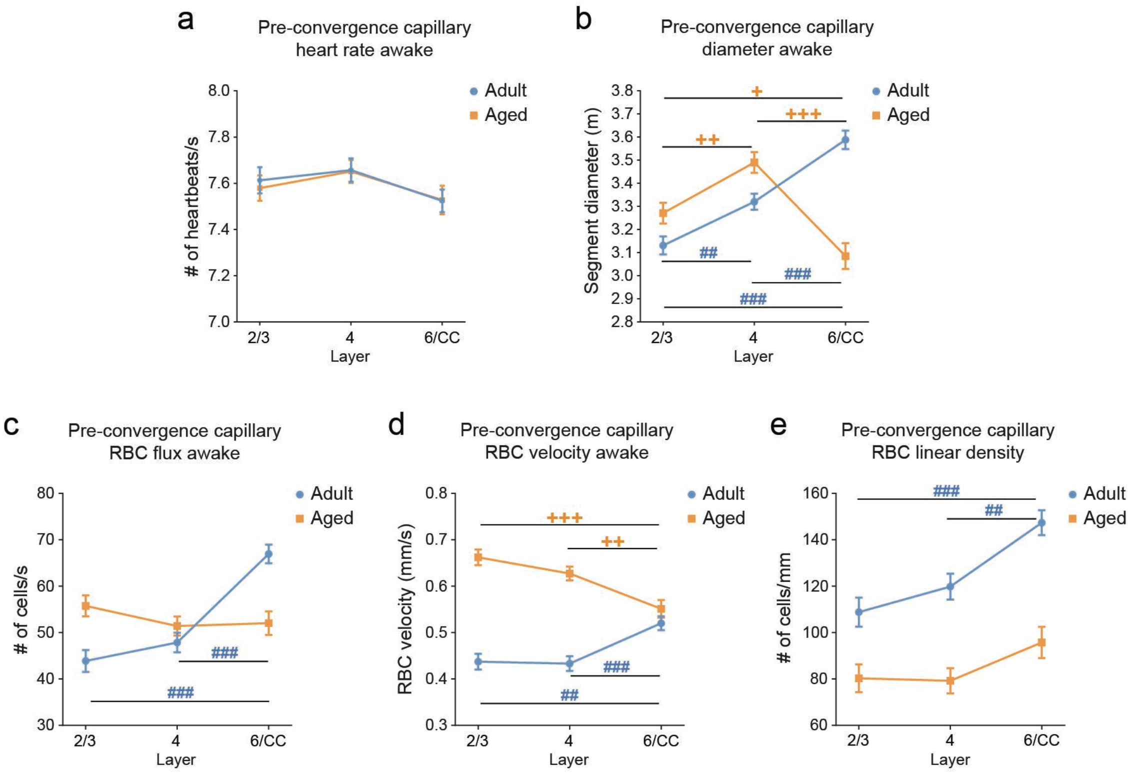


**Supplementary Figure 7. Age and cortical layer-specific changes in pre-convergence capillaries in the awake state. a.** Heart rate in pre-convergence capillary line scans across different cortical layers within age groups. Two-way ANOVA analysis with Holm-Sidak's post-hoc comparison testing, Layer F(2,1958)=1.223; p=0.295. Adult, 2/3 vs. 4 p=0.918, 2/3 vs. 6/CC p=0.552, 4 vs. 6/CC p=0.169; Aged, 2/3 vs. 4 p=0.993; 2/3 vs. 6/CC p=0.903; 4 vs. 6/CC p=0.967. **b.** Diameter of pre-convergence capillaries across different cortical layers within age groups. Two-way ANOVA analysis with Holm-Sidak's post-hoc comparison testing, Layer F(2,859)=12.374; p<0.001. Adult, 2/3 vs. 4 p=0.001, 2/3 vs. 6/CC p<0.001, 4 vs. 6/CC p<0.001; Aged, 2/3 vs. 4 p=0.002; 2/3 vs. 6/CC p=0.030; 4 vs. 6/CC p<0.001. **c.** RBC flux in pre-convergence capillaries across different cortical layers within age groups. Two-way ANOVA analysis with Holm-Sidak's post-hoc comparison testing, Layer F(2,1943)=11.643; p<0.001; Adult, 2/3 vs. 4 p=0.503, 2/3 vs. 6/CC p<0.001, 4 vs. 6/CC p<0.001; Aged, 2/3 vs. 4 p=0.398, 2/3 vs. 6/CC p=0.621, 4 vs 6/CC p=0.997. **d.** RBC velocity in pre-convergence capillaries across different cortical layers within age groups. Two-way ANOVA analysis with Holm-Sidak's post-hoc comparison testing, Layer F(2,1942)=1.189; p=0.305. Adult, 2/3 vs. 4 p=0.997, 2/3 vs. 6/CC p=0.001, 4 vs. 6/CC p<0.001; Aged, 2/3 vs. 4 p=0.325, 2/3 vs. 6/CC p<0.001, 4 vs. 6/CC p=0.006. **e.** RBC linear density in pre-convergence capillaries across different cortical layers within age groups. Two-way ANOVA analysis with Holm-Sidak's post-hoc comparison testing, Layer F(2,1928)=12.630; p<0.001. Adult, 2/3 vs 4 p=0.469, 2/3 vs 6/CC p<0.001, 4 vs 6/CC p=0.001; Aged, 2/3 vs 4 p=0.999, 2/3 vs 6/CC p=0.243, 4 vs 6/CC p=0.162. For plots a, c, d and e, the Adult group included n=5 mice for all layers; the Aged group included n=6 mice for layer 2/3 and layer 4, and 5 mice for layer 6/CC. For plot c, the Adult group includes n=3 mice per layer; the Aged group includes n=2 mice per layer. Data shown as mean ± SEM. # denotes statistically significant differences between layers in adult mice, + denotes statistically significant differences between layers in aged mice.

### Other tributary vessels - anesthetized

### Structural parameters

#### Analysis of differences between age groups within cortical layers

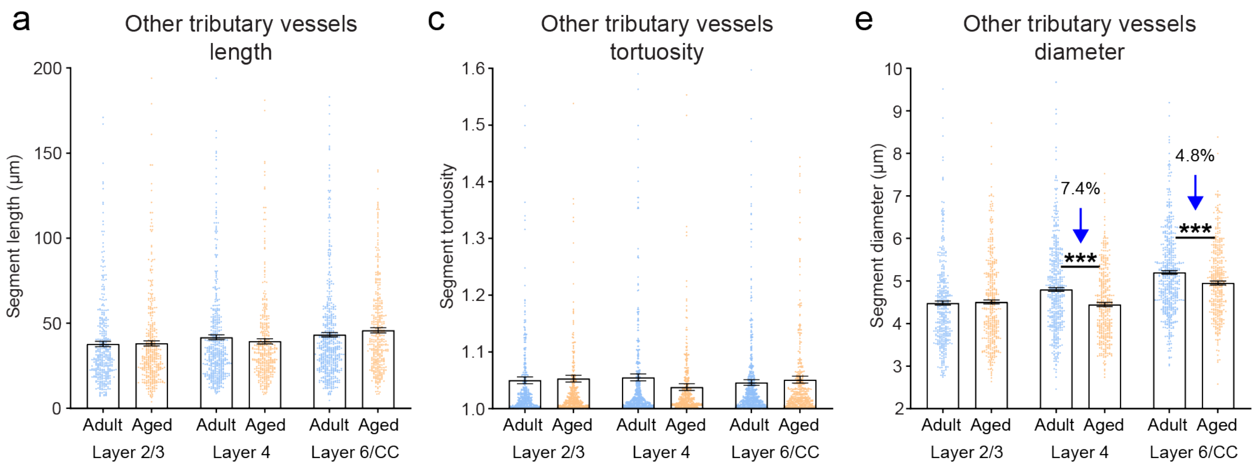


#### Analysis of differences between cortical layers within age groups

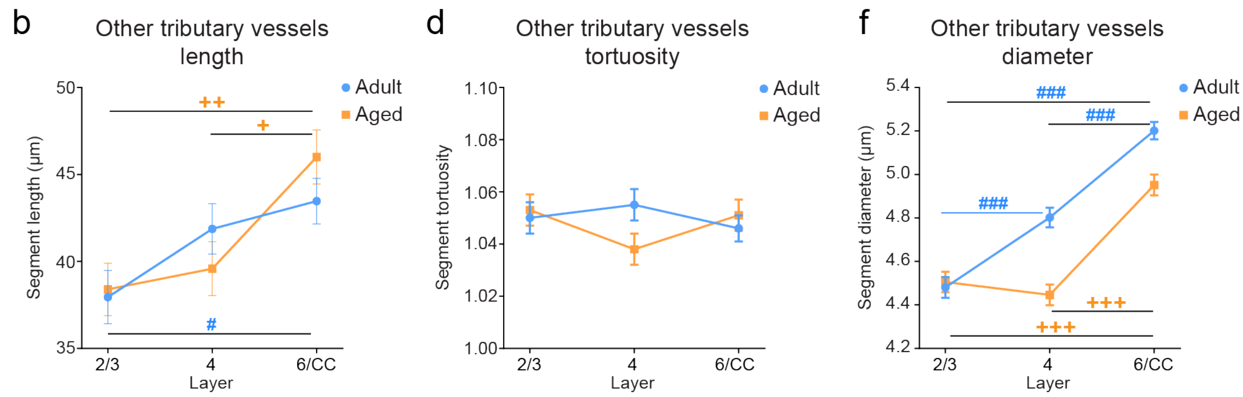


**Supplementary Figure 8. Age and cortical layer-specific changes in the structural properties of other tributary vessels. a, b.** Length of other tributary vessels across different cortical layers and age groups. Two-way ANOVA analysis with Holm-Sidak's post-hoc comparison testing, Age group F(1,2350)=0.092; p=0.762. Layer 2/3, p=0.835. Layer 4, p=0.324. Layer 6/CC, p=0.213; Layer F(2,2350)=10.112; p<0.001. Adult, 2/3 vs 4 p=0.217, 2/3 vs 6/CC p=0.019, 4 vs 6/CC p=0.736; Aged, 2/3 vs 4 p=0.927, 2/3 vs 6/CC p=0.001, 4 vs 6/CC p=0.010. **c, d.** Tortuosity of other tributary vessels across different cortical layers and age groups. Two-way ANOVA analysis with Holm-Sidak's post-hoc comparison testing, Age group F(1,2350)=0.437; p=0.509. Layer 2/3, p=0.732. Layer 4, p=0.048. Layer 6/CC, p=0.576; Layer F(2,2350)=0.237; p=0.789. Adult, 2/3 vs 4 p=0.894, 2/3 vs 6/CC p=0.959, 4 vs 6/CC p=0.582; Aged, 2/3 vs 4 p=0.261, 2/3 vs 6/CC p=0.994, 4 vs 6/CC p=0.403. **e, f.** Diameter of other tributary vessels across different cortical layers and age groups. Two-way ANOVA analysis with Holm-Sidak's post-hoc comparison testing, Age group F(1,2370)=23.667; p<0.001. Layer 2/3, p=0.706. Layer 4, p<0.001. Layer 6/CC, p<0.001; Layer F(2,2370)=90.798; p<0.001. Adult, 2/3 vs 4 p<0.001, 2/3 vs 6/CC p<0.001, 4 vs 6/CC p<0.001; Aged, 2/3 vs 4 p=0.757, 2/3 vs 6/CC p<0.001, 4 vs 6/CC p<0.001. For all plots Adult, n=12 mice for layers 2/3 and 4, and 11 mice for layer 6/CC; Aged, n=11 mice for all layers. Data shown as mean ± SEM. * denotes statistically significant differences between age groups, # denotes statistically significant differences between layers in adult mice, + denotes statistically significant differences between layers in aged mice.

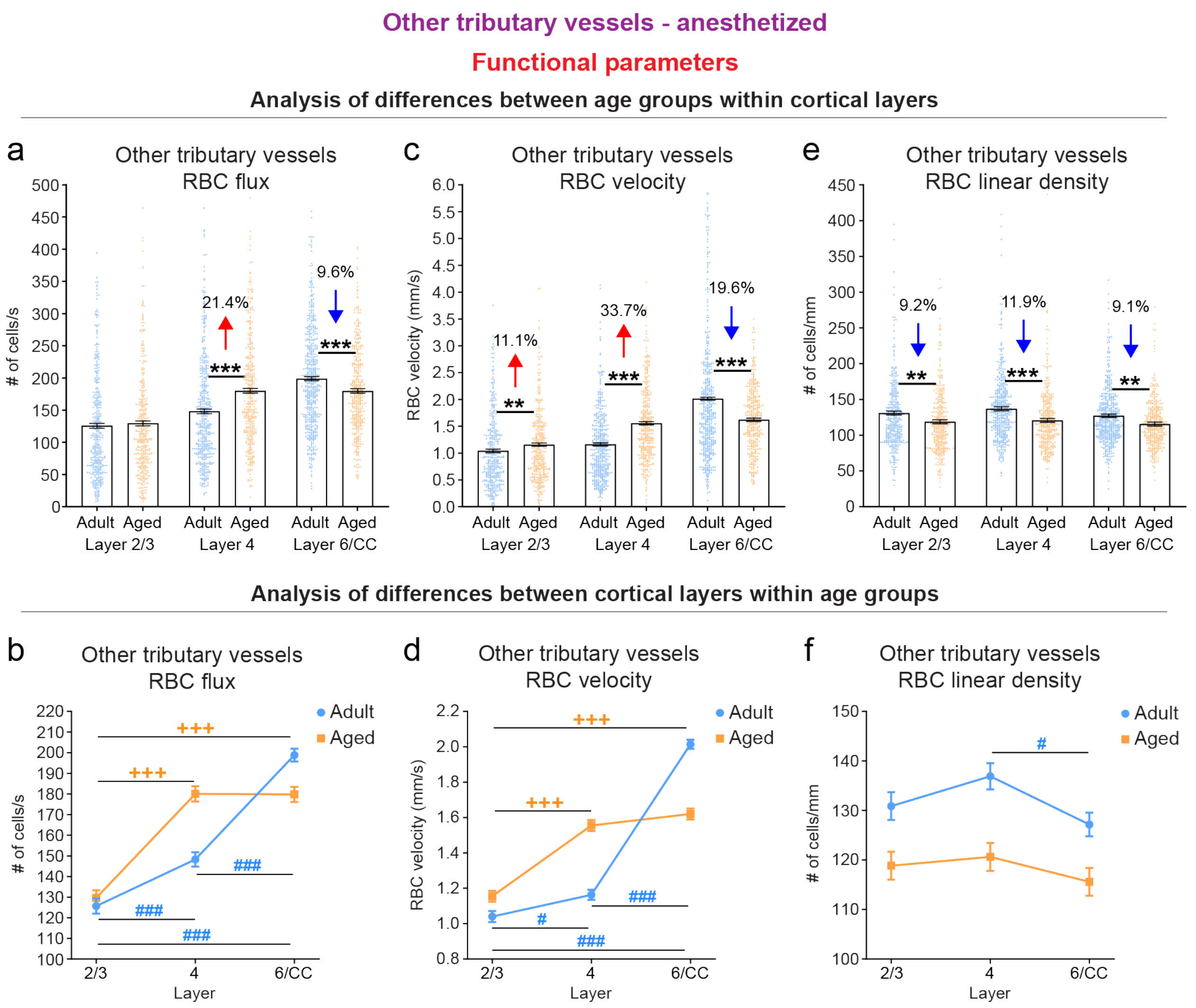


**Supplementary Figure 9. Age-dependent reduction of blood flow in other tributary vessels of L6/CC PCV vascular networks. a, b.** RBC flux in other tributary vessels across different cortical layers and age groups. Two-way ANOVA analysis with Holm-Sidak's post-hoc comparison testing, Age group $F(1,2350)=5.251$; $p=0.022$. Layer 2/3, $p=0.452$. Layer 4, $p<0.001$. Layer 6/CC, $p<0.001$; Layer $F(2,2350)=149.464$; $p<0.001$. Adult, 2/3 vs 4 $p<0.001$, 2/3 vs 6/CC $p<0.001$, 4 vs 6/CC $p<0.001$; Aged, 2/3 vs 4 $p<0.001$, 2/3 vs 6/CC $p<0.001$, 4 vs 6/CC $p=1.000$. **c, d.** RBC velocity in other tributary vessels across different cortical layers and age groups. Two-way ANOVA analysis with Holm-Sidak's post-hoc comparison testing, Age group $F(1,2353)=4.929$; $p=0.027$. Layer 2/3, $p=0.009$. Layer 4, $p<0.001$. Layer 6/CC, $p<0.001$; Layer $F(2,2353)= 302.663$; $p<0.001$. Adult, 2/3 vs 4 $p=0.012$, 2/3 vs 6/CC $p<0.001$, 4 vs 6/CC $p<0.001$; Aged, 2/3 vs 4 $p<0.001$, 2/3 vs 6/CC $p<0.001$, 4 vs 6/CC $p=0.371$. **e, f.** RBC linear density in other tributary vessels across different cortical layers and age groups. Two-way ANOVA analysis with Holm-Sidak's post-hoc comparison testing, Age group $F(1,2344)=36.151$; $p<0.001$. Layer 2/3, $p=0.003$. Layer 4, $p<0.001$. Layer 6/CC, $p=0.002$; Layer $F(2,2344)=4.191$; $p=0.015$. Adult, 2/3 vs 4 $p=0.316$, 2/3 vs 6/CC $p=0.679$, 4 vs 6/CC $p=0.019$; Aged, 2/3 vs 4 $p=0.959$, 2/3 vs 6/CC $p=0.800$, 4 vs 6/CC $p=0.500$. For all plots Adult, n=12 mice for layers 2/3 and 4, and 11 mice for layer 6/CC; Aged, n=11 mice for all layers. Data shown as mean ± SEM. * denotes statistically significant differences between age groups, # denotes statistically significant differences between layers in adult mice, + denotes statistically significant differences between layers in aged mice.

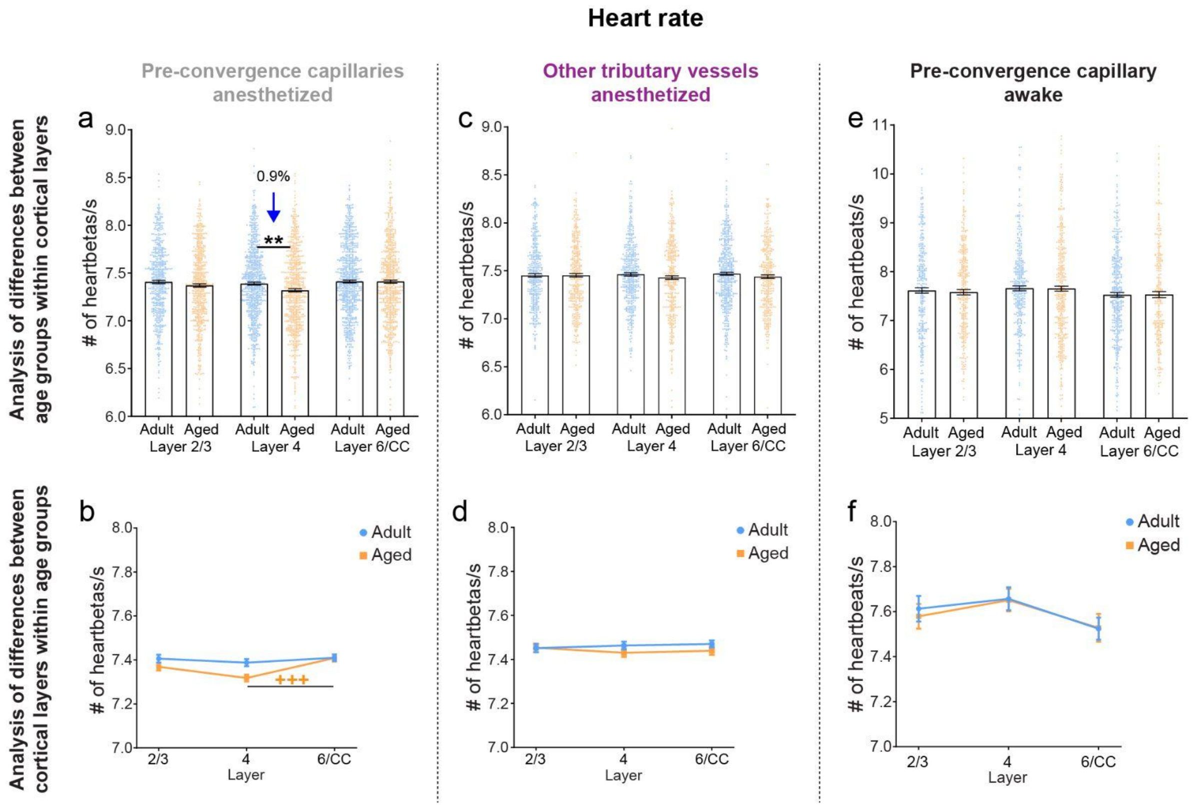


**Supplementary Figure 10. Analysis of heart rate in tributary vessels. a, b.** Heart rate measured in line scans collected from pre-convergence capillaries across different cortical layers and age groups under isoflurane anesthesia. Two-way ANOVA analysis with Holm-Sidak's post-hoc comparison testing, Age group F(1,4053)=6.862; p=0.009. Layer 2/3, p=0.148. Layer 4, p=0.002. Layer 6/CC, p=0.970; Layer F(2,4053)=6.128; p=0.002. Adult, 2/3 vs 4 p=0.850, 2/3 vs 6/CC p=0.997, 4 vs 6/CC p=0.681; Aged, 2/3 vs 4 p=0.086, 2/3 vs 6/CC p=0.249, 4 vs 6/CC p<0.001. **c, d.** Heart rate measured in line scans collected from other tributary vessels across different cortical layers and age groups under isoflurane anesthesia. Two-way ANOVA analysis with Holm-Sidak's post-hoc comparison testing, Age group F(1,2353)=1.829; p=0.176. Layer 2/3, p=0.965. Layer 4, p=0.217. Layer 6/CC, p=0.220; Layer F(2,2353)=0.075; p=0.928. Adult, 2/3 vs 4 p=0.961, 2/3 vs 6/CC p=0.846, 4 vs 6/CC p=0.989; Aged, 2/3 vs 4 p=0.798, 2/3 vs 6/CC p=0.943, 4 vs 6/CC p=0.985. **e, f.** Heart rate measured in line scans collected from pre-convergence capillaries across different cortical layers and age groups in the awake state. Two-way ANOVA analysis with Holm-Sidak's post-hoc comparison testing, Age group F(1,1958)=0.848; p=0.357; Layer 2/3, p=0.664; Layer 4, p=0.177; Layer 6/CC, p=0.960. Layer F(2,1958)=1.223; p=0.295. Adult, 2/3 vs. 4 p=0.918, 2/3 vs. 6/CC p=0.552, 4 vs. 6/CC p=0.169; Aged, 2/3 vs. 4 p=0.993; 2/3 vs. 6/CC p=0.903; 4 vs. 6/CC p=0.967. For all plots Adult, n=12 mice for layers 2/3 and 4, and 11 mice for layer 6/CC; Aged, n=11 mice for all layers. Data shown as mean ± SEM. * denotes statistically significant differences between age groups, + denotes statistically significant differences between layers in aged mice.

### Pre-convergence capillaries

#### Functional parameters and diameter anesthetized (ANE) vs. awake (AWA)

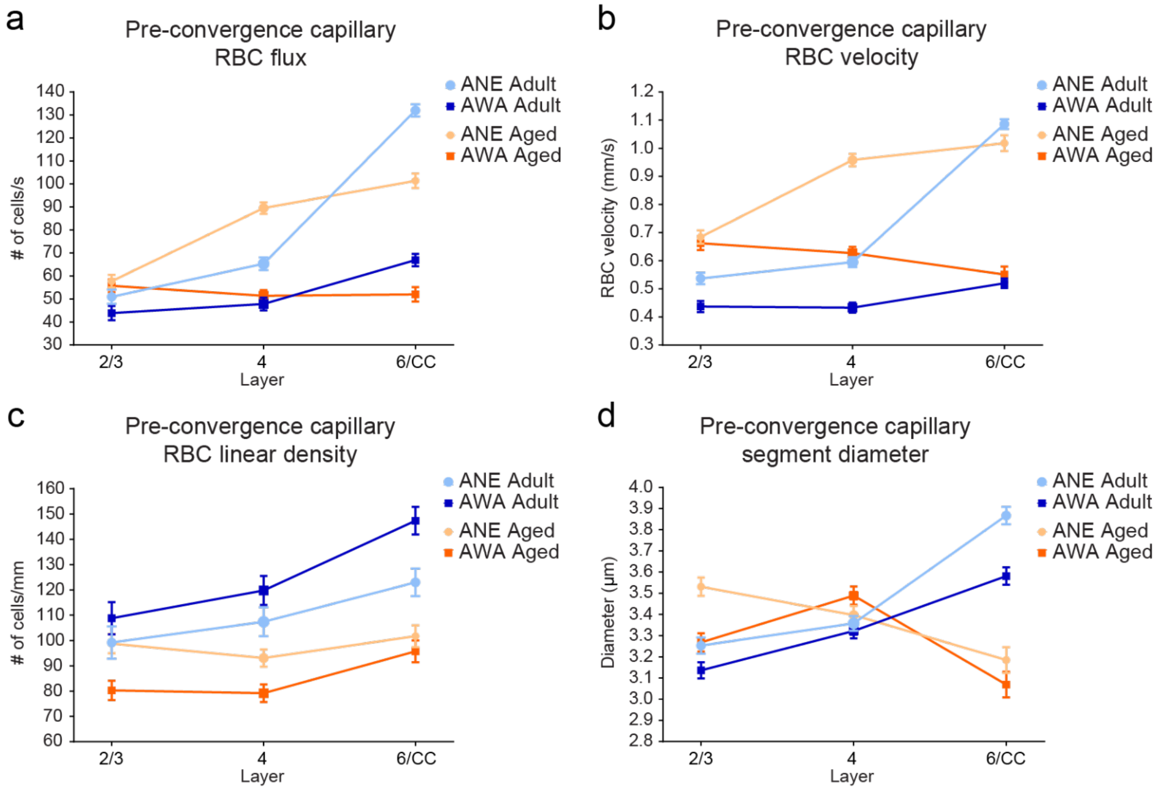


e

**Change in hemodynamic parameters and diameter (isoflurane anesthetized / awake state)**

| Parameter | Age group | Adult | | | Aged | | |
|---|---|---|---|---|---|---|---|
| | Layer | Layer 2/3 | Layer 4 | Layer 6/CC | Layer 2/3 | Layer 4 | Layer 6/CC |
| RBC flux | p value | 0.105 | <0.001 | <0.001 | 0.614 | <0.001 | <0.001 |
| | magnitude of change (%) | +16.21 | +36.51 | +97.19 | +3.55 | +74.22 | +94.95 |
| RBC velocity | p value | 0.001 | <0.001 | <0.001 | 0.517 | <0.001 | <0.001 |
| | magnitude of change (%) | +22.88 | +37.41 | +108.84 | +3.32 | +52.79 | +84.75 |
| RBC linear density | p value | 0.288 | 0.125 | 0.002 | 0.001 | 0.005 | 0.326 |
| | magnitude of change (%) | -8.48 | -10.34 | -16.55 | +22.28 | +17.51 | +6.28 |
| Segment diameter | p value | 0.032 | 0.480 | <0.001 | <0.001 | 0.128 | 0.184 |
| | magnitude of change (%) | +3.73 | +1.08 | +7.99 | 8.05 | -2.64 | +3.78 |

**Supplementary Figure 11. Comparison of hemodynamic parameters and diameter of pre-convergence capillaries between the awake and anesthetized state. a-d.** RBC flux (a), RBC velocity (b), RBC linear density (c) and diameter (d) of pre-convergence capillaries across different cortical layers and age groups in the anesthetized and awake state. For plots in a, b and c, Adult, n=5 mice per layer; Aged, n=5 mice per layer. For plot in d, Adult, n=3 mice per layer; Aged, n=2 mice per layer. Data shown as mean ± SEM. **e.** Table summarizing the statistical outcomes after comparison of each parameter in the anesthetized and the awake state of the same layer within age group. Underlined p values denote statistical significance. Two-way ANOVA analysis with Holm-Sidak post-hoc comparison testing. The included magnitude of change values describes the increase or decrease in the mean value of the parameter for the same layer in the anesthetized compared to the awake state.

**Pearson correlation between structural parameters and RBC flux of pre-convergence capillaries under anesthesia**

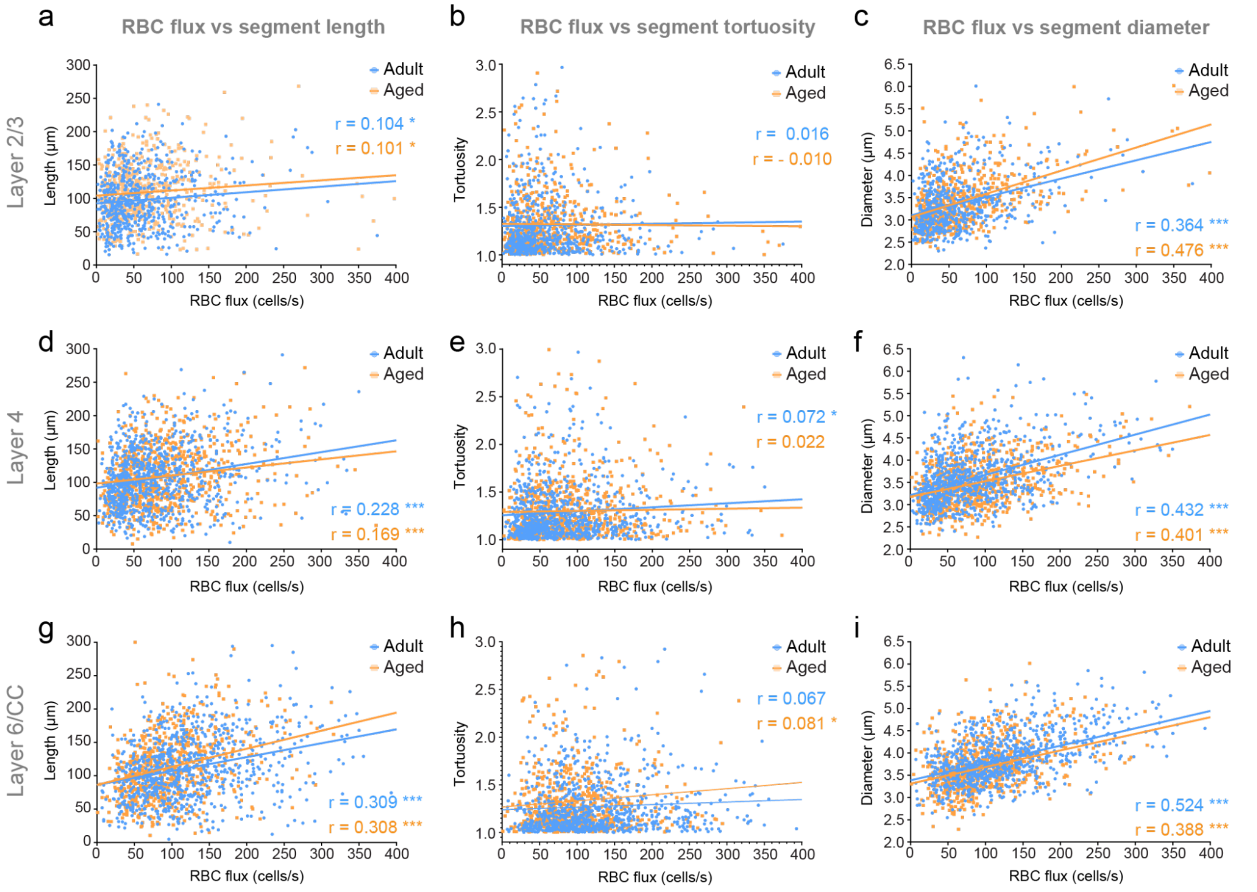


**Supplementary Figure 12. Analysis of correlation between structural parameters and RBC flux of pre-convergence capillaries in the anesthetized state**. Plots of correlation analysis between RBC flux and length (**a, d, g**), tortuosity (**b, e, h**) or diameter **(c, f, i**) of pre-convergence capillaries across different cortical layers and age groups. The values of the correlation coefficient (r) and the statistical significance of the correlation for both age groups are denoted in each plot. Pearson correlation analysis; Adult, n=609 analyzed vessels for layer 2/3, n=831 analyzed vessels for layer 4, and n=836 analyzed vessels for layer for layer 6/CC; Aged, n=611 analyzed vessels for layer 2/3, n = 779 analyzed vessels for layer 4, and n=718 analyzed vessels for layer for layer 6/CC.

**Pearson correlation between structural parameters and RBC flux of pre-convergence capillaries in the awake state**

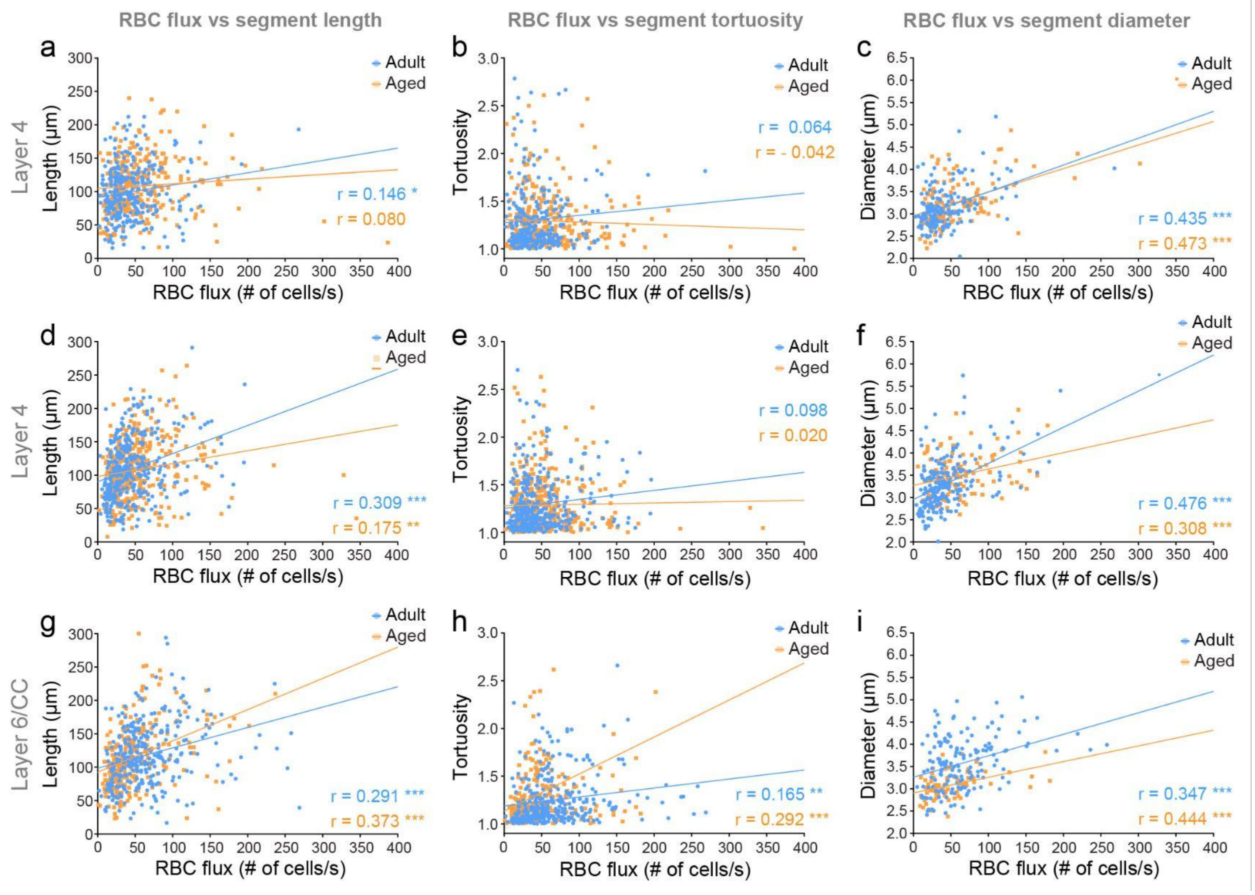


**Supplementary Figure 13. Analysis of correlation between structural parameters and RBC flux of pre-convergence capillaries in the awake state.** Plots of correlation analysis between RBC flux and length (**a, d, g**), tortuosity (**b, e, h**) or diameter (**c, f, i**) of pre-convergence capillaries across different cortical layers and age groups. The values of the correlation coefficient (r) and the statistical significance of the correlation for both age groups are denoted in each plot. Pearson correlation analysis; Adult, n=285 (173 for diameter) analyzed vessels for layer 2/3, n=349 (211 for diameter) analyzed vessels for layer 4, and n=308 (162 for diameter) analyzed vessels for layer for layer 6/CC; Aged, n=316 (123 for diameter) analyzed vessels for layer 2/3, n=384 (125 for diameter) analyzed vessels for layer 4, and n=251 (80 for diameter) analyzed vessels for layer for layer 6/CC.

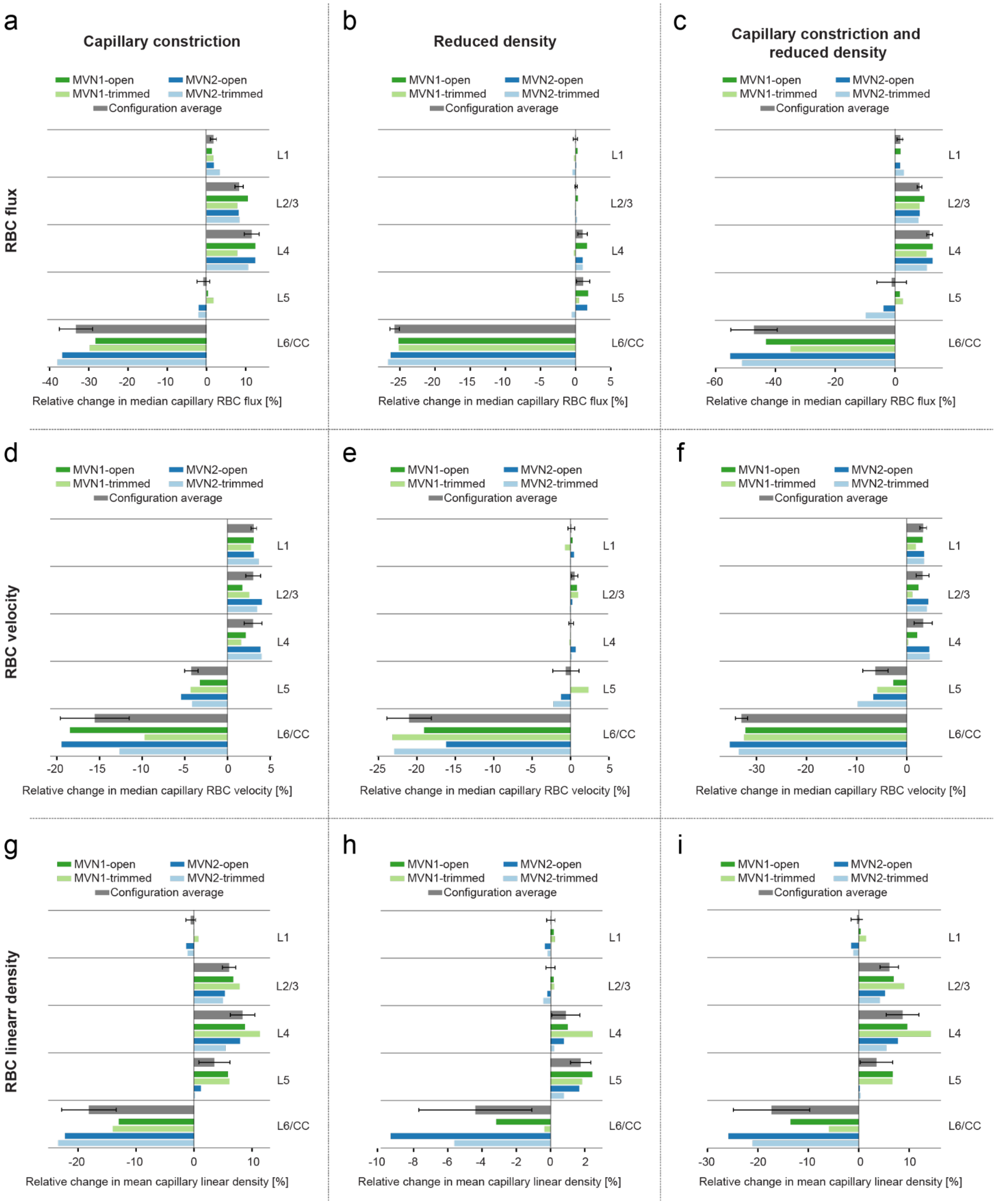


**Supplementary Figure 14. Separate effects of capillary constriction and reduced vascular density on hemodynamic properties in in silico modeling of awake conditions. a-c.** RBC flux across different cortical layers in in silico microvascular networks affected by capillary constriction in layer 6/CC (a), reduced vascular density in layer 6/CC (b), or both effects combined (c). **d-f.** RBC velocity across different cortical layers in in silico microvascular networks affected by capillary constriction in layer 6/CC (d), reduced vascular density in layer 6/CC (e), or both effects combined (f). **g-i.** RBC linear density across different cortical layers in in silico microvascular networks affected by capillary constriction in layer 6/CC (g), reduced vascular density in layer 6/CC (h), or both effects combined (i). For all plots n = 4 network configurations, 2 microvascular networks x 2 boundary conditions. All configuration average plots show median ± SD.

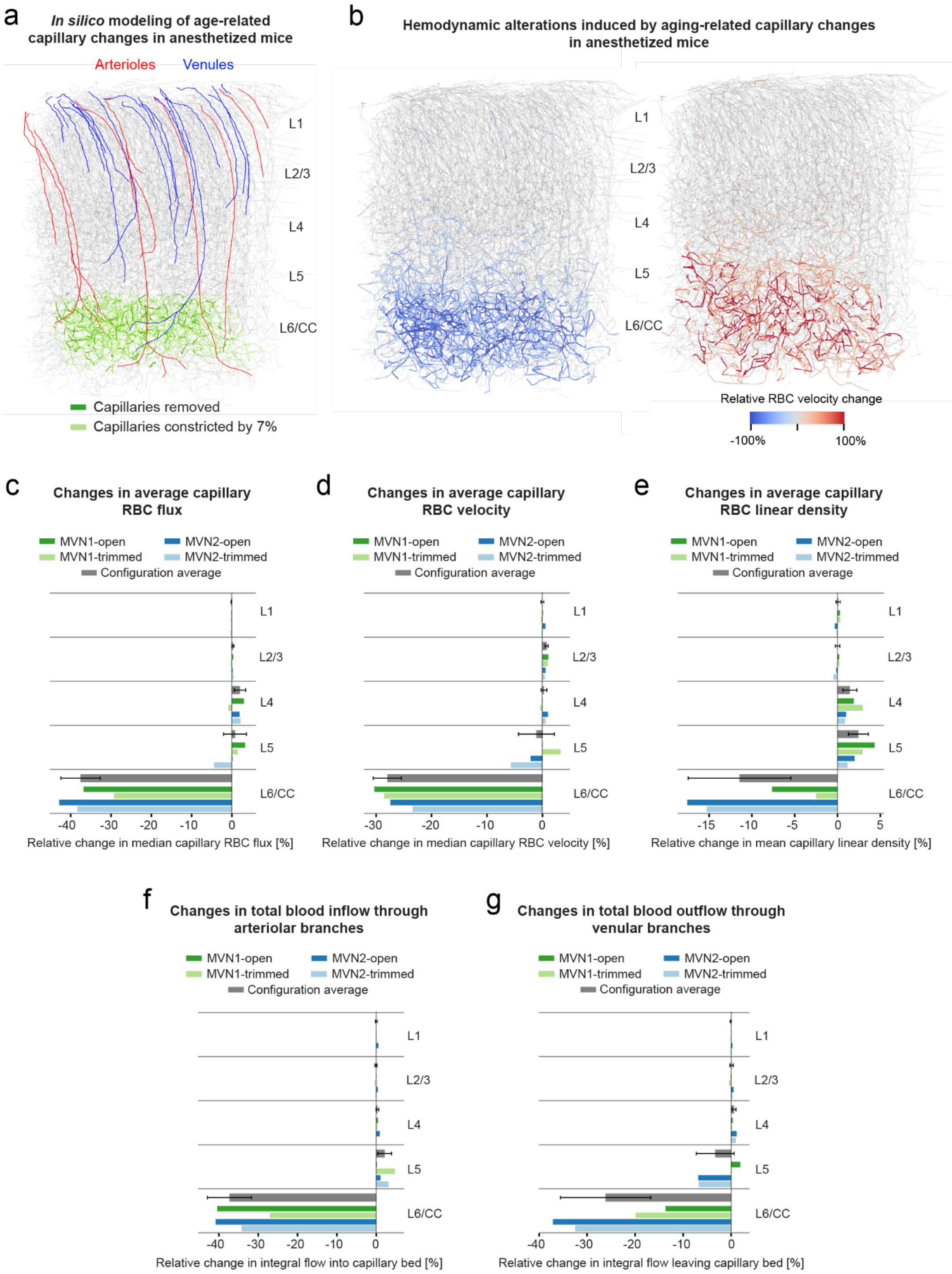


**Supplementary Figure 15. *In silico* modeling of age-related changes in capillary structure observed under isoflurane anesthesia. a.** Image of an *in silico* microvascular network with labeled penetrating arterioles (red) and ascending venules (blue). Age-related structural changes in layer 6/CC observed under isoflurane anesthesia have been implemented in the model, including capillary constriction (light green segments) and removed segments to mimic capillary regression (dark green segments). **b.** Images of an "aged" *in silico* microvascular network with highlighted segments exhibiting reduced (left panel) or increased (right panel) RBC velocity. Only vessels with an absolute change larger than 10% are displayed. Panels a and b are MVN1-open configuration (see Methods) **c-e.** Changes in average capillary RBC flux (c), RBC velocity (d) and RBC linear density (e) across different cortical layers (n = 4 network configurations, 2 microvascular networks x 2 boundary conditions). **f, g.** Changes in total blood inflow into the capillary network through arteriolar branches (f), and total blood outflow through venular branches (g) across different cortical layers (n = 4 network configurations, 2 microvascular networks x 2 boundary conditions). All configuration average plots show median ± SD.

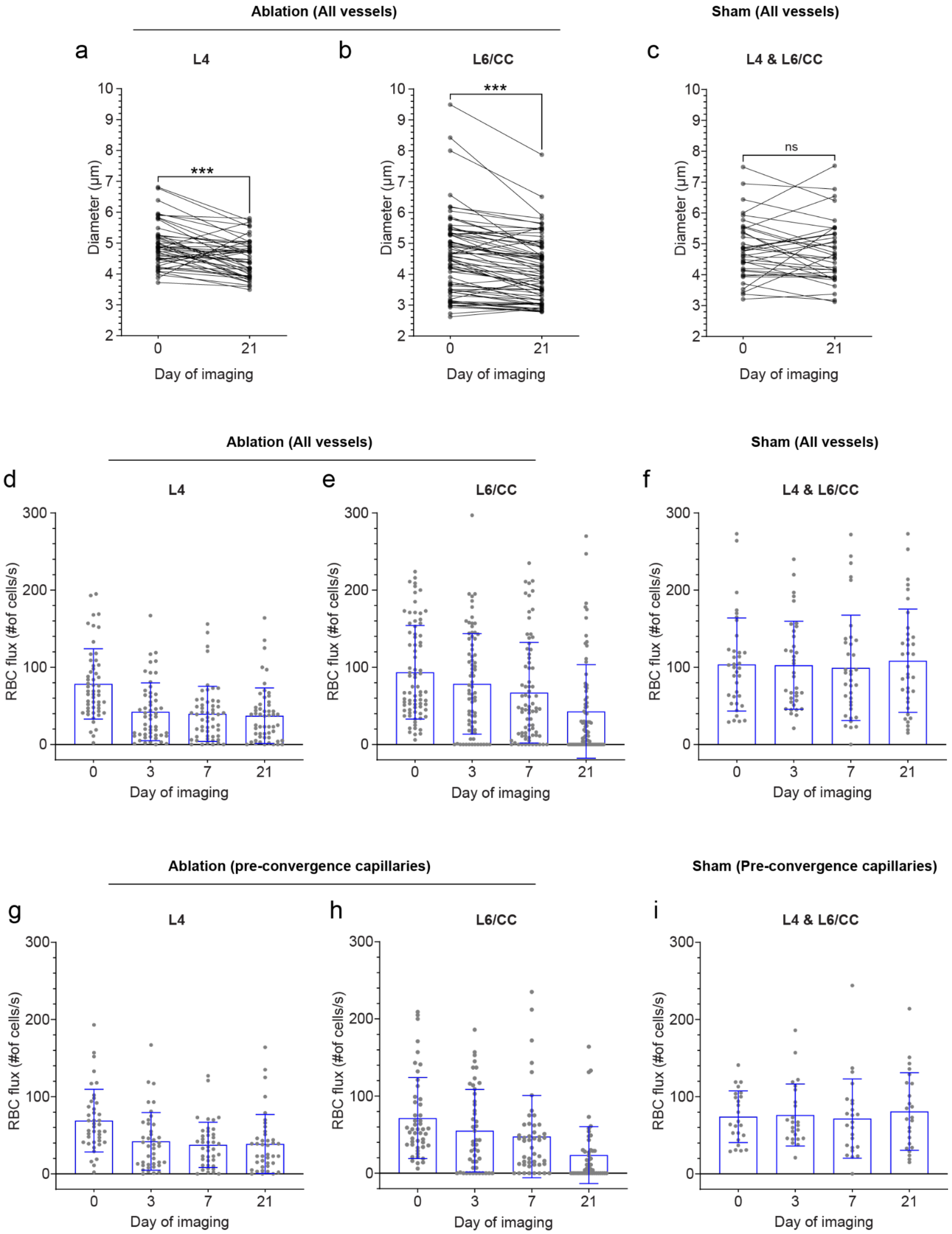


**Supplementary Figure 16. Reduction of vessel diameter and RBC flux following three-photon laser-induced capillary ablation. a-c**. Plots showing vessel diameter in analyzed microvascular networks before and 21 days after pre-convergence capillary ablation in L4 (a) and L6/CC (b), or after Sham irradiation (c). For L4, n=8 ROIs, n=49 vessels; For L6/CC, n=9 ROIs, n=72 vessels, For Sham irradiation, n=4 ROIs, n=36 vessels. For all plots, Data shown as mean ± SEM. Paired t test. L4, p = 0.0003, t,df (4.378, 48); L6/CC, p < 0.0001, t,df (6.545, 71); Sham, p = 0.479, t,df (0.7144, 35). **d-f.** Longitudinal plots showing RBC flux of vessels in analyzed microvascular networks after pre-convergence capillary ablation in L4 (d) and L6/CC (e), or after Sham irradiation (f). Day 0 denotes baseline RBC flux values, before laser irradiation was performed. Data presented as mean ± SD. For L4, n=8 ROIs, n=49 vessels; For L6/CC, n=9 ROIs, n=72 vessels, for Sham irradiation, n=4 ROIs, n=36 vessels. **g-i.** Longitudinal plots showing RBC flux of pre-convergence capillaries in analyzed microvascular networks after pre-convergence capillary ablation in L4 (g) and L6/CC (h), or after Sham irradiation (i). Day 0 denotes baseline RBC flux values, before laser irradiation was performed. Data presented as mean ± SD. For L4, n=8 ROIs, n=40 vessels; For L6/CC, n=9 ROIs, n=48 vessels, for Sham irradiation, n=4 ROIs, n=24 vessels.

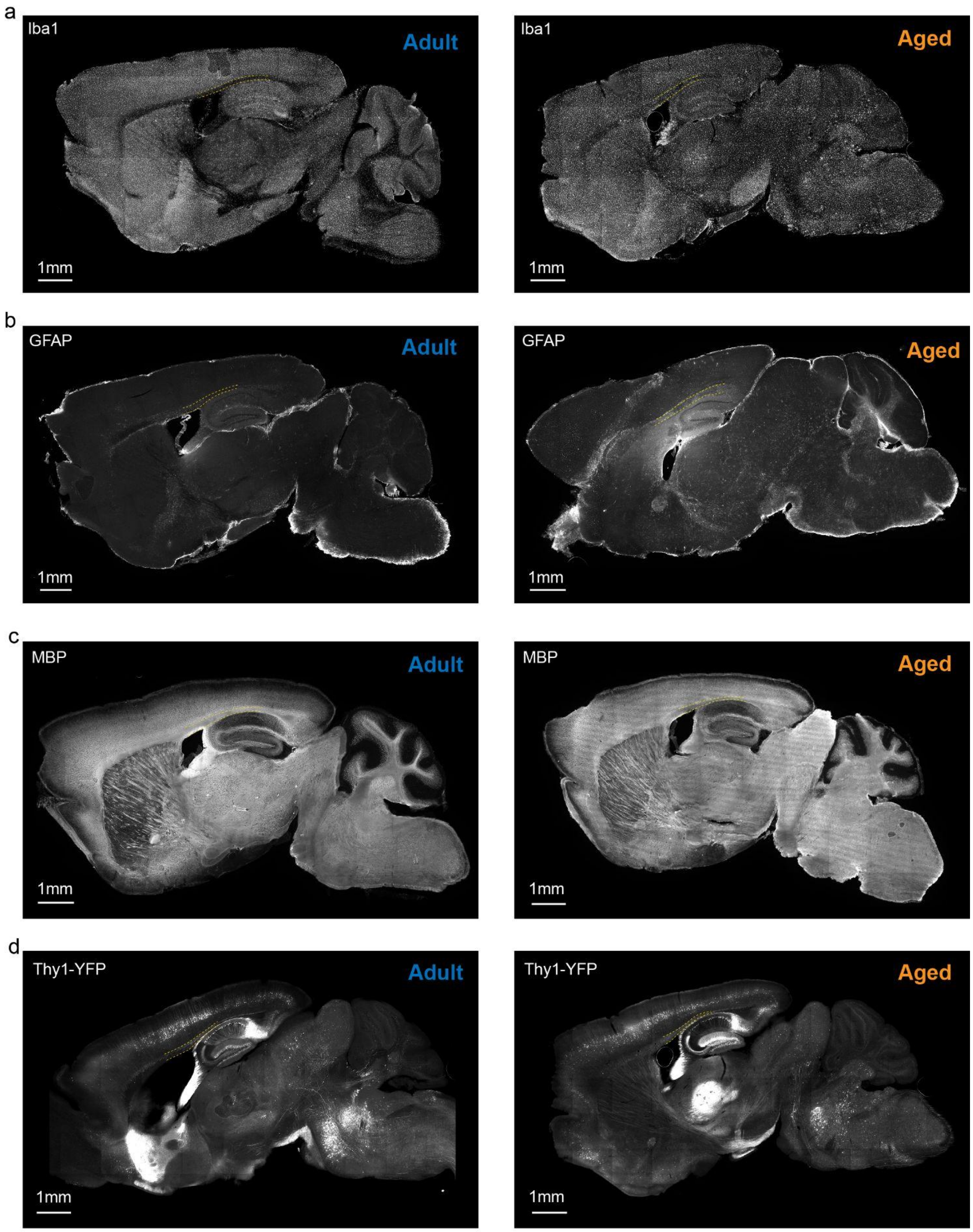


**Supplementary Figure 17. Immunohistological stains and endogenous YFP in adult and aged mice.** Epifluorescent images of adult and aged mice sagittal brain sections stained with anti-Iba1 antibody for labeling microglia (a), anti-GFAP antibody for labeling astrocytes (b), anti-MBP antibody for labeling myelin (c), and endogenous Thy-1 YFP signal (d).

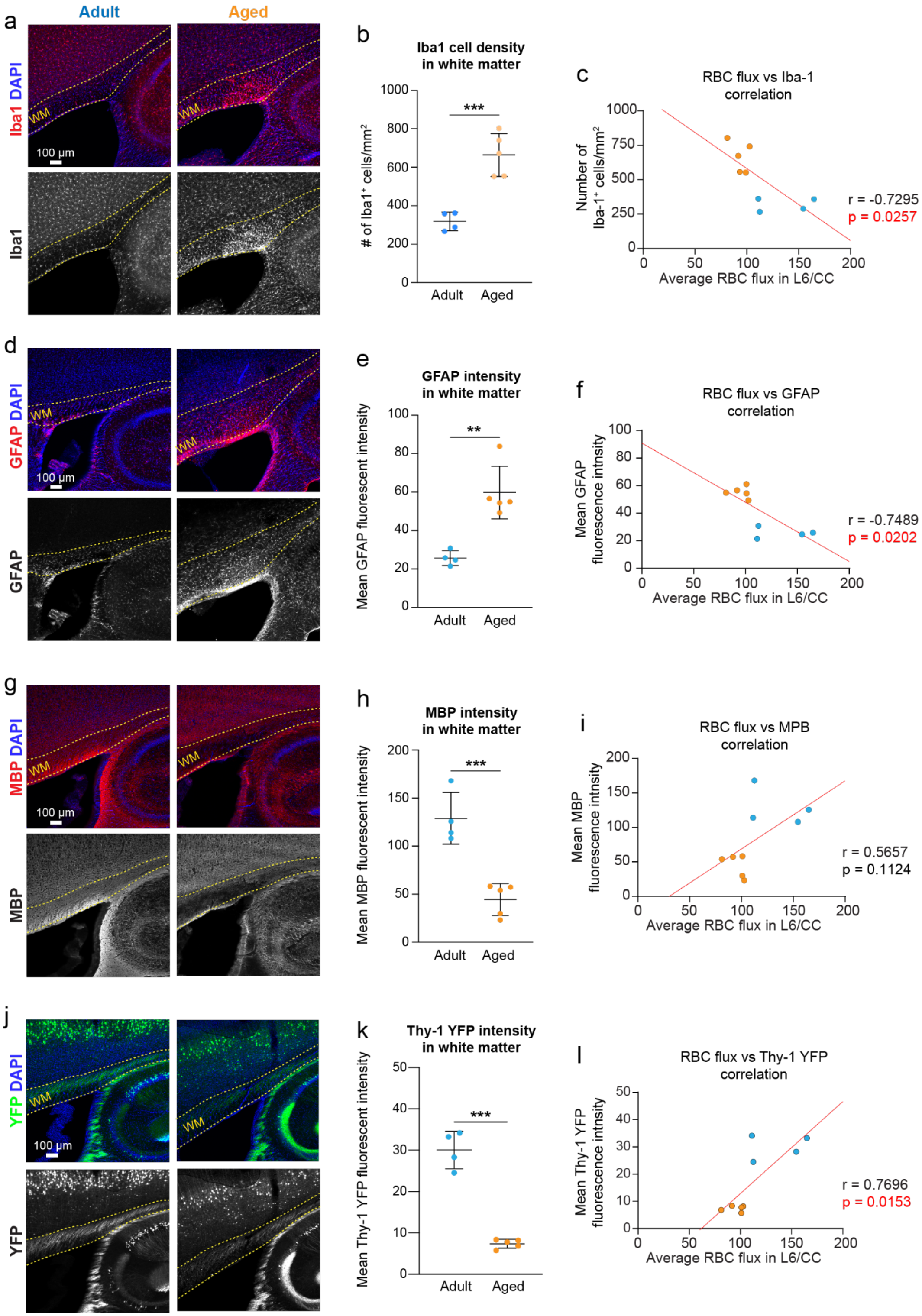


**Supplementary Figure 18. Gliosis and demyelination in the corpus callosum of aged mice. a, d, g, j.** Confocal images of adult and aged mice sagittal brain sections stained with anti-Iba1 antibody for labeling microglia (a), anti-GFAP antibody for labeling astrocytes (d), anti-MBP antibody for labeling myelin (g), and endogenous Thy-1 YFP fluorescence (j), taken in the region of the CC directly underneath the somatosensory cortex. **b, e, h, k.** Plots of Iba1 positive cell density (b), mean GFAP signal intensity (e), mean MBP signal intensity (h), and mean Thy-1 YFP signal intensity (k) in the CC analyzed in confocal images. For all plots, n=4 adult mice and 5 aged mice. Data shown as mean ± SD. Unpaired t-test. $p < 0.001$ (b); $p = 0.002$ (e); $p < 0.001$ (h); $p < 0.001$ (k). **c, f, e, l.** Plots showing Pearson correlation between average RBC flux in L6/CC measured in vivo and Iba1 positive cell density (c), mean GFAP signal intensity (f), mean MBP signal intensity (e), or mean Thy-1 YFP signal intensity (l) in the CC of corresponding animals. Correlation coefficient (r) and correlation significance (p) values are stated in each plot.

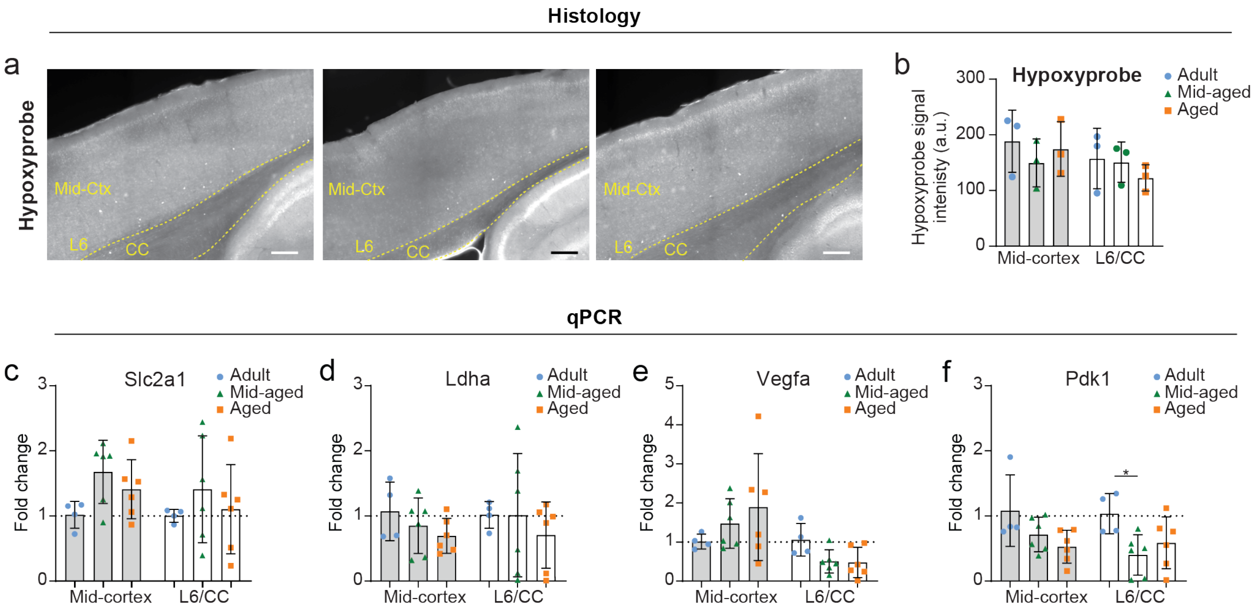


**Supplementary Figure 19. Hypoxyprobe and analysis of hypoxia-responsive gene expression in adult, mid-aged and aged mice. a,** Epifluorescent images of adult (left column), mid-aged (center column) and aged (right column) mouse somatosensory cortex in sagittal brain sections stained with anti-Hypoxyprobe antibody for detecting hypoxic tissue that was labeled by Hypoxyprobe injected i.v. in vivo. **b,** Three 150 x 300 µm ROIs in mid-cortex and three 150 x 150 µm ROIs in layer 6/CC were analyzed in each section. Plots of Hypoxyprobe signal intensity in the analyzed ROIs. N=3 adult, n=3 mid-aged, and 3 aged mice. For all plots, Data shown as mean ± SD. Two-way ANOVA with Holm-Sidak post hoc testing. * p<0.05, ** p<0.01, *** p<0.001. o-r. Plots of relative expression of glucose transporter protein type 1 gene (Slc2a1, o), Lactate dehydrogenase A gene (Ldha, p), Vascular endothelial growth factor A gene (Vegfa, q), and Pyruvate dehydrogenase kinase 1 gene (Pdk1, r), in mid-cortical and layer 6/CC brain tissue of adult, mid-aged and aged mice. For all plots, n=4 adult, 6 mid-aged and 6 aged mice. Data shown as mean ± SD. Two-way ANOVA with Holm-Sidak post hoc testing. * p<0.05.

**Excitation laser power versus cortical depth**

**(1210 nm excitation)**

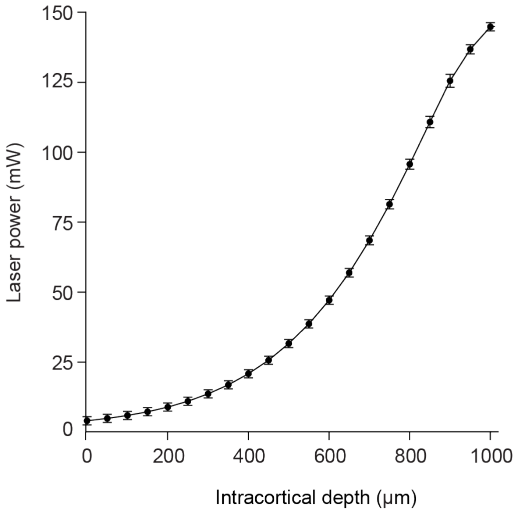


**Supplementary Figure 20. Depth dependence of laser power used for in vivo deep two-photon imaging.** Plot showing average laser power at 1210 nm excitation used for in vivo deep two photon imaging as a function of intracortical depth. Data presented as mean ± SD. N = 9 mice (5 adult and 4 aged mice).

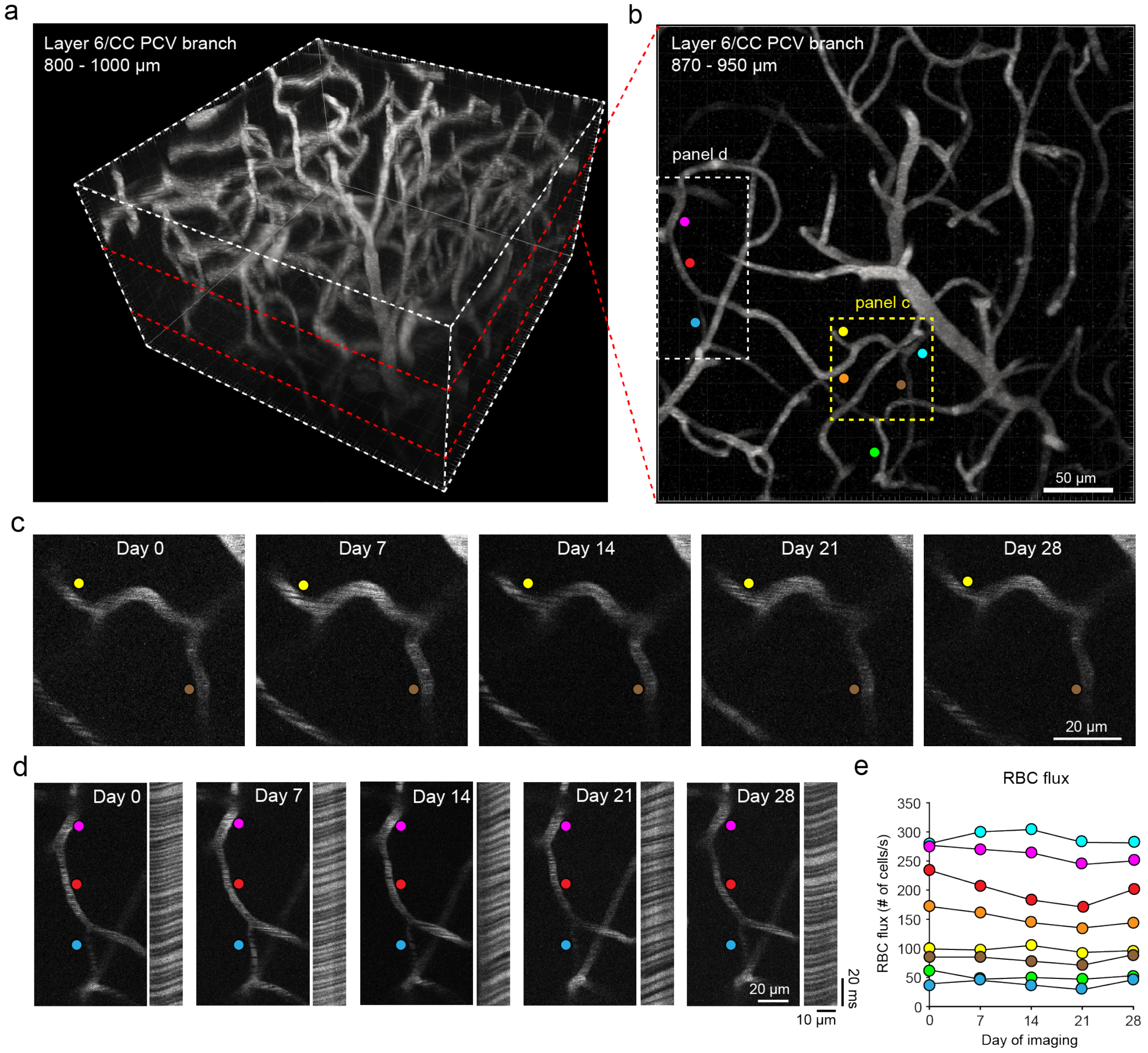


**Supplementary Figure 21. Longitudinal deep two-photon imaging of L6/CC PCV vascular network. a.** Two-photon image stack of the vasculature between 800 and 1000 µm of intracortical depth in an adult mouse, containing a Layer 6/CC PCV branch vascular network. **b.** Maximally projected top-down view of the Layer 6/CC PCV vascular network from panel (a). RBC flux in vessels segments labeled with colored dots was longitudinally analyzed once per week for 5 weeks. **c, d.** Magnified view of vessels within the PCV branching network from the region demarcated by the yellow (panel c) and white (panel d) dotted lines. Example line-scan shown on the right of panel d was obtained from the vessel marked with the red dot. **e.** RBC flux plotted as a function of imaging day for each vessel. The plot includes pre-convergence capillaries and some tributary vessels. Note the marked stability of flux over time. Imaging was performed under isoflurane anesthesia.

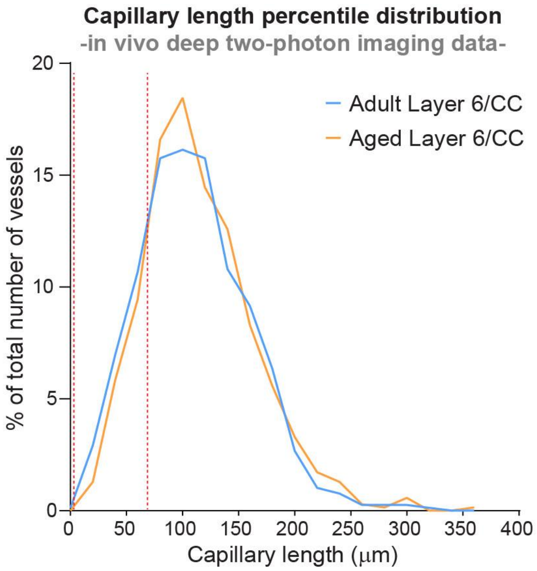


**Supplemental Figure 22. Capillary length percentile distribution from in vivo imaging data**. Plot of the percentile distribution of pre-convergence capillary length in layer 6/CC of adult and aged mice (n = 11 mice per group) from *in vivo* deep two-photon imaging experiments. The red dotted line marks the inflection point at 70 μm where the distribution of capillary length changes in aged compared to adult mice.

| Parameter 1 | Parameter 2 | Age group | Pre-convergence capillaries anesthetized | | | | | | Other tributary vessels anesthetized | | | | | |
|---|---|---|---|---|---|---|---|---|---|---|---|---|---|---|
| | | | Layer 2/3 | | Layer 4 | | Layer 6/CC | | Layer 2/3 | | Layer 4 | | Layer 6/CC | |
| | | | r | *p* | r | *p* | r | *p* | r | *p* | r | *p* | r | *p* |
| **Segment length** | **RBC flux** | Adult | 0.104 | 0.010 | 0.228 | <0.001 | 0.309 | <0.001 | 0.017 | 0.732 | 0.028 | 0.534 | 0.085 | 0.048 |
| | | Aged | 0.101 | 0.012 | 0.169 | <0.001 | 0.308 | <0.001 | 0.107 | 0.040 | 0.015 | 0.760 | 0.133 | 0.008 |
| **Segment length** | **RBC velocity** | Adult | 0.063 | 0.123 | 0.182 | <0.001 | 0.270 | <0.001 | 0.094 | 0.060 | 0.063 | 0.164 | 0.026 | 0.547 |
| | | Aged | 0.083 | 0.039 | 0.164 | <0.001 | 0.300 | <0.001 | 0.184 | <0.001 | 0.109 | 0.029 | 0.189 | <0.001 |
| **Segment tortuosity** | **RBC flux** | Adult | 0.016 | 0.701 | 0.072 | 0.037 | 0.067 | 0.054 | -0.104 | 0.039 | -0.050 | 0.272 | -0.002 | 0.964 |
| | | Aged | -0.010 | 0.803 | 0.022 | 0.536 | 0.081 | 0.030 | 0.004 | 0.937 | -0.109 | 0.028 | -0.102 | 0.042 |
| **Segment tortuosity** | **RBC velocity** | Adult | 0.017 | 0.667 | 0.040 | 0.252 | 0.071 | 0.040 | -0.054 | 0.281 | -0.031 | 0.488 | -0.008 | 0.854 |
| | | Aged | -0.004 | 0.992 | 0.023 | 0.529 | 0.081 | 0.029 | 0.041 | 0.433 | 0.006 | 0.906 | -0.023 | 0.644 |
| **Segment length** | **Segment tortuosity** | Adult | 0.341 | <0.001 | 0.374 | <0.001 | 0.376 | <0.001 | 0.567 | <0.001 | 0.640 | <0.001 | 0.519 | <0.001 |
| | | Aged | 0.300 | <0.001 | 0.423 | <0.001 | 0.382 | <0.001 | 0.456 | <0.001 | 0.608 | <0.001 | 0.522 | <0.001 |
| **Segment diameter** | **RBC flux** | Adult | 0.364 | <0.001 | 0.432 | <0.001 | 0.524 | <0.001 | 0.375 | <0.001 | 0.323 | <0.001 | 0.266 | <0.001 |
| | | Aged | 0.476 | <0.001 | 0.401 | <0.001 | 0.388 | <0.001 | 0.453 | <0.001 | 0.436 | <0.001 | 0.399 | <0.001 |
| **Segment diameter** | **RBC velocity** | Adult | 0.201 | <0.001 | 0.229 | <0.001 | 0.312 | <0.001 | 0.168 | 0.001 | 0.122 | 0.007 | 0.175 | <0.001 |
| | | Aged | 0.251 | <0.001 | 0.238 | <0.001 | 0.178 | <0.001 | 0.296 | <0.001 | 0.191 | <0.001 | 0.288 | <0.001 |
| **Segment length** | **Segment diameter** | Adult | 0.184 | <0.001 | 0.223 | <0.001 | 0.179 | <0.001 | -0.085 | 0.090 | -0.112 | 0.013 | -0.097 | 0.024 |
| | | Aged | 0.163 | <0.001 | 0.115 | 0.001 | 0.251 | <0.001 | -0.135 | 0.010 | -0.150 | 0.002 | -0.055 | 0.273 |
| **Segment diameter** | **Segment tortuosity** | Adult | 0.028 | 0.485 | 0.089 | 0.011 | 0.061 | 0.078 | -0.050 | 0.321 | -0.101 | 0.026 | -0.116 | 0.007 |
| | | Aged | 0.033 | 0.409 | 0.118 | 0.001 | 0.117 | 0.002 | -0.053 | 0.310 | -0.153 | 0.002 | -0.142 | 0.005 |
| **RBC flux** | **RBC velocity** | Adult | 0.842 | <0.001 | 0.819 | <0.001 | 0.721 | <0.001 | 0.872 | <0.001 | 0.852 | <0.001 | 0.419 | <0.001 |
| | | Aged | 0.655 | <0.001 | 0.790 | <0.001 | 0.726 | <0.001 | 0.838 | <0.001 | 0.758 | <0.001 | 0.764 | <0.001 |

**Supplementary Table 1. Pearson correlation analysis between structural and functional properties of pre-convergence capillaries and other tributary vessels under isoflurane anesthesia.** Overview of the correlation coefficient (r) and statistical significance (*p*) for Pearson correlation analysis between different parameters of pre-convergence capillaries and other tributary vessels in adult and aged mice. Underlined *p* values denote statistical significance. For all analyses on pre-convergence capillaries: Adult, n=609 analyzed vessels for layer 2/3, n=831 analyzed vessels for layer 4, and n=836 analyzed vessels for layer for layer 6/CC; Aged, n=611 analyzed vessels for layer 2/3, n=779 analyzed vessels for layer 4, and n=718 analyzed vessels for layer for layer 6/CC. For all analyses on other tributary vessels: Adult, n=398 analyzed vessels for layer 2/3, n=489 analyzed vessels for layer 4, and n=539 analyzed vessels for layer for layer 6/CC; Aged, n=365 analyzed vessels for layer 2/3, n=402 analyzed vessels for layer 4, and n=395 analyzed vessels for layer for layer 6/CC.

| Parameter 1 | Parameter 2 | Age group | Pre-convergence capillaries awake | | | | | |
|---|---|---|---|---|---|---|---|---|
| | | | Layer 2/3 | | Layer 4 | | Layer 6/CC | |
| | | | r | *p* | r | *p* | r | *p* |
| **Segment length** | **RBC flux** | Adult | 0.146 | 0.013 | 0.309 | <0.001 | 0.291 | <0.001 |
| | | Aged | 0.080 | 0.158 | 0.175 | 0.001 | 0.373 | <0.001 |
| **Segment length** | **RBC velocity** | Adult | 0.139 | 0.019 | 0.270 | <0.001 | 0.279 | <0.001 |
| | | Aged | 0.142 | 0.011 | 0.255 | <0.001 | 0.333 | <0.001 |
| **Segment tortuosity** | **RBC flux** | Adult | 0.064 | 0.282 | 0.098 | 0.068 | 0.165 | 0.004 |
| | | Aged | -0.042 | 0.452 | 0.020 | 0.698 | 0.292 | <0.001 |
| **Segment tortuosity** | **RBC velocity** | Adult | 0.058 | 0.332 | 0.039 | 0.463 | 0.152 | 0.007 |
| | | Aged | 0.002 | 0.966 | 0.046 | 0.370 | 0.241 | <0.001 |
| **Segment length** | **Segment tortuosity** | Adult | 0.307 | <0.001 | 0.384 | <0.001 | 0.366 | <0.001 |
| | | Aged | 0.271 | <0.001 | 0.482 | <0.001 | 0.361 | <0.001 |
| **Segment diameter** | **RBC flux** | Adult | 0.435 | <0.001 | 0.476 | <0.001 | 0.347 | <0.001 |
| | | Aged | 0.473 | <0.001 | 0.308 | <0.001 | 0.444 | <0.001 |
| **Segment diameter** | **RBC velocity** | Adult | 0.341 | <0.001 | 0.303 | <0.001 | 0.245 | 0.002 |
| | | Aged | 0.212 | 0.019 | 0.122 | 0.175 | 0.313 | 0.005 |
| **Segment length** | **Segment diameter** | Adult | 0.073 | 0.342 | 0.255 | <0.001 | 0.343 | <0.001 |
| | | Aged | 0.227 | 0.012 | 0.078 | 0.387 | 0.236 | 0.035 |
| **Segment diameter** | **Segment tortuosity** | Adult | 0.077 | 0.316 | 0.149 | 0.031 | 0.123 | 0.119 |
| | | Aged | 0.041 | 0.650 | 0.107 | 0.239 | 0.200 | 0.075 |
| **RBC flux** | **RBC velocity** | Adult | 0.812 | <0.001 | 0.776 | <0.001 | 0.709 | <0.001 |
| | | Aged | 0.638 | <0.001 | 0.767 | <0.001 | 0.740 | <0.001 |

**Supplementary Table 2. Pearson correlation analysis between structural and functional properties of pre-convergence capillaries in the awake state.** Overview of the correlation coefficient (r) and statistical significance (*p*) for Pearson correlation analysis between different parameters of pre-convergence capillaries in adult and aged mice. Underlined *p* values denote statistical significance. For all analyses: Adult, n=285 (173 for diameter) analyzed vessels for layer 2/3, n=349 (211 for diameter) analyzed vessels for layer 4, and n=308 (162 for diameter) analyzed vessels for layer for layer 6/CC; Aged, n=316 (123 for diameter) analyzed vessels for layer 2/3, n=384 (125 for diameter) analyzed vessels for layer 4, and n=251 (80 for diameter) analyzed vessels for layer for layer 6/CC.
