## Supplementary figures and images for "Impaired capillary-venous drainage contributes to gliosis and demyelination in white matter during aging"

### Extended Figure 1

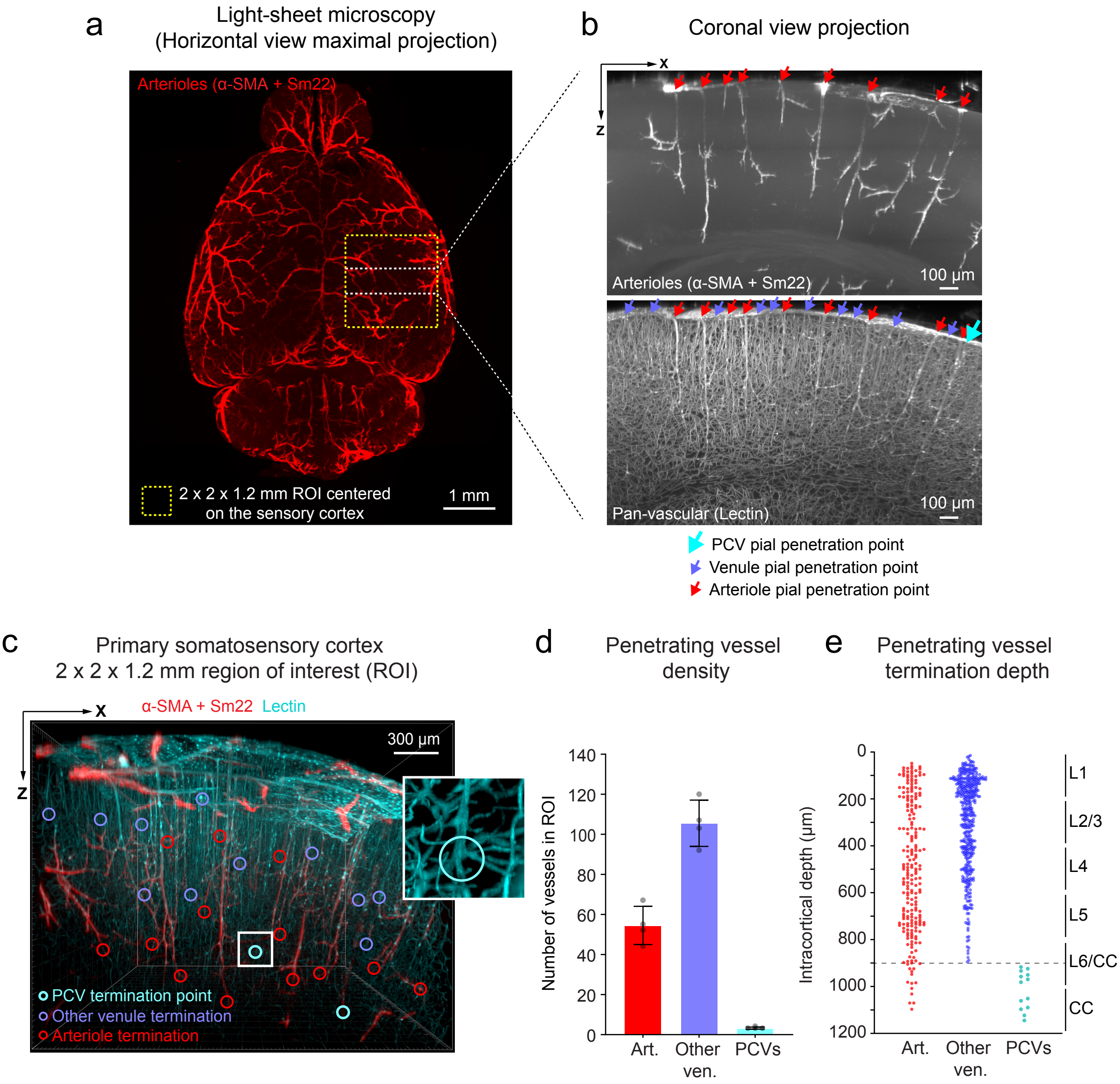


Extended Figure 1 - Stamenkovic et al.

### Extended Figure 2

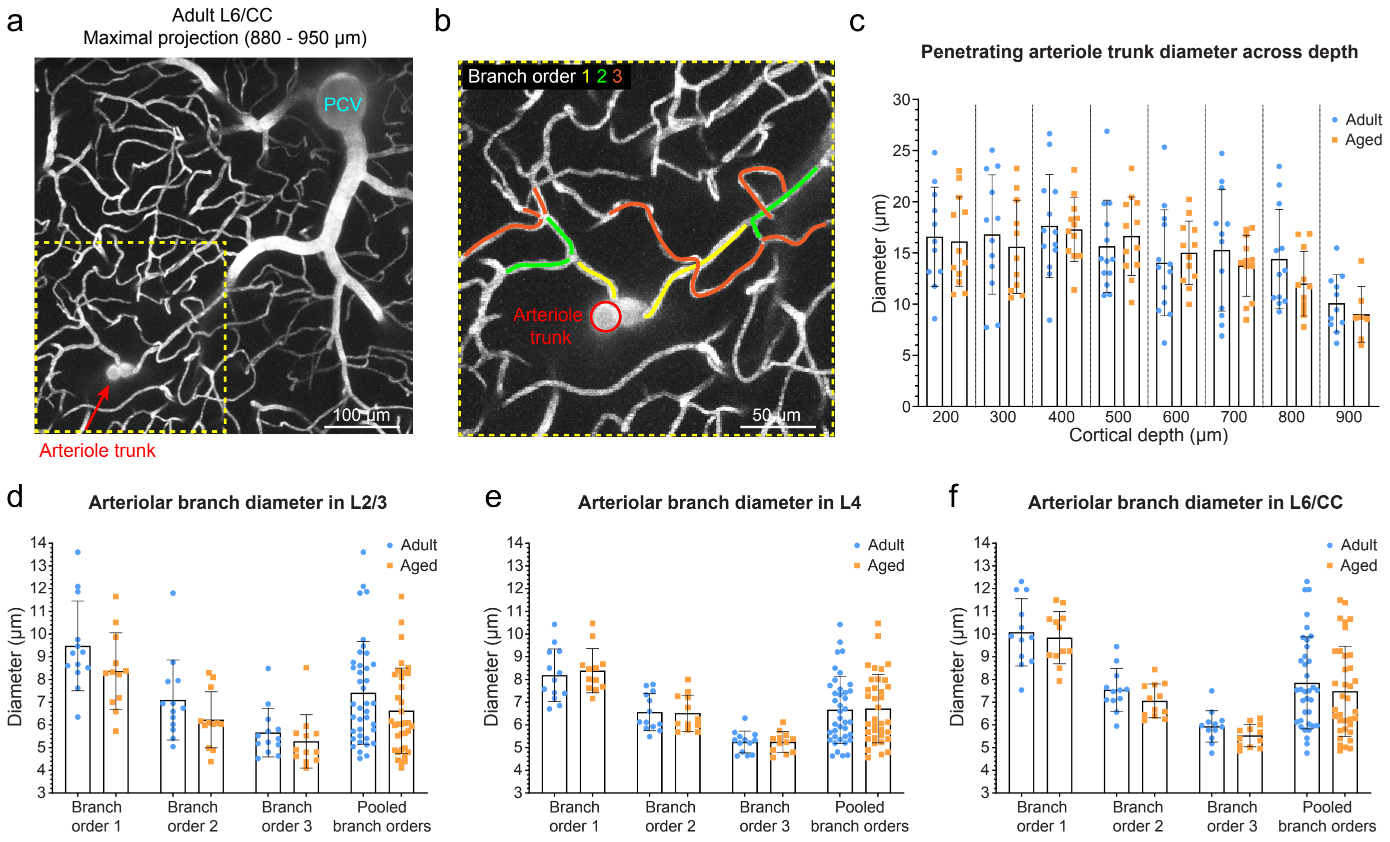


Extended Fig. 2 - Stamenkovic et al.

### Extended Figure 3

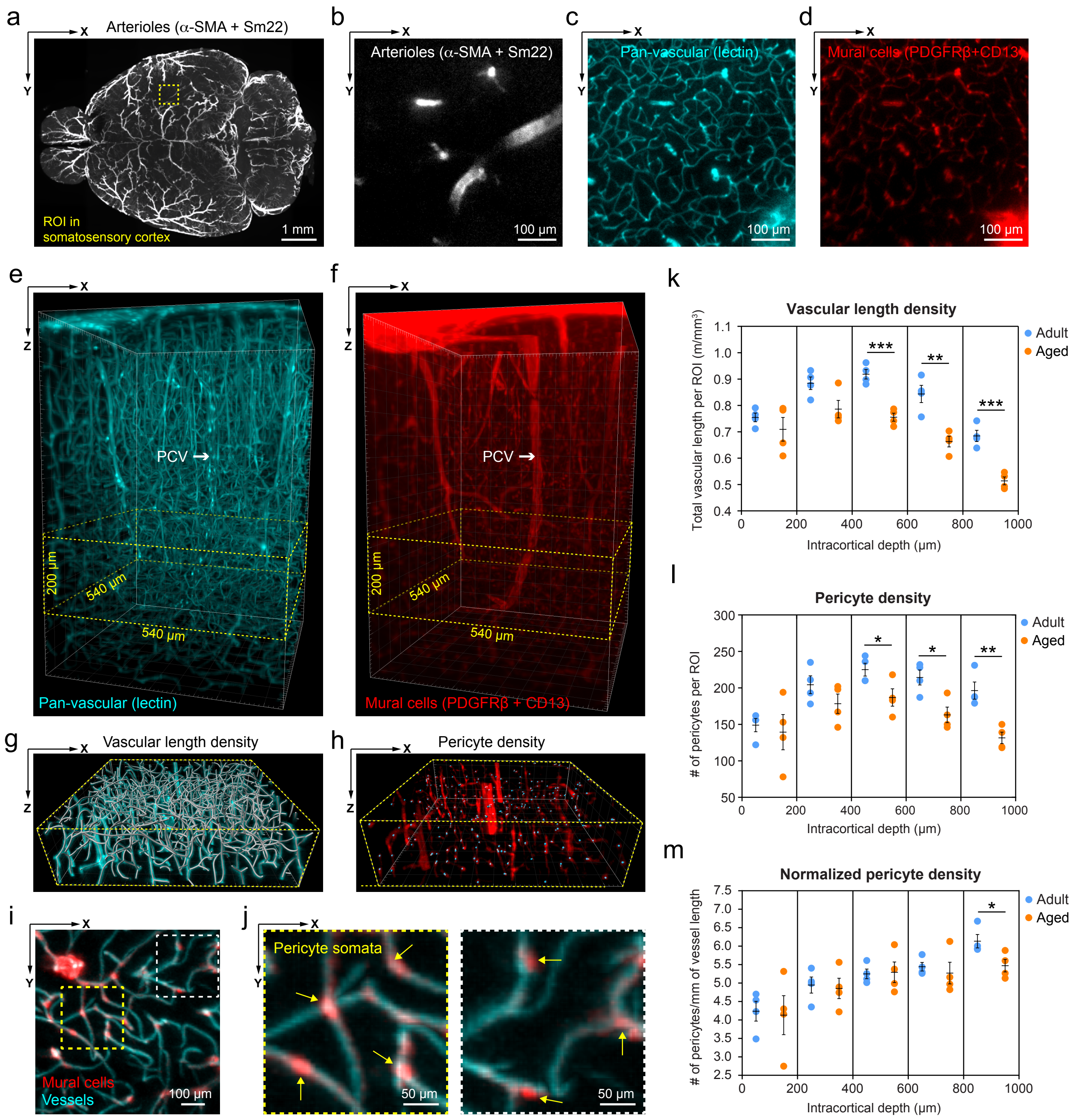


Extended Figure 3 - Stamenkovic et al.
